# Elucidation and Pharmacologic Targeting of Master Regulator Dependencies in Coexisting Diffuse Midline Glioma Subpopulations

**DOI:** 10.1101/2024.03.17.585370

**Authors:** Ester Calvo Fernández, Lorenzo Tomassoni, Xu Zhang, Junqiang Wang, Aleksandar Obradovic, Pasquale Laise, Aaron T. Griffin, Lukas Vlahos, Hanna E. Minns, Diana V. Morales, Christian Simmons, Matthew Gallitto, Hong-Jian Wei, Timothy J. Martins, Pamela S. Becker, John R. Crawford, Theophilos Tzaridis, Robert J. Wechsler-Reya, James Garvin, Robyn D. Gartrell, Luca Szalontay, Stergios Zacharoulis, Cheng-Chia Wu, Zhiguo Zhang, Andrea Califano, Jovana Pavisic

## Abstract

Diffuse Midline Gliomas (DMGs) are universally fatal, primarily pediatric malignancies affecting the midline structures of the central nervous system. Despite decades of clinical trials, treatment remains limited to palliative radiation therapy. A major challenge is the coexistence of molecularly distinct malignant cell states with potentially orthogonal drug sensitivities. To address this challenge, we leveraged established network-based methodologies to elucidate Master Regulator (MR) proteins representing mechanistic, non-oncogene dependencies of seven coexisting subpopulations identified by single-cell analysis—whose enrichment in essential genes was validated by pooled CRISPR/Cas9 screens. Perturbational profiles of 372 clinically relevant drugs helped identify those able to invert the activity of subpopulation-specific MRs for follow-up *in vivo* validation. While individual drugs predicted to target individual subpopulations—including avapritinib, larotrectinib, and ruxolitinib—produced only modest tumor growth reduction in orthotopic models, systemic co-administration induced significant survival extension, making this approach a valuable contribution to the rational design of combination therapy.

## Introduction

Diffuse midline gliomas (DMGs) represent aggressive, central nervous system (CNS) malignancies associated with dismal outcome, with <10% of patients surviving two years post-diagnosis^1^. They occur primarily in childhood, accounting for 20% of all pediatric CNS tumors, and represent the leading cause of pediatric brain tumor-related mortality^1^. They are spatially restricted to the midline structures of the brain (e.g., brainstem, thalamus, spine), thus making surgical resection highly challenging. While radiation therapy (RT) can provide temporary palliative control^2,3^, decades of research and clinical trials have yet to yield effective systemic therapies. In addition to presenting established driver mutations that are not directly actionable^4^, their heterogeneity and plasticity is proposed as a key determinant of adaptive resistance.

DMG-specific oncogenic drivers have been extensively characterized, including recurrent somatic histone mutations in H3.3 (*H3F3A*) and H3.1 (*HIST1H3B* and *HIST1H3C*) and overexpression of *EZHIP* in patients with wildtype (WT) H3, resulting in broad epigenetic dysregulation^5–9^. Identification of additional actionable oncogenes has had limited success, as DMG harbors few such recurrent alterations and resistance has consistently emerged upon their pharmacologic targeting^10–12^. Due to their intrinsic heterogeneity^7,9,12–14^, these failures are consistent with selection of epigenetically distinct, drug-resistant subpopulations, highlighting the urgent need for combination approaches^12,15,16^.

To address these challenges, we leveraged established network-based methodologies to systematically elucidate Master Regulator (MR) proteins that, as previously reported^17^, represent mechanistic determinants of cancer cell states. Critically, we have shown that, despite being rarely mutated, MR proteins are highly enriched in both tumor state essential genes and in genes eliciting synthetic lethality, thus providing potentially actionable non-oncogene dependencies^17–20^. Indeed, interrogation of large-scale drug perturbation profiles—characterizing the transcriptional response of tumor cells to clinically relevant drugs—has been effective in identifying drugs capable of abrogating tumor growth *in vivo* by either targeting a single essential MR (OncoTarget^21^) or by inverting the activity of an entire MR module (OncoTreat^22^). Both methods have been approved by the NY and CA Dept. of Health as CLIA compliant and are extensively validated, both preclinically^23,24^ and clinically^25^. Specifically, they have shown remarkable disease control rates (64% and 91%, respectively) in low-passage, patient derived xenograft (PDX) models from treatment-refractory tumors ^25^.

In this manuscript, we apply these methodologies to the identification and pharmacological targeting of MRs representing critical non-oncogene dependencies of molecularly distinct DMG subpopulations, see **Fig. 1** for a schematic representation of the study design. Specifically, we first assessed whether analysis of bulk RNA-seq profiles, from three publicly available cohorts, may identify MRs and associated drugs for cell states representing the majority subpopulation, thus providing potentially novel insight into clinical outcome stratification and treatment. Then, to elucidate the role of DMG heterogeneity in modulating drug sensitivity, we analyzed single-cell profiles from six DMG patients^26^. Indeed, while previous studies have helped elucidate a hierarchy of transcriptional DMG states representing co-existing subpopulations^26,27^ associated with astrocytic (AC-like), oligodendrocytic (OC-like) and oligodendrocytic precursor (OPC-like) differentiation, their mechanism-based dependencies and potential pharmacologic targeting remain elusive and represent the motivation for this study.

**Figure 1.**
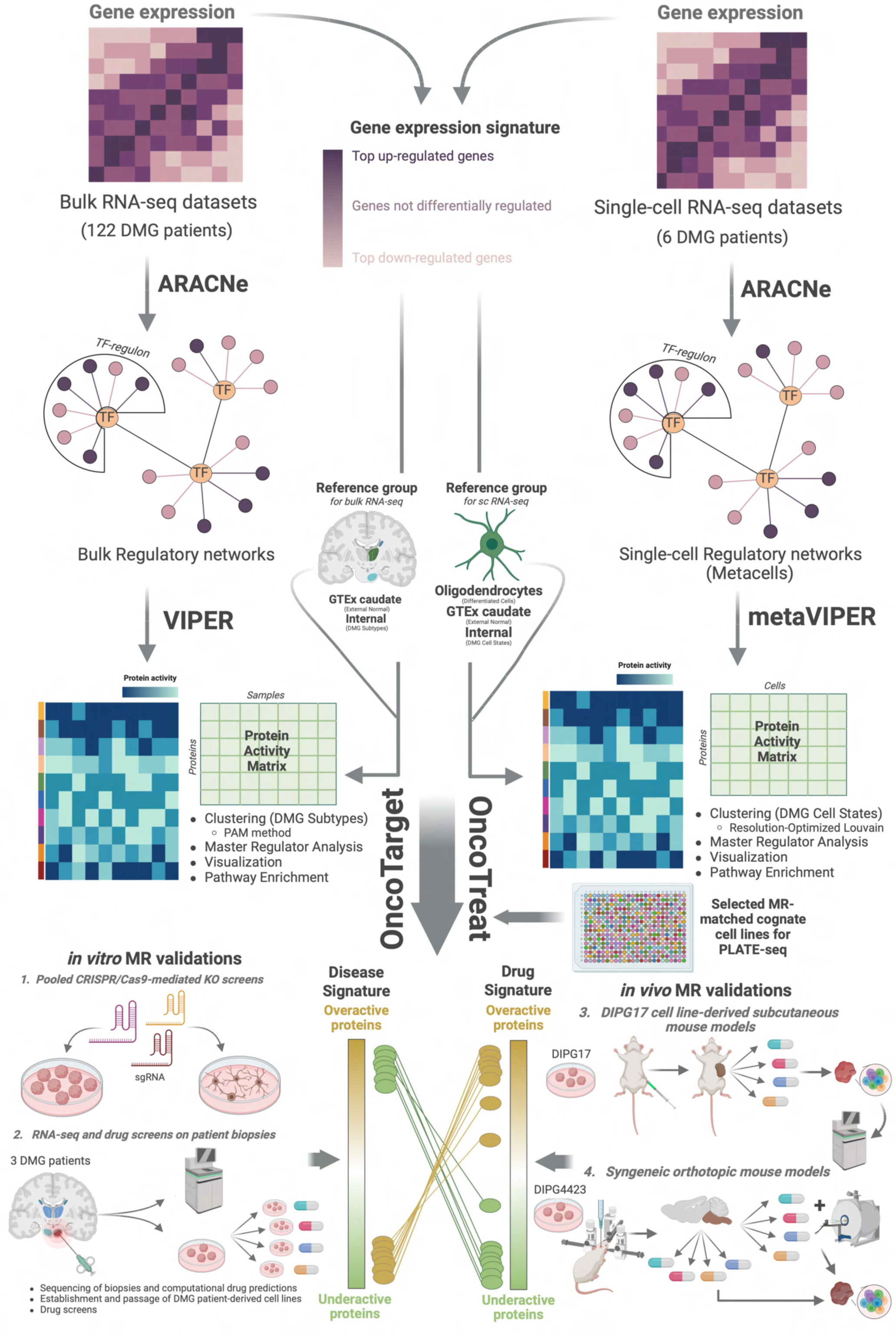
Schematic overview of the study design. Gene expression signatures across 122 bulk and 6 single-cell RNA-seq samples from patients with DMG were transformed by VIPER into protein activity signatures based on enrichment of transcriptional targets in the context of an ARACNe-generated DMG tumor bulk and single-cell gene regulatory network, respectively. Bulk protein activity analysis was used to define disease subtypes, and master regulators (MR) of tumorigenesis whose essentiality was validated in a pooled CRISPR/Cas9-mediated knock-out screen in DMG cell lines (1). While bulk analysis focuses on predominant tumor populations, single-cell protein activity analysis was used to define tumor subpopulations co-existing in patients that may have orthogonal molecular drivers and drug sensitivities which would be averaged out in bulk data. We then leveraged two NYS/CA CLIA-certified clinical tests, OncoTarget and OncoTreat, to identify clinically relevant drugs, for each bulk sample or single-cell subpopulation, that can directly inhibit individual tumor MRs or reverse the activity of a repertoire of the most dysregulated MRs, respectively. For the latter, we generated drug perturbed RNA-seq profiles for 372 FDA-approved and late-stage experimental drugs in two cognate DMG cell lines by PLATE-seq to define drug-induced differential protein activity signatures. Bulk-based drug predictions were validated via functional drug screens in primary tumor-derived cells from patient biopsies (2). Single-cell drug predictions prioritized drugs targeting distinct tumor compartments whose subpopulation-specific effects were validated in drug-perturbed scRNA-seq analysis of DMG cell line-derived subcutaneous mouse models (3). Effects on tumor volume and survival were then evaluated for individual drugs and drug combinations targeting complementary subpopulations in a syngeneic orthotopic DMG mouse model (4).

Protein activity-based analysis, using the VIPER algorithm^28^, identified candidate MRs as mechanistic determinants of seven transcriptionally-distinct DMG states. These were used to prioritize state-specific MR-targeting drugs, using a single cell implementation of the OncoTarget^21^ and OncoTreat^22^ algorithms. For the latter, we generated large-scale perturbation profiles representing the response of two DMG cell lines, chosen to recapitulate state-specific MRs, to treatment with a library of clinically relevant oncology drugs.

Single-cell RNA-seq (scRNA-seq) profile analysis of drug and vehicle-treated DMG cell line-derived subcutaneous mouse models confirmed the state-specific activity for eight of the nine predicted drugs. Specifically, it confirmed avapritinib, dinaciclib, trametinib, mocetinostat, and etoposide as statistically significant inhibitors of OPC-like cells, as well as ruxolitinib, larotrectinib, and venetoclax as inhibitors of more elusive, drug-resistant AC-like cells, with larotrectinib also inhibiting OC-like cells. Combination therapies targeting distinct cell states significantly outperformed monotherapy in an orthotopic syngeneic pontine DMG mouse model that faithfully recapitulated the heterogeneity of human DMGs. Most notably, the combination of avapritinib and either larotrectinib or ruxolitinib induced significant survival extension compared to both vehicle control and to the best monotherapy. Similar results were observed for other combinations. Thus, predicted drugs not only specifically ablated their predicted target subpopulations, but drug combinations targeting complementary states significantly improved outcome compared to their corresponding monotherapies.

Since MR proteins are identified based on the differential expression of genes enriched in their *bona fide* physical targets^18,29,30^, this study introduces a mechanism-based approach to prioritize combination therapy at the single cell level. Tested drugs were restricted to those with immediate potential for clinical translation, including in terms of blood brain barrier (BBB) permeability. As such, these results may help develop novel clinical trials for children with this universally fatal disease. More important, the approach provides a novel, highly generalizable paradigm for the prediction of combination therapies to target highly heterogeneous tumors that undergo treatment adaptation based on rapid selection of epigenetically distinct, drug-resistant subpopulations.

## Results

### Bulk protein activity inference by DMG regulatory network analysis

We have extensively shown that—akin to a multiplexed gene reporter assay—the activity of regulatory proteins, including transcription factors (TFs), co-factors (co-TFs), and signaling proteins (SPs), can be measured by the differential expression of their regulatory targets (regulon)^28^, as implemented by the VIPER algorithm^28^. Regulons are highly context-specific and can be inferred by ARACNe3^31^, the latest incarnation of ARACNe-AP^30^, an algorithm that leverages large scale gene expression data to infer protein-target interactions with >70% accuracy, based on experimental validation^29,30^. To reverse engineer a DMG-specific gene regulatory network (interactome), we used ARACNe3 to analyze 120 bulk RNA-seq profiles from three publicly available DMG cohorts—including St. Jude^9,32^, PNOC^33^ and CBTTC^34^ (Extended Data **Table 1**). DMG samples were identified as high-grade glial tumors occurring in a midline structure of the brain. The resulting interactome comprised regulons with ≥ 50 targets for 6,138 proteins with 7,002,347 interactions (median regulon size of 1,097 targets per regulatory protein).

We then used VIPER to assess the activity of the 6,138 regulatory proteins in each DMG sample based on the enrichment of their regulons in genes differentially expressed in the sample vs. the average of two control populations. First, to optimally distinguish between molecularly distinct DMG subtypes, we compared each sample to the average of all the samples in the cohort (*internal signature*). Second to assess the mechanistic contribution of each protein to DMG oncogenesis— i.e., as represented by the “normal → tumor” cell state transition—we compared each sample to the centroid of 246 normal caudate tissue samples from GTEx^35^ (*caudate signature*). As previously shown^19,36^, VIPER analysis is especially effective in removing (a) technical batch effects that are not consistent with the regulatory architecture of the network and (b) sample-specific biological bias, e.g., associated with many differentially expressed genes in patient specific amplicons that do not contribute to the activity of MR proteins controlling tumor cell state. Indeed, while gene expression-based PCA analysis stratified patients into three clusters, each comprising samples from the same cohort, protein activity-based analysis virtually eliminated this effect, without requiring normalization or batch effect correction, thus reducing the risk of potential artifacts (Extended Data **Fig. 1a**).

### Activity-based analysis identifies DMG subtypes associated with survival

Unsupervised DMG sample clustering of VIPER-inferred protein activity (*internal signature*), using partition around mediods (PAM)^37^, identified an optimal two-cluster solution, based on silhouette score (SS) analysis, including cluster C1 (average SS = 0.32), with 62 samples, and cluster C2 (average SS = 0.30), with 58 samples (**Fig. 2a**, Extended Data **Fig. 1b,c**). Clusters were significantly associated with >1-year survival (p = 0.038, by Fisher’s exact test (FET)), with 51.8% vs. 26.0% of evaluable subjects in C1 vs. C2 surviving to one year, respectively (log-rank p = 0.026) (**Fig. 2b**). Equally important, clusters were not associated with potential confounding features, including dataset provenance (p = 0.74), sample type (autopsy vs. biopsy; p = 1), tumor type (initial CNS, progressive, or recurrent disease; p = 0.34), tumor location (pons, thalamus, other midline; p = 0.39), or molecular features such as TP53 mutation status (p = 0.28) or histone mutation status (p = 0.78). Notably, gene expression-based cluster analysis presented much weaker average SS (0.093)—i.e., below the 0.25 threshold generally used as a standard for good cluster tightness—and failed to stratify survival (log-rank p = 0.17) (Extended Data **Fig. 1c,d**).

**Figure 2.**
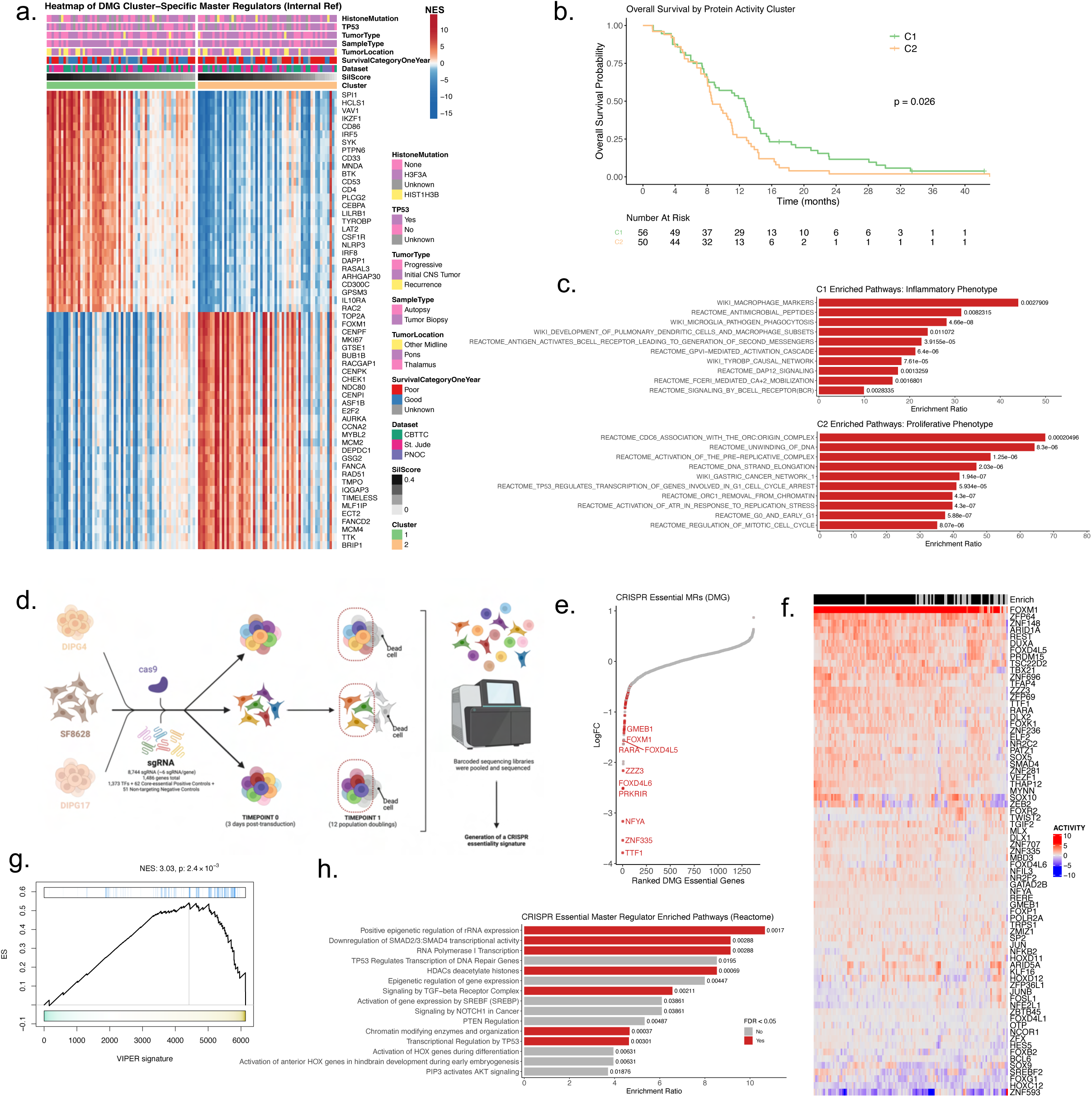
Bulk protein activity inference by DMG regulatory network analysis identifies subtypes associated with survival and master regulators (MRs) of tumorigenesis validated in CRISPR/Cas-9 KO screens. **a.** Unsupervised clustering of sample-specific, VIPER-inferred differential protein activity identifies two clusters, C1 (green) and C2 (orange). Heatmap depicts the VIPER normalized enrichment scores (NES) across the samples for the top and bottom 25 most dysregulated MRs in each cluster. Relevant sample clinical and molecular features are depicted in the annotation bars. **b.** DMG patient overall survival by protein activity-based cluster for 106 patients with outcome information. P-value computed by log-rank test. **c.** Pathway enrichment analysis for the top 50 cluster-specific MRs identify an inflammatory vs. proliferative phenotype. Pathways are ranked by the enrichment score using an over-representation analysis and the associated FDR-corrected p-value is displayed. **d.** Schematic overview of the pooled CRISPR/Cas9 knock-out experiment in three patient-derived DMG cell lines used to validate the essentiality of VIPER-inferred DMG MRs. **e.** Scatter plot showing the CRISPR DMG gene essentiality signature integrated across the three cell lines with genes ranked by their log2FC with the most essential genes on the left. VIPER-inferred DMG candidate MRs are shown in red and are enriched among the most essential genes. **f.** Heatmap of VIPER-inferred differential protein activity (*caudate* signature) for 73 DMG CRISPR essential MRs belonging to the leading edge of at least one DMG sample from this analysis. The enrich annotation bar reflects presence (black) or absence (gray) of statistical significance (p ≤ 0.05, one-tailed GSEA) for enrichment of DMG essential genes in the sample’s differential protein activity signature. **g.** Results of GSEA analysis demonstrating significant enrichment of DMG essential genes (blue ticks) in the ranked Stouffer-integrated protein activity signature of the 94 samples showing statistically significant enrichment from f. **h.** Pathway enrichment analysis for the 73 DMG CRISPR essential MRs with pathways ranked by enrichment score. Statistically significant pathways after FDR correction are in red.

To nominate MR proteins for the two DMG subtypes, we used Stouffer’s method to integrate the Normalized Enrichment Score (NES), representing the VIPER-inferred activity of each protein, across the samples in each of the two clusters, weighted by their SS. Pathway enrichment analysis, for the top 50 most differentially active proteins in each cluster (i.e., candidate MRs), showed enrichment of innate and adaptive immune pathways vs. cell cycle and DNA damage response pathways in C1 vs. C2 (**Fig. 2c**), leading to their characterization as inflammatory and proliferative subtypes, respectively. We identified statistically significant enrichment of WikiPathways “macrophage markers” (p = 0.003) and “microglia pathogen phagocytosis” (p = 4.66 x 10^-8^) in C1 MRs, suggesting that tumors presenting greater myeloid cell infiltration have improved survival.

### CRISPR/Cas9-mediated validation of candidate MR proteins

We proceeded to functionally validate the essentiality of candidate MRs—as nominated by VIPER using the caudate control signature (i.e., MRs involved in DMG tumorigenesis). For this purpose, we first generated a consensus DMG activity signature by integrating the NES of each protein across all DMG samples, using Stouffer’s method. This showed consistent, aberrant activity (p < 10^-5^) of a common repertoire of candidate MRs, previously associated with aggressiveness and proliferation (*e.g.,* FOXM1, CENPF, TOP2A, etc.^38–40)^ (Extended Data **Fig. 2a**). We then performed pooled CRISPR/Cas9-mediated KO screens in three human DMG cell lines—SF8628 (H3F3AK27M), DIPG4 (HIST1H3BK27M, TP53^Mut^, ACVR1^Mut^), and DIPG17 (H3F3AK27M, TP53^Mut^)—using guide RNAs (sgRNAs) targeting 1,433 genes (mean of 6 sgRNA per gene) (**Fig. 2d**, Extended Data **Table 2**). These include all TFs, as well as selected core essential and non-targeting genes as positive and negative controls, respectively (Extended Data **Fig. 2b**). Guide depletion at a late (12 population doublings) vs. early (3 days) time point was assessed using the MAGeCK^41^ algorithm, yielding both cell line specific and integrated DMG essentiality signatures.

The most aberrantly active proteins in the consensus DMG activity signature were represented among the top DMG essential genes when integrated across all three cell lines (**Fig. 2e**). On an individual sample basis, the vast majority of DMG samples (94 of 122, 77%) presented significant enrichment of DMG essential genes (negative log2FC and FDR ≤ 0.05) in aberrantly active proteins (p ≤ 0.05, one-tailed GSEA) (**Fig. 2f**). Significantly enriched samples were more likely to belong to C2 (p = 0.001, by FET), to be biopsy vs. autopsy derived (p = 0.04), and non-pontine vs. pontine tissue derived (p = 0.004). When integrated across all three cell lines, DMG essential genes were significantly enriched in genes encoding for aberrantly active protein in the consensus activity signature across these 94 samples (p = 2.4×10^-3^) (**Fig. 2g**). Taken together, the analysis identified a set of 73 CRISPR essential MRs—i.e., genes in the leading edge of at least one DMG sample—with aberrant, integrated VIPER activity across all samples (**Fig. 2f**).

When assessed on an individual cell line basis, essential genes were also highly enriched in aberrantly active proteins (Extended Data **Fig. 2c,d**). Moreover, for two of the three cell lines, DIPG4 and SF8628, the optimal result was achieved when assessing enrichment of essential genes in genes encoding for proteins aberrantly active in that same cell line (p_DIPG4_ = 5.6×10^-6^ and p_SF8628_ = 2.0×10^-5^) across six DMG cell lines for which RNA-seq profiles were generated, including SF8628, SF7761, SU-DIPG-IV (DIPG4), SU-DIPG-VI (DIPG6), SU-DIPG-XVII (DIPG17), and SU-DIPG-XXXVI (DIPG36). Both results represent a two-order of magnitude improvement over the integrated analysis, thus confirming VIPER’s ability to nominate sample-specific MRs. The only exception was DIPG17, which produced the 4^th^ best enrichment against its own differentially active proteins (one-tailed GSEA p = 6.4×10^-3^) (Extended Data **Fig. 2e**).

Among the top CRISPR-essential MRs, Forkhead box M1 (FOXM1) emerged as the most conserved across DMG samples, suggesting it may play a critical role in regulating DMG cell state and nominating it as a candidate therapeutic target^42^ (**Fig. 2f**). Pathway enrichment analysis of the 73 DMG essential MRs in the Reactome database (**Fig. 2h**) highlighted activation of FOXM1-related pathways, including “PTEN regulation” (p = 0.005) and “PIP3 activates AKT signaling” (p = 0.019). Other pathways highly enriched in essential MRs including “transcriptional regulation by TP53” (p = 0.003), “HDACs deacetylate histones” (p = 0.0007), “chromatin modifying enzymes and organization” (p = 0.0004), and “activation of HOX genes during differentiation” (p = 0.0063), reflect established molecular, epigenetic, and developmental dysregulation mechanisms in DMG. Finally, 14 CRISPR-essential candidate MRs were annotated as H3K27me3 targets in the cerebellum (pontine data not available) by the Encode Project^43,44^, including DLX1, ZZZ3, GMEB1, DLX2, TGIF2, TWIST2, GRHL3, NR2F2, SOX10, ELK3, NR5A1, HOXB4, SOX9, and HOXD9, suggesting that epigenetic de-repression associated with H3K27M mutations may mechanistically account for their aberrant activation. Taken together, these data show that VIPER-inferred MRs recapitulate DMG essential proteins and are consistent with DMG genetics, thus providing a wealth of novel potential targets for pharmacologic intervention and a critical signature for identification of MR-inverter drugs using the OncoTarget and OncoTreat algorithms.

### Elucidating the DMG-specific mechanism of action (MoA) of clinically relevant drugs

Given the rich signature of essential MRs identified in the previous section and the high likelihood that many of the non-essential MRs may elicit synthetic lethality, as shown in previous studies^18,20^, we focused on the identification of candidate MR-inverter drugs—i.e., drugs that simultaneously activate and inactivate the most aberrantly inactive and active DMG tumor proteins, respectively— which we have shown to abrogate tumor viability *in vivo*^22,25^. For this purpose, we generated perturbational profiles characterizing the transcriptional response of two high-fidelity DMG cell lines to 372 clinically relevant drugs, including FDA-approved and late-stage experimental oncology drugs (Extended Data **Table 3**). Among the six available DMG cell lines, SF8628 (H3F3AK27M^Mut^) and DIPG6 (H3F3AK27M^Mut^, TP53^Mut^) were selected as optimal models to report on MR-specific drug activity, based on their ability to recapitulate the MRs of complementary DMG sample subsets for a total of 119/122 (97.5%) DMG samples (p < 10^-5^ OncoMatch Analysis^45^) (Extended Data **Fig. 3c**). The OncoMatch algorithm—designed to assesses MR conservation between a model (e.g., cell line, PDX, GEMM, etc.) and a tumor sample—has proven highly effective in predicting *in vivo* drug mechanism of action based on *in vitro* (i.e., cell line-based) perturbation assays^25,36^.

RNA-seq profiles were then generated at 24 hours following treatment of each cell line with the 372 drugs—titrated at the lowest of their highest sublethal concentration (48h EC_20_) or max tolerated serum concentration from the literature—as well as DMSO as vehicle control, using the PLATE-Seq microfluidic automation platform^46^. For each drug, the VIPER-inferred differential activity of 6,138 regulatory proteins in drug vs. DMSO-treated cells was taken to represent its DMG-specific MoA, as supported by numerous publications^22,25,47^. **Fig. 3a,b** highlights the hierarchical clustering of VIPER assessed drug MoA in each cell line for the subset of drugs demonstrating the strongest effect on the cells’ transcriptional signature. As shown, and consistent with expectations and prior publications, drugs with related high-affinity targets were often found in the same cluster. However, drugs with different high-affinity targets may also induce highly similar MoA profiles, due to the cell context-specific canalization of their effects.

**Figure 3.**
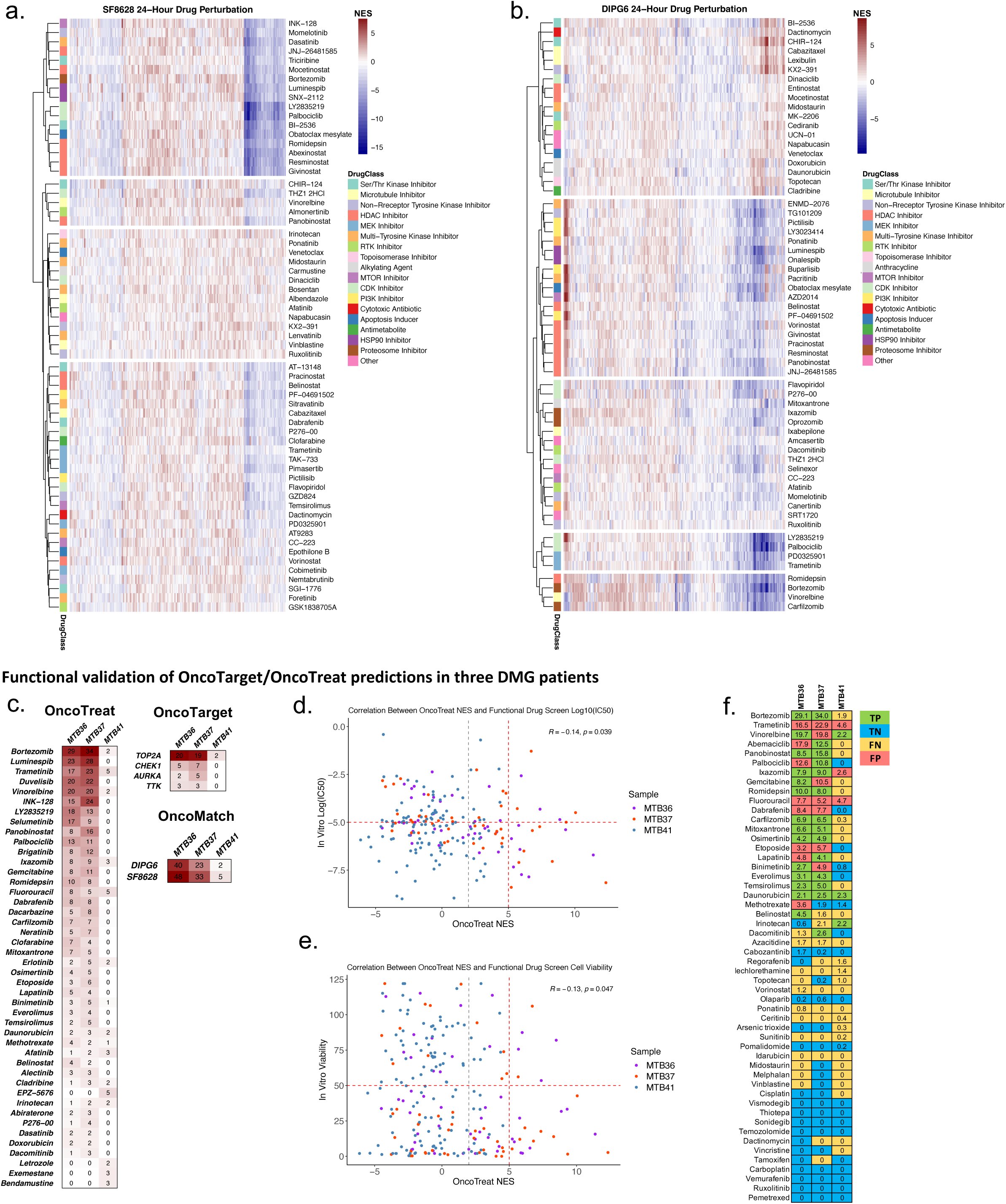
DMG-specific Mechanism of Action (MoA) of oncology drugs and functional validation of bulk OncoTarget/OncoTreat drug predictions in primary tumor cells from patient biopsies. **a-b.** Heatmap for the 24-hour PLATE-seq drug perturbations in the SF8628 (a) and DIPG6 (b) DMG cell lines reflecting the differential protein activity for a subset of drugs demonstrating the strongest effects on the cells’ transcriptional signature. Drugs are annotated by their canonical mechanism. The most dysregulated VIPER-inferred proteins are shown in columns and unsupervised hierarchical clustering highlights drugs that induce similar transcriptional responses in the DMG context. **c.** OncoTarget and OncoTreat drug predictions from bulk RNA-seq generated from three DMG patient tumors. Patients are in columns. Values represent -log10(Bonferroni p-values) for enrichment of patient MRs (*caudate signature*) in the drug-induced differential protein activity signature (OncoTreat), for the activity of actionable proteins (OncoTarget), and for the enrichment of patient MRs in the cell line models’ differential protein activity signature (OncoMatch). Functional drug screens were performed in primary tumor cells from these three patients and OncoTarget/OncoTreat-predicted score and *in vitro* growth abrogation could be compared in 223 instances. Scatter plots demonstrate association of OncoTreat scores with **d.** *in vitro* log10(IC50) and **e.** cell viability in the context of *in vitro* drug screening, with statistically significant correlation identified for both by Spearman’s correlation. **f.** Comparison of OncoTarget/OncoTreat predictions (Bonferroni p ≤ 0.01) with drug efficacy based on functional screens (IC50 ≤ 10uM and cell viability ≤ 50%) in primary tumor cells from 3 patient biopsies with colors reflecting true positives (green), true negatives (blue), false negatives (yellow) and false positive (red). OncoTarget/OncoTreat achieved 80% specificity and 65% positive predictive value.

OncoTreat was then used to rank candidate MR-inverter drugs across the 122 bulk DMG samples based on the statistical significance of the enrichment of the top 50 most active and inactive MRs in each DMG sample (*caudate signature*) in proteins differentially inactivated or activated in drug vs. vehicle control-treated cells. In addition, when available, we also considered high-affinity inhibitors of the most statistically significant MRs (p ≤ 10^-5^) (OncoTarget algorithm). The analysis stratified predicted drug sensitivity into four pharmacotypes (Extended Data **Fig. 3a,b**), representing sample subsets predicted to be sensitive to the same drugs; among others, this identified samples with predicted sensitivity to HDAC, MEK, CDK2/4/6, EZH2, and TOP2A inhibitors. Pharmacotypes showed highly significant co-segregation with the proliferative versus inflammatory DMG subtypes (FET p = 1.64×10^-6^). Notably, confirming the effectiveness of the analysis, many of the inferred MR-inverter drugs had been reported among the most active drugs by multiple DMG high-throughput drug screens^12,48,49^. The proposed analysis, however, also provides critical MR-based stratification of candidate responders vs. non-responders, which is not possible when drug screening assays are used.

### Functional validation of OncoTarget/OncoTreat drug predictions

To demonstrate the accuracy and specificity of bulk profile-based OncoTarget/OncoTreat drug predictions, we analyzed data from three DMG patients (MTB36, MTB37, and MTB41) whose tumor tissue had been subjected to bulk RNA-seq as well as *in vitro* drug screening as part of a functional precision medicine program at Rady Children’s Hospital^50^. The MTB36 and MTB37 screens comprised a library of 63 drugs, of which 52 were also included in the PLATE-Seq panel, while the MTB41 screen comprised 175 drugs, of which 124 were also included in the PLATE-Seq panel. Data included dose response curves to assess each drug’s IC_50_ and cell viability post treatment measured by CellTiter Glo. For each sample, we generated VIPER-inferred differential protein activity (*caudate signature*) and predicted MR-reversing drugs by OncoTarget/OncoTreat (**Fig. 3c**). Only drugs that affected viability at a concentration lower than 10uM were considered in this comparison, since higher concentrations were not profiled by PLATE-Seq.

Among 228 drugs tested by both OncoTarget/OncoTreat and the functional drug screen, OncoTarget/OncoTreat predictions (p ≤ 0.01) were twice as likely to demonstrate *in vitro* efficacy (IC_50_ ≤ 10uM and viability ≤ 50%) in the respective patient, compared to non-predicted drugs (odds ratio OR = 2.28, 95% CI 1.21 – 4.41, two-tailed p = 0.0076). OncoTarget/OncoTreat predictions demonstrated high specificity—91 of the 113 drugs (80.5%) that failed to show *in vitro* effects were correctly predicted by the algorithms—and positive predictive value—41 of the 63 drugs (65.1%) predicted by the algorithms indeed demonstrated *in vitro* effect (**Fig. 3f**). Overall accuracy was 132/228 (57.9%) with statistically significant Spearman’s correlation seen between the OncoTreat score and *in vitro* IC_50_ (p = 0.039) and cell viability (p = 0.047) (**Fig. 3d,e**). This demonstrates the ability to leverage these algorithms to predict drugs specifically likely to elicit response in individual DMG patients, based only on bulk profile analyses, thus providing a mechanism-based rationale for the design of clinical trials, as well as to inform selection of candidate drugs for DMG patients who lack standard-of-care options after initial RT.

### Single cell analysis reveals transcriptionally distinct cell states co-existing in virtually all DMG patients

A critical challenge of bulk analyses is that they fail to account for tissue heterogeneity, including stromal infiltration and presence of genetically or epigenetically distinct yet coexisting malignant cell states, which have been shown to exist in DMG ^51^. This can affect bulk-based drug predictions because the average MR signature generated by such heterogeneous tissues may not recapitulate any of the individual cell states they comprise, thus resulting in severe artifacts. In addition, smaller subpopulations that fail to respond to drugs predicted by bulk analyses may rapidly lead to the emergence of drug resistant tumors.

We thus sought to perform OncoTarget/OncoTreat drug predictions at the level of each transcriptionally distinct subpopulation that comprise a DMG tumor for potential development of combination therapy approaches. For this purpose, we analyzed 4,860 high-quality single cells from published scRNA-seq profiles from six DMG patient biopsies^51^ using the single cell versions of the ARACNe-AP and VIPER algorithms^24^. Using scRNA-seq from previously defined putative tumor cells^68^ and ARACNe-AP, we generated a DMG-specific single-cell interactome for each patient, which comprised regulons with ≥ 50 targets for a median of 1,743 regulatory proteins (714 to 2469) and 87,125 transcriptional interactions (35,700 to 123,450). We then transformed single-cell gene expression signatures computed relative to the centroid of all the cells into protein activity profiles by metaVIPER^22,24^—the extension of VIPER to single cell datasets.

Unsupervised, protein activity-based cluster analysis identified nine clusters, four of which comprised non-malignant cells, as assessed from high activity of CD33 and of a microglia gene signature^52^ (Extended Data **Fig. 4a,b**). Expression of the microglia gene signature and CD33 was not detected in these populations, preventing their cell type characterization with gene expression analysis alone. InferCNV analysis^53^ compared to normal myeloid cells, confirmed the malignant nature of cells in the remaining five clusters (n = 3,039) (Extended Data **Fig. 4c,d**).

We then refined the metaVIPER analysis to assess the activity of 2,821 regulatory proteins relative to the centroid of the malignant cells in the remaining five clusters. The analysis identified seven transcriptionally distinct cell states and associated candidate MR signatures that were detected in virtually every sample (**Fig. 4a,b**, Extended Data **Fig. 4e,f**). Among the most aberrantly active proteins in each cluster, we identified distinct yet established glial lineage markers, allowing further characterization of each state’s biology (**Fig. 4c,d**). Specifically, three states were characterized as OPC-like, based on the activity of markers such as OLIG1, OLIG2, and PDGFRA; these were further divided into a more quiescent OPCQ state (OPC-like Quiescent), a more proliferative OPC state (OPC-like), and a highly proliferative OPCC state (OPC-like Cycling), as determined by the relative activity of proliferative markers, such as TOP2A, CENPF, and TYMS. Two additional states were characterized as OC-like, based on the activity of differentiated OC markers, such as SIRT2, PLP1, and MPZL1, including an OPC/OC state (OPC/OC-like) with partial activation of OPC markers and a second OC state (OC-like) with no OPC marker activity. One additional AC state (AC-like) was characterized by high activity of astrocyte differentiation markers, such as SOX9 and APOE, while a final OPC/AC state (OPC/AC-like) presented a transitional signature with co-activation of both OPC and AC markers. For simplicity, given their protein activity similarity, we will refer to OPC states (i.e., OPC, OPCQ, and OPCC), OC states (i.e., OC and OPC/OC) and AC states (i.e., AC and OPC/AC).

**Figure 4.**
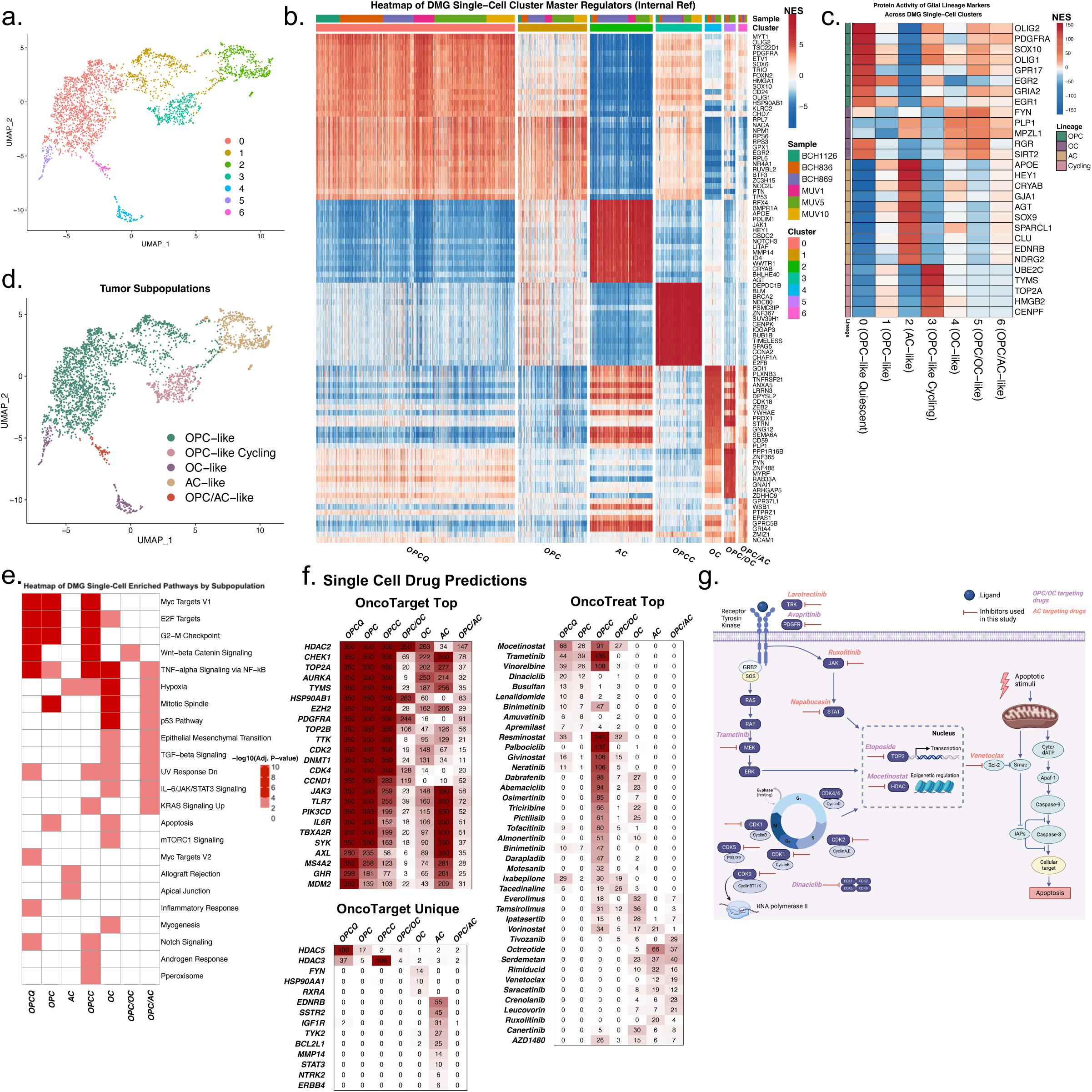
Single cell protein activity analysis reveals seven distinct DMG cell states with orthogonal MR signatures and predicted drug sensitivities. **a.** UMAP projection of protein activity for 3,039 DMG tumor single cells from 6 DMG patients. Resolution-optimized Louvain clustering by VIPER-inferred protein activity identified 7 clusters. **b.** Heatmap of VIPER-inferred protein activity across the single cells in columns for the top 15 most activated MRs in each of the 7 clusters annotated across the bar on top. Each of the 6 DMG patients had cells belonging to each of the 7 clusters as reflected in the Sample annotation bar. **c.** Heatmap of Stouffer-integrated protein activity across cells in each of the 7 clusters for known oligodendrocyte precursor cell (OPC), oligodendrocyte (OC), and astrocyte (AC) lineage markers allowing annotation of clusters (see column names) by representative glial lineage and degree of proliferation based on cycling marker activity. **d.** UMAP projection of the DMG tumor single cells colored by cell annotation used for comparing DMG tumor model single cells to patient tumor populations in future sections. **e.** Heatmap showing -log10(FDR-adjusted p-values) for pathways significantly enriched in at least one cluster using the MSigDB Hallmark 2020 database and each population’s significantly differentially expressed genes. **f.** Single cell OncoTarget and OncoTreat scores, represented as -log10(Bonferroni p-value), for each of the 7 populations. Activity of the top 20 actionable proteins assessed by OncoTarget and top population MR-reversing drugs predicted by OncoTreat for each of the 7 populations is represented in the OncoTarget and OncoTreat Top heatmaps, while activity of actionable proteins uniquely activated in each cluster is depicted in OncoTarget Unique. **g.** Schematic of known mechanisms for the five top drugs selected to uniquely target the OPC- and OC-like cell states (purple) and four drugs selected to uniquely target the AC-like cell states (pink) that were validated *in vivo*. Figure adapted from ^66,67^ and generated with BioRender.

Further enrichment analysis, using genes identified as uniquely differentially expressed in each state (Extended Data **Table 4**), identified established signaling and regulatory pathways related to glial lineage development, thus providing additional insight into potentially targetable state-specific mechanisms (**Fig. 4e**). For example, several pathways regulating the balance of proliferation and OPC-to-OC differentiation were dysregulated in OPC cells, including “MYC targets”^54^, “Wnt-beta Catenin signaling”, “TNF-alpha Signaling via NF-kB”, and “Notch Signaling^55,56,57,58^. Further, “mTORC1 Signaling”, promoting OC differentiation and transition into the myelinating phase, was uniquely activated in the OC cluster ^59,60^. The transitional OPC/AC cluster demonstrated unique activation of “IL6/JAK/STAT3 Signaling”, promoting angiogenesis ^61^, reactive astrogliosis ^62,63^, and inhibition of OPC-to-OC differentiation^64^. Pathways critical to cell cycle regulation—“E2F targets” and “G2-M Checkpoint”—were activated in OPC states, confirming their proliferative nature^65^. ^66,67^

Despite some cross-sample differences, OPC states consistently comprised the majority of cells in each patient (Extended Data **Fig. 4f**). Not surprisingly, MRs associated with these states—e.g., E2F8, CHEK1, MYBL1, CENPK, FOXM1, TOP2A, etc.—overlapped with those identified by bulk-level analyses and validated by CRISPR-mediated KO. In contrast, bulk-level analyses failed to identify any MRs representing mechanistic determinants of minority cell states, including OC- and AC-related ones.

### Predicting drugs targeting cell state-specific MR proteins

The presence of molecularly distinct states that were not recapitulated by the bulk-level analysis suggests that successful DMG treatment may require targeting multiple subpopulation dependencies via combination therapy. Conversely, the monotherapies predicted by bulk analysis are likely to fail due to selection of AC and OC states, whose dependencies are orthogonal to those of the dominant OPC state. To address this challenge, we leveraged the OncoTarget and OncoTreat algorithms to prioritize candidate MR-targeting and MR signature-reversing drugs at the individual cell state level, respectively. For this purpose, MR protein activity was assessed by VIPER analysis of synthetic bulk profiles—generated by combining the reads of all cells comprising each of the seven transcriptional states—compared to normal caudate tissue in GTEx (tumorigenic signature).

As previously described, OncoTreat relies on drug perturbational profiles from histology-matched cell lines that recapitulate the MR signature of patient-based cell states^22^ to identify whether MR activity is effectively inverted in drug- vs. vehicle control-treated cells. OncoMatch analysis confirmed that the perturbational profiles generated for bulk tissue analysis could also be utilized for this purpose. Specifically, DIPG6 and SF8628 emerged as effective models for OPC and OC states (OncoMatch p ≤ 10^-30^) and for the AC state (OncoMatch p ≤ 10^-20^), respectively (Extended Data **Fig. 4g**).

OncoTarget and OncoTreat drug predictions were highly cell state dependent, with key differences between drugs predicted to target OPC (OPCQ, OPC, and OPCC), AC (AC and OPC/AC), and OC (OC and OPC/OC) MRs (**Fig. 4f**). Not surprisingly, many drugs targeting the dominant OPC state overlapped with those predicted by bulk-level analysis, including by OncoTarget (*e.g.*, inhibitors of TOP2A, TTK, CHEK1, AURKA, etc.) and OncoTreat (*e.g.*, trametinib, palbociclib, mocetinostat, vinorelbine, etc.). Critically, however, drug predictions for minority cell states did not overlap with bulk-level predictions. Since MRs and their drug-mediated inversion are assessed based on regulons that are enriched in their physical targets, these analyses provide mechanism-based hypotheses for combination therapy.

To refine candidate drugs to a manageable number for *in vivo* validation, we applied additional criteria. Specifically, (a) only drugs with statistical significance p ≤ 10^-5^ (Bonferroni corrected) were considered, see^25^, (b) drugs with an EC_20_ > 2μm were excluded as unlikely to achieve a physiologically relevant concentrations *in vivo*; further (c) drugs targeting more than one cell state (*e.g.,* trametinib targeting OPCQ, OPC, and OPCC or venetoclax targeting AC and OPC/AC cell states) were prioritized, and (d) drugs predicted by both OncoTarget and OncoTreat (e.g. dinaciclib as a CDK2 inhibitor for OPC states) were also prioritized; finally, we considered clinical relevance, bio-availability, pediatric safety profile, and BBB permeability as additional criteria. Drugs with conclusive limited BBB permeability (*e.g.,* octreotide) were removed; yet those with mixed literature reports (*e.g.*, etoposide) were included.

Three drugs emerged as OncoTreat-predicted inhibitors of OPC states, respectively, including trametinib (highly selective inhibitor of the mitogen-activated protein kinase enzymes MEK1 and MEK2^68^), mocetinostat (an HDAC inhibitor with highest affinity to HDAC1, but also targeting HDAC2, 3 and 11^69^), and dinaciclib (a potent small molecule cyclin-dependent kinase inhibitor targeting CDK2, CDK5, CDK1 and CDK9^70^). Two additional drugs emerged as OncoTarget-predicted inhibitors of OPC and OC states, respectively, including etoposide (a TOP2 inhibitor, targeting both TOP2 isoforms: TOP2A and TOP2B^71^), and avapritinib (a CNS-penetrant PDGFRA inhibitor^72^). Similarly, two drugs emerged as OncoTreat-predicted inhibitors of AC states, including venetoclax (a highly selective BCL2 inhibitor^73^) and ruxolitinib (an ATP-competitive JAK1/2 inhibitor^73^) and an additional two were predicted by OncoTarget, including larotrectinib (a highly selective TRK inhibitor^74^) and napabucasin (a STAT3 inhibitor^75^).

While the majority of selected drugs were among the top 10 predictions by each method for the respective cell states, larotrectinib (targeting NTRK2) and Napabucasin (targeting STAT3) were selected for AC cells despite their lower rank given their state specificity. Further, the ability to target more than one state, specifically OC and OPC states, was leveraged to prioritize drugs that were not among the top 10 for either state in isolation, such as mocetinostat, etoposide (TOP2A in OncoTarget) and avapritinib (PDGFRA in OncoTarget). Selected drugs and their direct targets are summarized in **Fig. 4g**. Additional top state specific drugs for OC states, such as temsirolimus, ipatasertib, dabrafenib, and abemaciclib may be evaluated in future studies.

### *In vivo* but not *in vitro* models recapitulate DMG tumor heterogeneity

To identify models that recapitulate the heterogeneity of human DMGs, for the assessment of drug efficacy and state specificity, we analyzed scRNA-seq profiles from both *in vitro* and *in vivo* DMG models. For this purpose, we used GSEA to classify each single cell isolated from an available model by the most statistically significant (p ≤ 0.05) representative DMG cell state (Extended Data **Table 5**). Cells with the most significant enrichment for the OPCQ and/or OPC cell state were labeled as OPC, those with the most significant enrichment for the OPC/OC and/or OC cell state were labeled as OC, while those with the most significant enrichment for OPCC, AC, or OPC/AC cell states were labeled as the respective cell state.

When analyzing scRNA-seq profiles from six available DMG cell lines (n = 11,325 cells, Extended Data **Fig. 4h**), the vast majority (79.4%) of cells could not be classified (p > 0.05), while the remaining cells were all classified as OPCC (Extended Data **Fig. 4i**). We then assessed whether DMG states were better recapitulated *in vivo*, in a subcutaneous (subQ), vehicle control-treated (NMP, see methods) DIPG17 xenograft model, harboring the hallmark DMG mutation H3F3AK27M (**Fig. 5a**, Extending Data **Fig. 5a,b**). Analysis of 14,176 cells isolated from this model confirmed that all seven DMG states could be recapitulated, with individual cells co-clustering with human DMG clusters (**Fig. 5b**, Extended Data **Fig. 5e**). Consistent with human tumors, while OPC cells still dominated (66.2%), we now additionally observed AC (27.0%) and OC (2.2%) cells (p ≤ 0.05, FDR-corrected, **Fig. 5b**, Extended Data **Fig. 5f**). Critically, only 4.6% of the *in vivo* cells, compared to 79.4% of *in vitro* cells, failed to be significantly classified. Consistent with this observation, state-specific predictions for the nine candidate drugs were conserved in this model (**Fig. 6c**), confirming it as a robust *in vivo* model for the rapid evaluation of state-specific drug efficacy and specificity before further studies in a more challenging orthotopic context.

**Figure 5.**
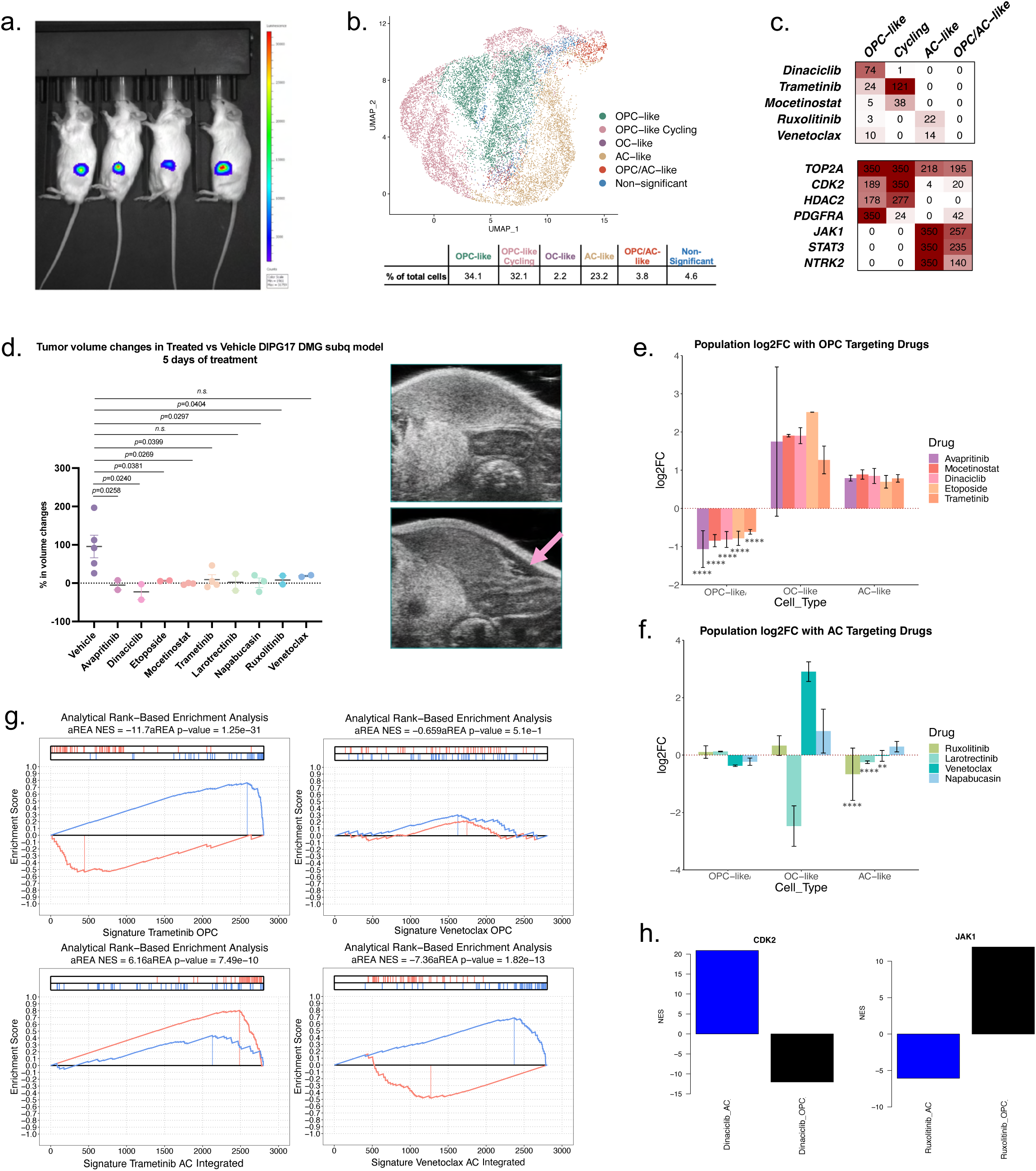
Validation of single cell OncoTarget and OncoTreat drug predictions in subcutaneous DMG patient cell line (DIPG17)-derived murine models. **a.** Bioluminescence signal in a DMG patient cell line-derived subcutaneous mouse models (DIPG17) one month after injection confirming tumor implantation. **b.** UMAP projection of 14,176 single cells from four vehicle control (NPM)-treated tumors by VIPER-inferred protein activity with cells annotated by the most representative patient population showing that the tumor model recapitulates DMG patient tumor heterogeneity. **c.** OncoTarget and OncoTreat scores, represented as -log10(Bonferroni p-values), for drugs and/or their direct targets being validated in this model showing conservation of the subpopulation specificity for the predictions compared to human tumors. **d.** Percent change in tumor volume at 5 days of treatment for each replicate across the nine drug treatment arms and vehicle control. Drug vs. vehicle volumes were compared by Welch’s t-test. Ultrasound images of the subcutaneous model pre-treatment (top) and after Avapritinib treatment for 5 days (bottom) is shown with observable tumor morphology changes indicated by the pink arrow. Log2 fold change in proportion of cells belonging to each population for each treatment condition across **e.** 38,615 single cells profiled from the 5 drug arms targeting the OPC-like cell states and **f.** 42,896 single cells profiled from the 4 drug arms targeting the AC-like cell states, compared to the 14,176 vehicle-treated cells. Bar length reflects mean log2FC across replicates in each treatment arm and error bars indicate Standard Error of the Mean (SEM). ✱ denote chi-square p-value significance comparing distribution of proportions for each population versus the rest in treated vs. vehicle cells: ✱p<0.05, ✱✱p<0.01, ✱✱✱p<0.001, ✱✱✱✱p<0.0001. **g.** Pharmacodynamic assessment of tumor checkpoint module (TCM) inversion *in vivo* after 5 days of treatment. Analytical Rank-based Enrichment Analysis (aREA) of the top (red ticks) and bottom (blue ticks) 50 DMG patient population-specific MRs in the differential protein activity signature of drug- vs. vehicle-treated single cells within that population in the DIPG17 mouse tumors. Trametinib inverting the TCM in OPC-like cells and Venetoclax conversely inverting the TCM in AC-like cells are depicted here as examples. **h.** Differential protein activity of CDK2 and JAK1 in drug- vs. vehicle-treated cells within the AC- and OPC-like populations, confirms direct targeting of CDK2 activity in OPC-like cells by Dinaciclib (left) and JAK1 activity in AC-like cells by Ruxolitinib (right).

**Figure 6.**
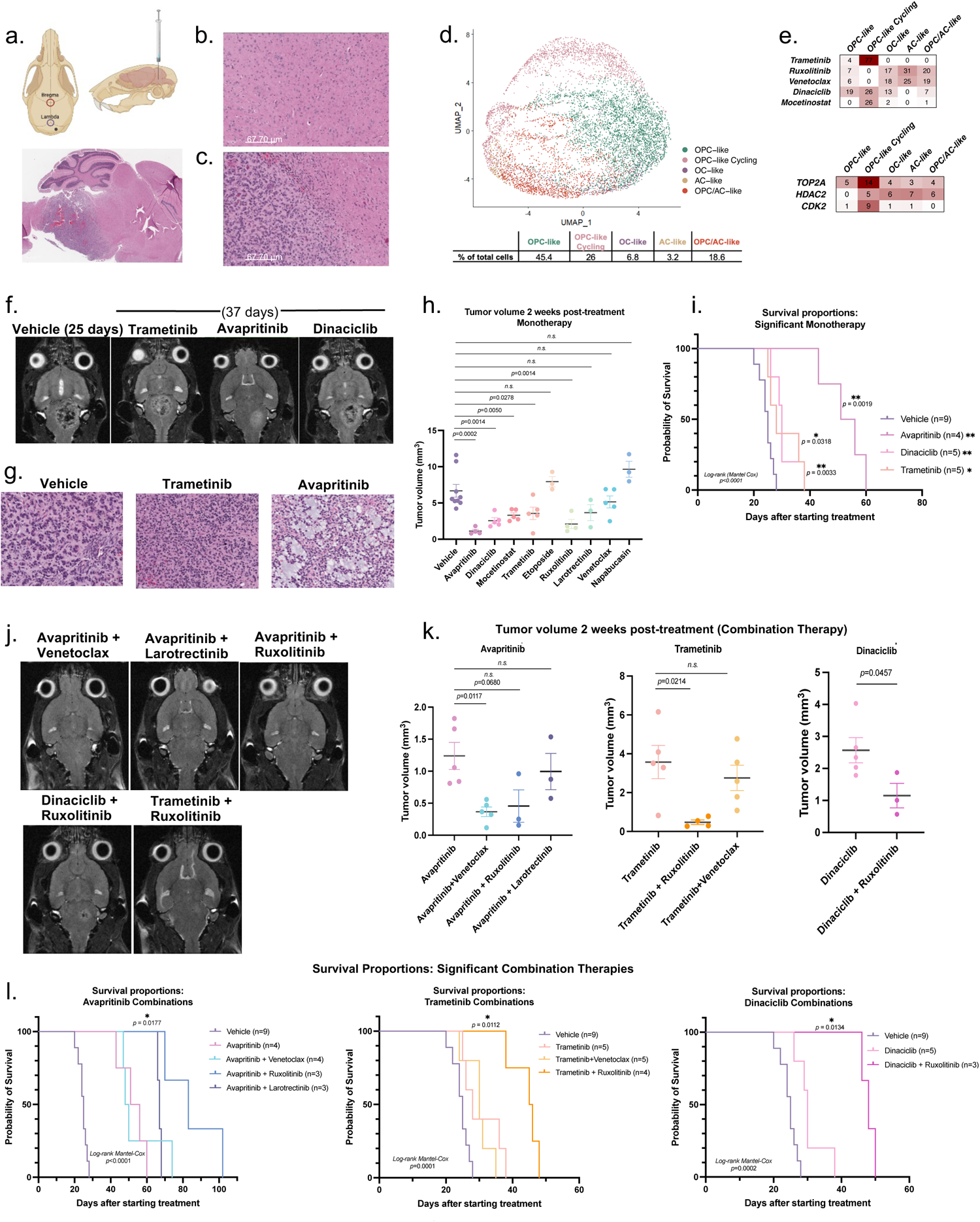
Tumor growth and survival studies with predicted drugs as monotherapy and in combinations targeting complementary cell states in a syngeneic orthotopic DMG murine model. **a.** DIPG4423 syngeneic orthotopic DMG mouse model indicating coordinates for the stereotactic injection (top). Injections were done 1mm down and to the right from the Lambda suture (purple circle). Gray dot indicates site of injection. H&E of a sagittal cut showing correct location of tumor (bottom), **b.** normal parenchyma and **c.** periphery of the tumor (showing normal parenchyma, infiltrative tumor cells and tumor). **d.** UMAP projections of 8,013 tumor single cells from two untreated DIPG4423 pontine tumors by VIPER-inferred protein activity with cells annotated based on the most representative patient subpopulation, demonstrating that the tumor model recapitulated DMG patient tumor heterogeneity. **e.** Single cell OncoTarget (bottom) and OncoTreat (top) scores as -log10(Bonferroni p-values) for drugs and/or their direct targets being validated in this model showing conservation of the subpopulation specificity for the predictions compared to human tumors. **f.** MRI images of tumors for monotherapy treatments (at 37 days) compared to vehicle control (25 days) for the three drugs significantly prolonging survival as monotherapy. Every image is one mouse at the indicated time for a particular treatment. **g.** High powered H&E stain images of end stage tumors in Vehicle, Trametinib, and Avapritinib treated mice. Avapritinib lesions demonstrate much greater and more diffuse infiltration and a notable absence of necrosis. **h.** Tumor volume after two weeks of treatment for each replicate across the five drugs targeting OPC cells on the left and four drugs targeting AC cells on the right (each ordered by significance compared to vehicle control treated tumors by Welch’s t-test). **i.** Survival analysis (Kaplan-Meier) of all monotherapy drug arms with significant differences in survival with respect to vehicle controls assessed by log-rank test. **j.** MRI images of tumors for combination treatments of significant OPC-targeting drugs (shown in i) and AC-targeting drugs at 37 days demonstrating smaller size as compared to vehicle control and the monotherapies depicted in f. **k.** Tumor volume after two weeks of treatment for (left to right) Avapritinib, Trametinib and Dinaciclib combinations with AC-targeting drugs. Tumor volume in combination arms was compared to the respective monotherapy by Welch’s t-test. **l.** Survival analysis (Kaplan-Meier) of vehicle, monotherapy, and combination therapy treatment arms in the context of (left to right) Avapritinib, Trametinib, and Dinaciclib as the monotherapy. Four out of six combination therapies significantly improved survival compared to the respective monotherapy by log-rank test. ✱ denote p-value significance, where ✱p<0.05, ✱✱p<0.01, ✱✱✱p<0.001, ✱✱✱✱p<0.0001. *n.s.* = Not significant.

### Predicted drugs specifically deplete their target cell states and reduce tumor growth in a subQ DIPG17 xenograft

To validate the subpopulation-specific effects of candidate drugs— including five targeting OPC states and four targeting AC states—juvenile NOD.Cg Prkdcscid/J (NSG^TM^) mice were inoculated subcutaneously with luciferase-tagged DIPG17 cells. Upon reaching ∼100 mm^3^, tumor-bearing mice were randomly assigned to distinct drug or vehicle control arms and assessed by ultrasound longitudinally, pre- and post-treatment, to monitor tumor volume changes (Extended Data **Fig. 5a,b**). Animals were treated for five days—an appropriate time point to capture drug pharmacodynamic effects on MR activity and subpopulation depletion, while avoiding significant cell death which would confound the analysis—and euthanized two hours after the last dose. Depletion of predicted cell states in drug- vs. vehicle control-treated mice was assessed by single cell analysis. Specifically, 95,687 high quality scRNA-seq profiles were generated, representing tagged single cells isolated from 24 samples—including 9 drug treatment arms in replicate (n = 2 or 3) and a vehicle control arm in quadruplicate (Extended Data **Fig. 5d**). Cell state was assessed as described in the previous section—confirming that samples adequately recapitulated human tumors’ heterogeneity (Extended Data **Fig. 5f**, Extended Data **Table 6**)—and was then used to assess differential cell state occupancy in drug vs. vehicle control-treated animals.

Drug-induced cell state depletion was highly consistent with predictions for 8 of 9 tested drugs (**Fig. 5e,f**). Specifically, by Χ^2^ analysis (Extended Data **Table 5**), all five drugs predicted to target the OPC states (OPCQ, OPC, OPCC) induced highly statistically significant OPC (OPCQ and OPC) and OPCC cell depletion, including trametinib (p_OPC_ = 4.44×10^-91^, p_OPCC_ = 1.34×10^-143^), dinaciclib (p_OPC_ = 9.34×10^-138^, p_OPCC_ = 2.57×10^-122^), mocetinostat (p_OPC_ = 2.55×10^-319^, p_OPCC_ = 6.11×10^-03^), etoposide (p_OPC_ = 2.26×10^-246^, p_OPCC =_ 5.95×10^-20^), and avapritinib (p_OPC_ = 8.13×10^-201^, p_OPCC_ = 7.27×10^-56^) (**Fig. 5e**, Extended Data **Fig. 6a**, Extended Data **Table 6**). In sharp contrast, but also as predicted, these drugs induced highly significant AC cell (AC and/or OPC/AC) repletion (p_AC_ = 0, p_AC_ = 5.58×10^-142^, p_AC_ = 1.61×10^-160^, p_AC_ = 1.54×10^-88^, and p_AC_ = 5.58×10^-82^, respectively), confirming that their use as monotherapy would eventually induce selection of cell states eliciting adaptive drug resistance. OC cells were also significantly repleted by these drugs, especially by avapritinib (p_OC_ = 0) and etoposide (p_OC_ = 1.29×10^-218^).

Consistent with these observations, three of four drugs predicted to target AC states induced significant AC cell (AC and/or OPC/AC) depletion, including ruxolitinib (p_AC_ = 4.68×10^-17^, p_OPC/AC_ = 0.87), venetoclax (p_AC_ = 5.72×10^-08^, p_OPC/AC_ = 1.02×10^-10^), and larotrectinib (p_AC_ = 8.21×10^-03^, p_OPC/AC_ = 7.11×10^-35^) (**Fig. 5f**, Extended Data **Fig. 6b**, Extended Data **Table 6**). Napabucasin failed to deplete AC states but induced significant depletion of OPC cells (p_OPC_ = 5.11×10^-165^, p_OPCC_ = 8.60×10^-41^). The effects of ruxolitinib, venetoclax, and larotrectinib on OPC and OC states was variable, yet consistent with model-specific predictions. Ruxolitinib emerged as the most effective and specific inhibitor of the AC state, resulting in significant repletion of OPC cells (p_OPC_ = 9.28×10^-3^), and non-significant effects in OPCC (p = 0.075) and OC cells (p = 0.17). Venetoclax, which was also predicted to target OPC cells in this model, depleted them (p_OPC_ = 3.69×10^-22^, p_OPCC_ = 3.98×10^-57^), while inducing highly significant repletion of OC cells (p = 0). Of all the drugs, only larotrectinib induced highly significant depletion of OC cells (p_OC_ = 1.98×10^-26^). Indeed, OC and AC states presented increased NTRK2 activity in this model.

We then assessed pre- vs. post-treatment tumor volume differences in drug vs. vehicle control-treated animals by ultrasound (i.e., right before euthanizing). Despite the short (5d) treatment cycle and small treatment arm size, modest tumor volume reductions were induced by virtually all drugs determined to target the dominant OPC states, with dinaciclib and avapritinib showing the most significant effects (**Fig. 5d**). Napabucasin—which was shown to target OPC rather than AC cells—and ruxolitinib also induced significant tumor growth reduction.

### MR activity inversion is confirmed *in vivo*

State-specific MR activity inversion by OncoTreat-predicted drugs was then assessed following each drug treatment (*i.e.,* drug MoA conservation). This is important because the effect was predicted based on perturbational profiles *in vitro*. All seven OncoTreat-predicted drugs (mocetinostat, trametinib, and dinaciclib targeting OPC states and venetoclax and ruxolitinib targeting the AC state) induced significant activity inversion of top and bottom 50 MRs of depleted cell states (p ≤ 0.05, by 1-tailed aREA), in drug versus vehicle-control treated animals (**Fig. 5g**, Extended Data **Fig. 7a,b**). While AC cell-specific MR activity inversion by ruxolitinib was just above statistical significance (p = 0.058), JAK1 activity inhibition was highly significant, supporting its assessment *in vivo* based on OncoTarget prediction (**Fig. 5h**). Further, the analysis confirmed that PDGFRA inhibition by avapritinib induced significant MR activity inversion in OPC cells (Extended Data **Fig. 7a**). Additionally, larotrectinib inhibited NTRK2 activity in the two states it most significantly depleted (OC and OPC/AC) (Extended Data **Fig. 7c**). Napabucasin, an AC state-specific, OncoTarget-predicted STAT3 inhibitor, did not inhibit STAT3 activity *in vivo* providing a potential rationale for its failure to deplete the AC state (Extended Data **Fig. 7d**).

### Generation of a syngeneic orthotopic DMG murine model and validation of model fidelity by scRNA-seq

Following drug specificity validation in the subQ model, we proceeded to test drugs in a syngeneic orthotopic model generated by stereotactic injection of PDGF-B^+^ p53^-/-^ H3.3 K27M murine glioma cells (DIPG4423) into the pons of B6(Cg)-Tyrc-2J/J (B6) mice (**Fig. 6a-c**). This is a well-characterized brainstem orthotopic, immunocompetent DMG model^76^. Tumor engrafting in the correct location was confirmed by MRI and H&E (**Fig. 6a-c**, Extended Data **Fig. 8a,b**). Similar to the subQ model, analysis of 8,013 scRNA-seq profiles of cells isolated from two untreated tumors confirmed that this model recapitulates the heterogeneity and state-specific drug predictions of human DMGs (**Fig. 7d-e**, Extended Data **Fig. 8****c-f**).

### Monotherapies targeting predominant tumor populations improve survival in a syngeneic orthotopic DMG model

MRI imaging, at 12d after stereotactic injection, was used to confirm presence of tumor in the correct location. Tumor-bearing mice were then enrolled in independent monotherapy treatment arms (n = 3 – 5 mice per arm), including the nine drugs evaluated in the subQ model and vehicle control (see Methods for treatment details). Drug response was evaluated by tumor volume measured at 2 weeks post treatment by MRI and by overall survival (Extended Data **Fig. 9a,b**, Extended Data **Fig. 10a,b**). Drug toxicity was monitored by weight loss and other parameters (see Methods). Napabucasin was included in this study despite its failure to induce AC cell depletion, because of its confirmed depletion of OPC cells, inversion of the associated MR signature, and significant tumor volume reduction in the subQ model. Drugs targeting OPC states induced statistically significant tumor growth reduction compared to vehicle control (Welch’s t-test p ≤ 0.05), except for etoposide, likely due to lower BBB permeability (**Fig. 6f,h**). Of these, avapritinib, dinaciclib, and trametinib also showed significant survival extension (log-rank p ≤ 0.05), with median survival of 53.5d, 30d and 28d, compared to 25d for vehicle control, respectively (**Fig. 6i**, Extended Data **Fig. 9b**, **10b**).

Among drugs targeting the AC state, only the top prediction (ruxolitinib) induced significant tumor growth reduction at 2 weeks (Welch’s t-test p = 1.4×10^-3^); however, consistent with the fact that these states harbor only a small minority of malignant cells, none of these drugs induced significant survival benefit (Extended Data **Fig. 9b**).

MRI imaging suggests that avapritinib treated tumors are more diffusely infiltrating (**Fig. 6f**, Extended Data **Fig. 10a**). Postmortem H&E staining of avapritinib, trametinib and vehicle control-treated tumors confirmed that—compared to vehicle control and trametinib—avapritinib affects tumor morphology, resulting in lesions with greater diffuse infiltration and notable absence of necrosis, which was instead evident in trametinib-treated tumors (**Fig. 6g**, Extended Data **Fig. 11**). This is consistent with previous reports of similar morphological features induced by sunitinib—a small molecule inhibitor of multiple receptor tyrosine kinases, including PDGFR, that was reported to induce similar morphological features in glioblastoma (GBM)^77^

### Combination therapy, targeting complementary cell states, further improves survival

To assess whether combination therapy targeting complementary cell states may improve tumor control, we enrolled mice in distinct treatment arms (n = 3 – 5 replicates per arm) testing drug pairs shown to target the OPC and AC states, representing the most populated complementary cell states. For each combination, tumor volume (at 2 weeks post treatment) and overall survival of drug-treated animals were compared to vehicle control-treated as well as to monotherapy-treated animals. To target OPC states, we selected avapritinib, dinaciclib, and trametinib, three drugs that elicited the most significant survival improvement. To target AC states, we selected ruxolitinib, which also induced tumor volume reduction, as well as venetoclax. We also assessed the combination of avapritinib (OPC targeting drug) in combination with larotrectinib, the only drug that induced effective depletion of the OC state.

Overall, all combinations were well tolerated, except for dinaciclib/venetoclax, which was excluded from the study given significant early weight loss (see Methods for further details on treatment regimens). Compared to vehicle controls, all combinations demonstrated statistically significant reductions in tumor growth (Extended Data **Fig. 12b**). More critically, several combinations also significantly outperformed the individual monotherapies, including avapritinib/venetoclax (p = 0.01), trametinib/ruxolitinib (p = 0.02), and dinaciclib/ruxolitinib (p = 0.046); avapritinib/ruxolitinib showed a trend towards significance (p = 0.068) (**Fig. 6j-k**, Extended Data **Fig. 12****-14**). More relevant, an even greater number of combinations significantly outperformed their respective monotherapies in terms of survival, including virtually all combinations with ruxolitinib, such as avapritinib/ruxolitinib (83d vs. 53.5d median survival, p = 0.02), trametinib/ruxolitinib (45.5d vs. 28d, p = 0.01), and dinaciclib/ruxolitinib (48d vs. 30d, p = 0.01) (**Fig. 6l**). Avapritinib/larotrectinib, targeting OPC and OC states, also significantly improved survival compared to avapritinib alone (67d vs. 53.5d, p = 0.02), suggesting that a combination of avapritinib, ruxolitinib, and larotrectinib, targeting all three states, may outperform the corresponding pairwise combinations, which will be tested in future studies. Since the drugs targeting AC and OC states did not significantly improve outcome as monotherapy, all statistically significant combinations were by definition synergistic.

While avapritinib/venetoclax induced significant tumor volume reduction over the first 6 weeks, compared to avapritinib alone (Extended Data **Fig. 14**), overall survival did not significantly improve (**Fig. 6l**). While this is possibly related to the emergence of a resistant population or to the borderline significance of the effect of venetoclax on AC cells, an equally plausible rationale may be related to the fact that tumor morphology was critically affected by this treatment compared to other combinations (e.g., trametinib/venetoclax) and vehicle control, with tumors showing less necrosis and a much more diffuse, infiltrative growth pattern (Extended Data **Fig. 15**). Indeed, these mice died acutely from seizures rather than from gradual emergence of neurological changes.

Overall, avapritinib/ruxolitinib produced the most significant outcome improvement in these highly aggressive orthotopic DMG models, specifically ∼3-fold and ∼1.5-fold compared to vehicle control and avapritinib monotherapy, respectively.

### Synergy studies confirm lack of synergy in individual subpopulations

To validate that the synergistic effect on tumor growth and survival *in vivo* are mediated by the differential sensitivity of different cell states to the drugs, rather than to cell-autonomous synergy in the predominant OPC states, we performed BLISS independence studies for avapritinib/ruxolitinib in three DMG cell lines (DIPG4423, SF8628 and DIPG36), which are mostly comprised of OPC cells. Specifically, if, as predicted, the effect is mediated by the differential sensitivity of different cell states to each drug, then the synergy revealed by the *in vivo* assays should not be recapitulated in cell lines that are effectively mono-state. Indeed, the avapritinib/ruxolitinib combination demonstrated no synergy across these cell lines (Extended Data **Fig. 16a**). As such, the statistically significant synergistic effect elicited by these drugs *in vivo* would have been completely missed by any *in vitro* screen. Additionally, as monotherapy, ruxolitinib caused limited cell death in all three cell lines, as evidenced by the dose-response curves, with one cell line (DIPG36) successfully expanding while on treatment (Extended Data **Fig. 16b**) as might have been expected given the lack of AC cells in these cell lines.

## Discussion

DMG is a universally fatal childhood CNS malignancy affecting the midline structures of the brain. Despite significant progress in its biological and genomic characterization, translation to therapeutic benefit has been lagging and treatment options to prolong life are still mostly limited to palliative radiation therapy. Critical challenges include lack of actionable molecular alterations and significant intra-tumor heterogeneity. Global H3K27me3 hypomethylation, which represents the hallmark of DMG, induces broad epigenetic dysregulation and activation of devastating transcriptional programs^5–9^. H3 alterations can co-segregate with other somatic driver mutations (i.e., TP53, ACVR1, EGFR), resulting in a high level of interpatient heterogeneity^5–9^.

H3 dysregulated programs are further modulated by highly pleiotropic posttranslational modifications—including acetylation, methylation, phosphorylation and ubiquitination of a large number of proteins—thus further challenging the discovery of targeted therapies^5–9^. In addition, recent studies have unveiled a remarkable heterogeneity of DMGs, with several epigenetically distinct populations—likely presenting orthogonal drug sensitivity—co-existing in virtually all tumors ^27,51^. As a result, decades of clinical trials—including with drugs that directly or indirectly target established mutations—have yielded underwhelming results^2,78–80^ and novel options are urgently needed.

This study pioneers the concept that systematic, mechanism-based identification of drugs targeting orthogonal non-oncogene dependencies in molecularly distinct, yet co-existing malignant subpopulations can lead to rapid nomination and validation of combination therapies that can significantly improve outcome compared to the corresponding monotherapies. Specifically, we extend a network-based methodology for the identification and pharmacologic targeting of non-oncogene dependencies, representing MRs of tumor transcriptional state, by VIPER analysis of bulk RNA-seq profiles, to the study of molecularly distinct malignant subpopulations at the single cell level. Given their heterogeneity and lack of response to monotherapy, DMGs constitute an ideal framework for the validation of this approach.

Bulk level analysis was highly effective in nominating MR proteins representing non-oncogene dependencies of the dominant DMG tumor compartments, as validated by pooled CRISPR KO screens, including FOXM1 and CENPF previously reported as synergistic MRs of several tumors associated with poor outcome, including metastatic castration resistant prostate cancer^19^. However, several MR-reversing drugs predicted by bulk analysis, had already failed in the clinic and induced only marginal improvement in DMG syngeneic model survival. To address this challenge, we leveraged the VIPER algorithm to identify seven molecularly distinct cell states, and associated MR signatures, that co-exist in virtually every DMG. We then generated a large-scale set of DMG specific perturbational profiles and used them to identify drugs targeting each state, based on the state-specific inversion of their MR protein activity, via the OncoTreat and OncoTarget algorithms. While OPC, OC, and AC subpopulations were previously reported for DMG, the identification of additional, finer grain cell states in this work has two important consequences. First, it avoids diluting the identification of cell state-specific candidate targets arising from averaging across molecularly distinct subpopulations. Additionally, it facilitates the identification of drugs that may more specifically target each of the identified states.

Validation studies in subQ DMG xenografts confirmed the efficacy and state specificity of 88% of predicted drugs, independent of genetic alterations routinely associated with sensitivity to these drugs. As expected, several drugs (avapritinib, trametinib, and dinaciclib), predicted to target the majority OPC cell population, overlapped with drugs predicted by bulk-level analysis and induced significant survival benefit in a syngeneic, orthotopic DMG model. In contrast, only one drug (ruxolitinib) targeting minority AC cell states had any significant effect on tumor growth, and no AC-targeting drugs improved survival as monotherapy. Yet, virtually all of the pairwise combinations targeting complementary cell states significantly and synergistically outperformed their associated monotherapies, even though the effect was not mediated by cell autonomous mechanisms.

Specifically, avapritinib/ruxolitinib, trametinib/ruxolitinib, dinaciclib/ruxolitinib, and avapritinib/larotrectinib significantly improved survival compared to their monotherapies as well as vehicle control. In particular, the combination of avapritinib and ruxolitinib had remarkable survival benefits compared to avapritinib alone and vehicle control (median survival of 83 days vs. 53.5 days and 25 days, respectively). Consistently, ruxolitinib had the strongest and most specific effect of all drugs on AC cells. The synergistic effect of avapritinib and larotrectinib was also notable, since the latter was the only drug presenting specific activity on OC cells. As a result, overall survival correlated with the cell state specificity of individual drug effects. This suggests that a triple combination therapy, including avapritinib, ruxolitinib, and larotrectinib, may outperform all the pairwise combinations. This will be tested in future studies.

Children with DMG face a heartbreaking diagnosis, with very limited treatment options. Altogether, this study improved our understanding of the oncologic programs driving unique DMG tumor cell states, defined intra-tumor heterogeneity, and ultimately prioritized highly promising drug combinations for further preclinical and clinical testing. However, the proposed methodology is not restricted to DMGs and can be used in virtually any tumors comprising multiple molecularly distinct, co-existing cell states.

A potential limitation of the approach is that OncoTreat analyses require large-scale perturbational profiles in cell lines or primary cells that faithfully recapitulate the MR activity signature of each co-existing cell state. This may be challenging for some tumors where *in vitro* culture conditions may ablate some cell states. However, with single cell technologies improving at a rapid pace, it may soon be possible to generate such profiles in short term primary cell cultures isolated by FACS from patient tumors or PDX models. In contrast, a potential advantage of this approach is that since drug synergy is not elicited by cell autonomous mechanisms but rather by independent ablation or reprograming of distinct cell states, predicted combinations may be also effective in sequential rather than concurrent administration, thus reducing potential co-administration toxicity. An additional advantage, largely responsible for the high validation rate of drug predictions, is that both non-oncogene dependencies and the drugs that target them are predicted on a mechanistic basis. Indeed, MRs are predicted by VIPER based on the differential expression of their targets, which we have shown to be highly enriched (70% to 80%) in bona fide mechanistic, physical targets. Similarly, prediction of optimal drugs is based on MR activity inversion, which is also based on assessing the differential activity of each MR in drug vs. vehicle control-treated samples, again based on the differential expression of their transcriptional targets.

Rational prediction of drug combinations that provide effective survival benefit is challenging and *in vivo* validation is still largely serendipitous. In contrast, our analysis predicted drugs that not only validated at an 88% rate in terms of specifically targeting individual tumor subpopulations but, more importantly, induced even more significant survival extension when used in combination to target cell states with orthogonal drug sensitivity. This suggests that this highly generalizable, network-based framework may be effectively used for the rational prediction of novel, mechanism-based combination therapies.

## Methods

### DMG patient bulk RNA-seq datasets

We downloaded publicly available DMG patient RNA-seq data and associated sample and clinical annotations for 122 patients from three cohorts (Extended Data **Table 1**). Two were obtained from the Pediatric Brain Tumor Atlas (PBTA) through the Gabriella Miller Kids First Data Resource Center. This included samples from the PBTA:PNOC^33^ (n = 31) and PBTA:CBTTC^34^ (n = 47) cohorts. All samples with the diagnosis “Brainstem glioma – diffuse intrinsic pontine glioma” or “diffuse midline glioma, H3K27M mutant, WHO grade IV” or “diffuse midline glioma” were included. Additionally, clinical annotations were reviewed with a pediatric neuro-oncologist for all samples labeled as “high grade glioma WHO Grade III-IV” and those occurring in midline structure (brainstem, thalamus, spinal cord) with relevant molecular features that would be treated as a DMG clinically were included. The third cohort named St. Jude (n = 44) included samples with the diagnosis of DIPG or, determined to be a high-grade glioma occurring in a midline structure and likely to clinically behave like DIPG, from the following publication^9^. To minimize technical batch effects arising from unique library preparation, sequencing protocols, and gene quantification methodologies across the datasets, we obtained raw FASTQ files or BAM files, from which FASTQ files were generated using the Picard Tools program “SamToFastq” version 2.25.0. We performed gene quantification with Kallisto, using the complementary DNA (cDNA), the non-coding RNA (ncRNA) and the Genome Reference Consortium Human Build 38 Organism (GRCh38): *Homo sapiens* (human) as a reference genome. Data was obtained and analyzed under IRB protocol #AAAS5252.

### DMG bulk regulatory network

Gene counts were normalized as transcripts per million (TPM). The ARACNe3 most recent implementation of the ARACNe-AP codebase was used. The goal of ARACNe3 is to reverse engineer a context-specific transcriptional regulatory network consisting of bivariate transcriptional regulatory interactions between a set of predefined putative transcriptional regulators and potential transcriptional targets. Candidate transcriptional regulators included 1,813 transcription factors (genes annotated in the Gene Ontology Molecular Function database (GO) as GO:0003700—‘DNA binding transcription factor activity’, or as GO:0003677—‘DNA binding’ and GO:0030528—‘Transcription regulator activity’, or as GO:0003677 and GO:0045449—‘Regulation of transcription’), 969 transcriptional co-factors (a manually curated list, not overlapping with the transcription factor list, built upon genes annotated as GO:0003712—‘transcription cofactor activity’ or GO:0030528 or GO:0045449) or 3,370 signaling-pathway-related genes (annotated in the GO Biological Process database as GO:0007165— ‘signal transduction’ and in the GO Cellular Component database as GO:0005622—‘intracellular’ or GO:0005886—‘plasma membrane’). ARACNe3 significantly improves rejection of false positives by using an analytical null model based on the principle of maximum entropy and substantially reduces the number of samples and bootstraps necessary to generate accurate networks^24,30^. As such, it is especially valuable for the analysis of tumors where sample availability is at a premium, as in the case of DMG.

To avoid technical artifacts resulting from batch effects across datasets as well as transcriptional bias between autopsy versus biopsy specimens, we generated a consensus network across 120 DMG gene expression profiles containing autopsy vs. biopsy information, integrating individual ARACNe3 networks reverse engineered from the following sample sets: CBTTC_Autopsy_Network (15 samples), CBTTC_Biopsy_Network (30 samples), PNOC_Biopsy_Network (31 samples), StJude_Autopsy_Network (28 samples), StJude_Biopsy_Network (16 samples). We accounted for the small sample size in each individual network by setting a higher FDR threshold and integrating the results weighted by the number of samples per network so that only the most robust interactions remained. As such, each network was pruned with a mutual information threshold calculated to control the FDR at 0.50 and subsequently by MaxEnt pruning (*i.e.,* DPI pruning^24^). Regulons from each network were integrated using a consensus scoring approach where Association Weight (*i.e.* likelihood) and Association Mode (*i.e.* tfmode) values were averaged for each target over all networks, and each networks’ data was weighted according to the number of gene expression profiles used to reverse engineer the network.

### Gene expression signatures and protein activity inference

RNA-seq counts for the 122 patient samples and 246 normal caudate tissue samples from GTEx were normalized using log2(Counts Per Million (CPM) + 1). We generated an internal gene expression signature for each sample on a gene-by-gene basis by z-score transformation scaling by the mean and standard deviation (sd) done within the sample’s specific dataset cohort and split by autopsy or biopsy (a total of 120 samples had this information and were included). Further, we generated an external gene expression signature for all 122 samples comparing DMG tumor samples relative to the normal GTEx caudate samples, again by z-score transformation scaling each gene to the mean/sd across the GTEx caudate cohort. Protein activity, reported as a normalized enrichment score (NES), for 6,138 regulatory proteins was then computed independently for each gene expression signature (internal and external) on a sample-by-sample basis by the VIPER function in the VIPER package (Bioconductor) using the aforementioned DMG bulk network with regulons pruned to the top 100 edges.

### Unsupervised clustering analysis to define DMG subtypes

To define distinct subtypes of DMG, we performed unsupervised clustering using the PAM method (PAMK R function from the FPC package in R) on the protein activity signature generated with the internal reference. The PAMK function optimizes the number of clusters by average silhouette score (SS) width. Distance, or the similarity between samples, was computed using the viperSimilarity function, available from the VIPER package (Bioconductor), using the top 100 regulators. As a comparison, similar unsupervised clustering by the PAMK function was performed on the gene expression signatures as well with distance between samples computed by Spearman correlation.

### Survival studies between DMG subtypes

When assessing differences in survival by DMG subtype, samples with a survival time of 0 (120→111), samples with unknown survival time (111→109) and outliers (a patient with survival of 4 days and 2 patients with survival >7 years, 109→106) were removed. We then assessed differences in overall survival by cluster assignment using the survfit function in the survival package in R. We generated a Kaplan-Meier survival curve with samples split by DMG subtype (C1 vs. C2) and computed significance of survival differences by log-rank test with cutoff for significance set at p ≤ 0.05.

### Pathway enrichment analysis for DMG subtype MRs

For each bulk protein activity cluster, we used the Stouffer method to integrate protein activity scores across the samples within that cluster weighted by the sample-specific SS for each cluster. We then performed an overrepresentation analysis using a hypergeometric test (via WebGestalt) across gene pathway databases for the top 50 most activated MRs for each cluster, providing the 6,381 regulators assessed as the reference list. We looked at overrepresentation in the following pathway databases: MSigDB Hallmarks (database V2023), Reactome, and WikiPathways. For advanced parameters, we set the minimum number of genes for a category to 5, and multiple hypothesis testing correction was performed using the Benjamini-Hochberg (BH) method with statistical significance set at an FDR ≤ 0.05.

### Culture of DMG cell lines

SU-DIPG-IV (DIPG4), SU-DIPG-VI (DIPG6), SU-DIPG-XVII (DIPG17), SU-DIPG-XXXVI (DIPG36) and SF7761 cells were cultured in Knockout DMEM/F-12 (Gibco #12660012), Stempro Neural Supplement (Gibco #A1050801), EGF Recombinant Human Protein (Gibco #PHG0313), FGF Basic (aa 10 155) Recombinant Human Protein (Gibco #PHG0021), 1% Penicillin-Streptomycin Solution and 1% L-glutamine. SF8628 cells were cultured in DMEM (Gibco #11995073), 10% Fetal Bovine Serum (FBS) (Sigma-Aldrich F2442), 1% Penicillin-Streptomycin Solution (Sigma-Aldrich P4458) and 1% L-glutamine (Corning #25-005-CI). DIPG4423 cells were cultured in Mouse Neurocult media (Stem Cell Technologies #05700), Mouse and Rat proliferation supplement 10% (Stem Cell Technologies #05701), Recombinant Human FGF (20 ng/mL) (PeproTech (100-18B), Recombinant Human EGF (10 ng/ml) (PeproTech AF-100-15), Heparin (2μg/ml) (Stem Cell Technologies #07980) and 1% Penicilin-Streptomycin Solution (Sigma-Aldrich P4458).

### Generation of Cas9 expressing DMG cell lines

For lentivirus production, HEK293T cells were maintained in DMEM (Gibco #11995073) containing 10% FBS (Gemini Bio 100106/500). Transfection was performed by using Polyethylenimine (PEI 25000). 6 μg of plasmid DNA, 4.5 μg of psPAX2 (Addgene #12260), 1.5 μg of pMD2.G (Addgene #12259) and 18 μl of 2 mg/ml PEI 25000 was mixed in Opti-MEM (Gibco #31985062) and then added into each 10-cm-dish culture of HEK293T cells. The culture media was changed with fresh media at 12 hours post transfection. The supernatant was collected at 48 hours and 72 hours post transfection and filtered with a 0.45-micron filter. Virus in the supernatant was precipitated with PEG-6000 (Santa Cruz Biotechnology sc-302016), suspended in appropriate media and pooled. For the sgRNA library of transcription factors, the scale of lentivirus production was calculated to maintain representation > 1,000.

To generate Cas9-expressing cells, DMG cells (DIPG4, DIPG17, and SF8628) were digested, spin-infected with lentivirus of LentiV_Cas9_puro vector (gift from Dr. Christopher R. Vakoc to Dr. Zhiguo Zhang) with an infection MOI ≈ 1.5 at 1,200 rpm at 25℃ for 1 hour. The media was refreshed at 24 hours post infection and 1 μg/ml Puromycin was added for 3 days. The cells were then digested and infected again. The media was refreshed at 24 hrs post 2^nd^ infection and 1.5 μg/ml Puromycin was added for 3 days. The expression of Cas9 was confirmed by Western blot with anti-Cas9 antibody (CST #65832) and the cutting efficiency was confirmed with sgRNA targeting *CDK1* and *PCNA*.

### Generation of luciferase-expressing cells

DMG cells (DIPG4, DIPG17 and SF8628) were digested, spin-infected with lentivirus of firefly-LUC_pHR-SIN-CSGW (gift from Dr. Rintaro Hashizume to Dr. Zhiguo Zhang) with an infection MOI ≈ 2.5 at 1,200 rpm at 25℃ for 1 hour. The media was refreshed at 24 hours post infection. The expression of luciferase was confirmed by immunofluorescence staining with anti-Luciferase antibody (Abcam #ab21176) and the cells with > 90% luciferase-positive were used for experiments.

### sgRNA library generation and CRISPR-Cas9 pooled screenings

Transcription factor (TF) domain-focused sgRNA library (gift from Dr. Christopher R. Vakoc to to Dr. Zhiguo Zhang) was amplified using ElectroMAX Stbl4 Competent Cells (Invitrogen #11635018) according to manufacturer’s instructions with a representation > 1,000 and used to produce lentivirus as described above. Cas9-expressing cells were spin-infected with lentivirus of the sgRNA library at 1200 rpm at 25℃ for 1 hour, with a MOI ≈ 0.3 and representation >1,000. The cells were cultured with a representation > 1,000 at any time point during the screenings. Samples were collected at starting time point (day 3 post infection, T0) and end time point (12 doubling times after starting time point, T1). Genomic DNA was extracted using QIAGEN DNeasy Blood and Tissue Kit (QIAGEN #69504) according to manufacturer’s instructions. Barcoded sequencing libraries were prepared as previously described^72^, pooled and analyzed by paired-end sequencing using NextSeq (Illumina) for each of the three DMG cell lines. The TF domain-focused sgRNA library was comprised of ∼6 sgRNA per gene (1,486 genes total, including 1,373 TFs, 62 core-essential positive controls and 51 non-targeting negative controls) (Extended Data **Table 2**).

### Analysis of CRISPR/Cas9 pooled KO experiments by MAGeCK

CRISPR experiments were analyzed using the MAGeCK algorithm^10^, using Robust Rank Aggregation (RRA) and total read count normalization, with default settings^10^. Each replicate was analyzed independently by comparing the late time point (T1) against day 3 (T0). A CRISPR/Cas9 essentiality signature for each gene in each cell line was computed as the log-fold change. Results were analyzed independently for each cell line. Then, core-essential genes were subtracted from the analysis using the Cancer Dependency Map, DepMap^12^, as a reference. Results were then integrated across the three cell lines as median log fold change to create a DMG essentiality signature.

### Quality control of CRISPR/Cas9 pooled KO experiments

An Empirical Cumulative Distribution Function (ECDF) of the log-fold change (FC) of each TF between T0 and T1 was established by performing a Kolmogorov-Smirnov’s test. Core essential positive controls and non-targeting negative controls had an overall more negative and positive FC distribution, respectively. It is critical to see an enrichment of the core-essential genes in the top essential genes (negative FC) and the non-essential genes should not be significant (FC close to zero), which is what was observed in these functional screens (Extended Data **Fig. 2b**)

### DMG cell line model fidelity to DMG patients by OncoMatch

Bulk RNA-seq data was obtained for six DMG cell lines (DIPG4, DIPG6, DIPG17, DIPG36, SF7761, and SF8628) and gene counts generated by Kallisto from raw FASTQ files as described above for DMG patients. Counts were normalized by log2(CPM+1) and gene expression signatures computed relative to the GTEx caudate samples by z-score transformation (subtracting the mean expression level across the GTEx caudate samples and dividing by the standard deviation). Protein activity for 6,138 regulatory proteins was then computed in each cell line by the VIPER algorithm using the DMG bulk network. Cell line model fidelity to the 122 DMG patients was evaluated based on MR conservation, as assessed by analytic-rank enrichment analysis (aREA) within the OncoMatch function^25^. Specifically, we evaluated enrichment of the top/bottom 50 MRs (GTEx caudate signature) in each patient sample in the protein activity signature of each of the six cell lines (aREA p ≤ 10^-5^ was considered a significant match). Among the available cell lines, we selected SF8628 and DIPG6 as the ones that together best recapitulated DMG patients by OncoMatch score and had adequate growth parameters as suitable models to assess drug mechanism of action within the DMG context.

### Drug perturbation studies by PLATE-seq

#### Drug dose response curves (DRC)

An established drug library, including 372 FDA-approved and late-stage experimental oncologic compounds, was used (Extended Data **Table 4**). To determine the maximum sublethal concentration of each drug—defined as its 48h EC_20_ concentration—SF8628 and DIPG6 cells were seeded onto 384-well tissue culture plates (Greiner 781080 Monroe, NC; columns 1-24, 50 μL volume of adequate media) at a density of 1,000 cells per well for DIPG6 and 500 cells per well for SF8628. Plates were incubated at room temperature for 30 minutes, and then overnight in an incubator (37°C, 5% CO2). The next morning (approximately 12 hours after seeding) compounds were added using the Echo 550 acoustic dispenser. After 40μL of media were removed from each well, each drug was dispensed in ten doses, with each plate run in triplicate. After 48 hours, 25μL of CellTiter-Glo (Promega Corp, Madison, WI) were added to each well. Plates were shaken for 5 minutes before enhanced luminescence readout (EnVision, Perkin Elmer, Shelton, CT). All values were normalized by internal DMSO controls contained on each plate and analyzed for EC_20_ determination using Pipeline Pilot, Dassault Systems.

#### Generating drug-perturbed RNA-seq profiles and protein activity (PLATE-Seq)

Cells were seeded onto two 384-well tissue culture plates per cell line, as for DRC generation. The next morning, compounds were added using the Echo 550 system. Each drug was dispensed at the smaller of its previously determined EC_20_ concentration or its calculated C_max_ (maximum tolerated serum concentration), but never above 10 μM in order to avoid confounding factors from activation of cell death and drug/stress response pathways^22,81,82^. The Cmax was used to avoid perturbing cells with non-physiologically relevant drug concentrations. For each cell line, two replicates were done with each plate including 372 distinct drug treatments, 6 DMSO vehicle controls, and 6 untreated samples. After 24 hours, plates were spun down at 295 × g for 1 minute. Media was removed and cells were re-suspended in 30 μL of Turbo Capture Lysis (TCL) buffer (Qiagen) containing 1% beta-mercaptoethanol (BME). Finally, multiplexed, low depth (< 5M reads) RNA-seq profiles at 24h following treatment of each cell line were generated for each drug, DMSO vehicle, and untreated control, using the PLATE-seq microfluidic automation platform^46^ The screen was performed by the Columbia JP Sulzberger Genome Center High-Throughput Screening facility of the Herbert Irving Comprehensive Cancer Center (HICCC).

RNA-seq reads were mapped to the human reference genome version GRCh38 using Kallisto, and individual drug gene counts demultiplexed by the barcoded reads. Expression data were normalized by equi-variance transformation, based on the negative binomial distribution with the DESeq R package (Bioconductor). To correct for potential cross-plate batch effects across the two replicates, we used the function *combat* from the sva package in R to compute the batch-corrected normalized gene expression and to fit a linear model for each compound against the six plate-matched DMSO controls, included to minimize the need for cross-plate normalization. We used the *limma* package for R to fit the linear model and to compute p-values and moderated t-statistics for each gene. For each compound, we used a vector of these statistics to generate a gene expression profile of Z-scores representing the differential compound effect between drug and DMSO vehicle control. Differential protein activity for each drug in the DIPG6 and SF8628 drug perturbation data was then computed by VIPER-inferred activity of the drug vs. DMSO control gene expression signature using the DMG bulk network.

#### Bulk OncoTarget and OncoTreat to define DMG pharmacotypes

We used the OncoTreat algorithm and the drug-perturbed differential protein activity in the DMG cell lines to predict drugs that would invert the activity of the top and bottom 50 MRs driving tumorigenesis (caudate signature) in each of the 122 DMG patient bulk protein activity signatures. Specifically, we evaluated enrichment of the top/bottom 50 sample-specific MRs in the drug-induced differential protein activity signature for each drug in each cell line by aREA. We then integrated the NES across the two cell lines by Stouffer integration weighted by the sample’s OncoMatch score to each cell line, ensuring that the OncoTreat score for drugs tested in the higher fidelity model for each sample would receive more weight for that sample (Weighted OncoTreat). This produced a ranked list of OncoTreat-predicted drugs for each DMG sample based on the strength of MR reversal for that sample. OncoTreat generated drug NES scores were converted to p-values using a one-tail p-norm function with correction for multiple hypothesis testing across the drugs in a sample accounted for by the Bonferroni method. Drugs with a Bonferroni p ≤ 10^-5^ are considered significant based on prior OncoTreat *in vivo* validation data^22,25^.

We then defined DMG patient pharmacotypes (groups of patients predicted to respond to the same drugs) by performing unsupervised clustering across the 122 DMG patients by OncoTreat scores using the optimalCluster function in the N1platform. This function generates an optimal cluster structure based on cluster reliability score analysis as described in^25^ We selected the top 20 OncoTreat-predicted drugs for each cluster by evaluating both their overall effects across the patients in that cluster by Stouffer integration of OncoTreat NES (global drug score) and their cluster-specific rank relative to its rank in other clusters (cluster-specific drug score). This prioritizes both drugs that are highly active across patients as well as drugs that might have a specific unique but strong activity in a specific pharmacotype. A heatmap representing the - log10(Bonferroni p-value) was generated with patients ordered by their pharmacotype and drugs ordered by their activity across patients.

Additionally, we employed OncoTarget to subset the VIPER outputted protein activity measurements across the 122 DMG patients (GTEx caudate signature) using a previously defined curated list of 180 actionable regulatory proteins representing validated targets of high-affinity pharmacological inhibitors, either FDA approved or in clinical trials ^25^. The final OncoTarget results included the VIPER-inferred NES scores across the 122 patients for these 180 druggable proteins with significance assigned by a one-tail p-norm function, adjusting for multiple hypothesis testing across the proteins for each sample by the Bonferroni method (Bonferroni p ≤ 10^-5^ considered significant). For the bulk OncoTarget heatmap representing the -log10(Bonferroni p-value) for each targetable regulator across the patients, we included all proteins that were significant in at least one patient, ordering patients by the OncoTreat-defined pharmacotypes.

#### DMG patient single-cell RNA-seq dataset

Publicly available scRNA-seq across six DMG patient tumor biopsies, specifically raw FASTQ files, were obtained from^51^ and aligned to the GRCh38 reference genome by Kallisto to derive a DMG single-cell count matrix. As a pre-processing step, low quality cells and genes without reads were removed from the analysis. Specifically, cells with fewer than 1000 or more than 100,000 total reads or mitochondrial transcript fraction greater than 15% were removed. Filtered data were then normalized to log10(counts per million + 1). A gene expression signature was generated from the normalized data using the Seurat SCTransform scaling function, and subsequently integrated using the Seurat AnchorIntegration function. The resulting data were projected into their first 5 principal components using the RunPCA function, and further reduced into a 2-dimensional visualization space using the RunUMAP function. Meta-cells were further computed by summing read-counts of each cell with its 10 nearest neighbor cells, then re-normalizing to log10(counts per million + 1) and re-scaling by SCTransform.

| Patient | BCH1126 | BCH836 | BCH869 | MUV1 | MUV10 | MUV5 |
| --- | --- | --- | --- | --- | --- | --- |
| # Cells before quality control | 384 | 864 | 960 | 624 | 960 | 1056 |
| # Cells after quality control | 265 | 514 | 747 | 289 | 411 | 806 |

#### Resolution-optimized Louvain clustering

Unsupervised clustering of single cells from the 6 DMG patients was performed in two steps. The Louvain algorithm (as implemented in Seurat) uses the FindNeighbors and FindClusters function. However, the FindClusters function includes a resolution parameter that allows selection of a higher or lower number of clusters depending on whether the parameter is increased or decreased, respectively. As such, over-clustering or investigator evaluation of ideal clustering purity is subjective. To address this, we optimized the resolution parameter based on average silhouette score (SS) width, an objective measure of the validity of a clustering solution^83^. Specifically, clustering was performed with resolution parameters ranging from 0.01 to 1.0 at intervals of 0.01, and cluster quality was evaluated by the average SS at each resolution value to select an optimal parameter within this range as described in^83^. For each resolution value, the clustered cells were subsampled to 1,000, and SS was computed for these 1,000 cells and their cluster labels. This was repeated for 100 random samples of 1,000 cells to compute a mean and standard deviation of average SS at each resolution value (Extended Data **Fig. 4e**). The resolution parameter that maximized mean SS was the one selected as the optimal resolution at which to cluster the data without over-clustering. This clustering methodology was used on the gene expression signatures generated by Seurat as well as the subsequent protein activity signatures generated for the tumor cells described below.

#### Microglia signature

Enrichment of a microglia gene signature in the gene expression and protein activity profiles for the DMG patient single cells was performed by GSEA using the gsea function in the atools package in R with 100 permutations for p-value estimation. The microglia gene signature included the following genes—Sall1, Hexb, Fcrls, Gpr43, Cx3cr1, Tmem119, Trem2, P2ry12, Mertk, Pros1, Siglech—and was extracted from^52^.

#### Copy number inference (InferCNV) analysis

Copy number variation (CNV) was inferred from gene expression counts at the single cell level using the InferCNV R package. Cells belonging to clusters defined as immune based on enrichment of the microglia signature or CD33 activity as determined on the protein activity space were used as a CNV reference set, and cells belonging to other clusters were assessed for CNV. These remaining presumed tumor cell clusters were confirmed to exhibit significant copy number alterations in at least a subset of cells belonging to each cluster. Patient-specific analysis had identified heterogeneity among individual patients in type and degree of CNV across the single cells with certain samples exhibiting a low degree of copy number alterations, known to occur in subsets of DMG patients. Therefore, rather than setting a CNV cutoff for individual single cells, we instead assumed that given their transcriptional similarity as defined by belonging to the same cluster, any cells belonging to a cluster that exhibited significant CNVs in at least a subset of cells were likely transformed as well.

#### DMG single cell regulatory networks

Networks were generated using the log2(CPM/10+1) normalized gene expression counts for tumor cells as defined in^53^ for each of the six patients individually using the ARACNe-AP algorithm. Putative regulatory proteins were defined as for the bulk network. Standard parameters were set including 0 DPI (Data Processing Inequality) tolerance and MI p-value threshold of 10^-8^. This resulted in 1,440 regulators, 5,118 targets and 72,000 interactions for BCH1126; 2,469 regulators, 7,088 targets and 123,450 interactions for BCH836; 2,045 regulators, 6,343 targets and 102,250 interactions for BCH869; 1,117 regulators, 4,558 targets and 55,850 interactions for MUV1; 714 regulators, 4,505 targets and 35,700 interactions for MUV10; and 2,281 regulators, 7,007 targets and 114,050 interactions for MUV5.

#### DMG single-cell protein activity inference

Single cell protein activity was computed on an internally scaled gene expression signature (generated by the Seurat SCTransform function) across meta-cells by the metaVIPER function in the VIPER package (Bioconductor) using the six patient-specific single cell ARACNe networks. Initially, metaVIPER was developed as an adaptation of VIPER to single-cell data^36^ interrogating multiple potential networks to accurately recapitulate protein activity in populations for which no known context-specific network was available. Briefly, protein activity is inferred for a given gene expression signature using multiple networks which are integrated on a protein-by-protein basis using the square of the NES generated by each individual network. Since a non-relevant network would generate a protein activity NES close to zero under the null model, networks that generate more extreme NES’s can be interpreted to more accurately match the given biological context and are thus weighted more heavily. We therefore generated context-matched patient-specific ARACNe networks for each patient and use these networks for optimal inference of protein activity.

#### Single cell pathway enrichment analysis

Using the clusters defined by Louvain resolution-optimized clustering on the protein activity space for the transformed tumor cells across the six DMG patient single cell samples, differential gene expression analysis was performed by the Wilcox Test comparing the cells of the cluster of interest to the cells of all other clusters in order to get the differentially upregulated genes in each cluster individually compared to the rest. Then, all the statistically significant (Bonferroni-adjusted p-value <0.05) genes that had positive log2FC values (positively upregulated) were taken as the set of significantly upregulated genes corresponding to each of the seven human DMG subpopulations (gene set). These genes were the input for the enrichR function in R as part of the enrichR package, and the gene set library ‘MSigDB Hallmark 2020’ was selected. Pathway enrichment based on an overrepresentation analysis was then done with multiple hypothesis correction testing across pathways within each population performed by the FDR method with FDR ≤ 0.05 considered significant.

#### Single cell OncoTarget and OncoTreat

To identify aberrantly activated, actionable proteins in each of the seven transformed human subpopulations (OncoTarget), we subset the single-cell VIPER matrices for the 180 directly druggable proteins previously described. Protein activity was computed by VIPER via generation of synthetic bulk samples from each subpopulation and interrogation of the bulk DMG network with the differential gene expression signature of the synthetic bulk samples relative to the centroid of the GTEx caudate; We thus produced a list of NES scores across the 180 druggable proteins for each of the seven subpopulations of interest. NES scores were converted to adjusted p-values by the p-norm function (one-tail) using the Bonferroni method for multiple hypothesis correction within each subpopulation. Proteins found to be significantly active (Bonferroni p ≤ 10^-5^) in at least one of the populations were considered. The proteins most significantly active across the populations of interest as well as those uniquely active within populations were plotted in the heatmap.

Single cell Oncotreat was performed for the VIPER protein activity signature described above to assess the statistical significance for each drug’s ability to invert the activity of the top and bottom 50 MR proteins in each of the seven subpopulations. This is based on the enrichment of subpopulation MR proteins in proteins differentially active in drug vs. DMSO-treated DIPG6 and SF8628 cells (PLATE-Seq), with negative NES values corresponding to MR activity inversion. NES scores across the two cell lines were again integrated by Stouffer integration weighted by the OncoMatch score representing enrichment of that subpopulation’s MRs in the respective cell line’s protein activity signature.

For both OncoTarget and OncoTreat analysis relative to GTEx caudate, synthetic bulk profiles for each subpopulation were generated by summing read counts of each gene across all the cells belonging to a specific cluster, then re-normalizing to log10(CPM + 1), and scaling to a z-score by gene using the mean and standard deviation of log10(CPM + 1) across the entire set of GTEx caudate samples.

#### Single-cell isolation and Gel Beads-in-emulsion (GEM) generation in DMG cell lines

Cells were detached with room temperature ACCUTASE^TM^ (Sigma-Aldrich #A6964) for SF8628. For neurospheres, cells were collected and spun down at 250 x g for 6 minutes at 4°C. Supernatant was removed and cells were resuspended in 1 ml of room temperature ACCUTASE^TM^ for 3 min and pipetted ∼20 times to dissociate spheres. Then, cells were filtered through a 70μm strainer to remove clumps. Neurospheres and attached cells were resuspended in scRNA-seq buffer (HBBS + 0.04% BSA, Fisher Scientific #14-175-095 and Miltenyi Biotec #130-091-376, respectively) and washed 3 times. On the last wash, cells were examined for cell viability and counts using the Countess II and resuspended in 1×10^6^cells/ml. Dissociated cells were processed for single-cell gene expression capture (scRNA-seq) using the 10X Chromium 3’ Library and Gel Bead Kit (10x Genomics), following the manufacturer’s user guide. Libraries were prepared following the manufacturer’s user guide and sequenced on the Illumina NovaSeq 6000 Sequencing System at the Columbia University Single Cell Analysis Core.

#### Drug dilutions

All drugs used in this study were received in powder form and treated with the same methods for dilution. Vehicle was made in quantities of 50 ml. The vehicle was comprised of 30% (15 ml) polyethylene glycol 400 (PEG 400, Selleck Chemicals #S6705), 5% (2.5 ml) polyoxyethylene sorbitan monooleate (Tween 80, Selleck Chemicals #S6702), and 65% (32.5 ml) dextrose 5% in water (D5W, Fisher Scientific NC0215480) (PTD). Total vehicle volume for dosing was prepared for the number of days of treatment and the number of mice receiving treatment. The number of days of treatment was selected to be a total of 10 days of dosing, Monday through Friday, before preparing more. Drug dose amounts (mg/kg) were determined in accordance with Section V. Drug, Reagent, Compound Administration under protocol number AC-AABL6551 (Y2 M12). Drug concentrations calculated were done with the assumption that each mouse averaged a weight of 30g and would be dosed at a total volume of 10 µl/g. After calculating the volume and drug aliquot weight, 200 µl of *N*-methyl pyrrolidinone (NMP, Sigma-Aldrich M79603) was added to increase the solubility of the drugs. The remaining volume needed was filled with the vehicle (PTD). The solution was vortexed, or probe sonicated, as needed, to ensure homogeny.

#### Generation of DIPG17 cell line-derived murine models and treatment arms

All mice were maintained under barrier conditions and experiments were conducted under the protocol approved by the Columbia University Institute of Comparative Medicine (ICM) and the Institutional Animal Care and Use Committee (IACUC) under protocol AC-AABL655. Juvenile male and female NOD.Cg Prkdcscid/J (NSG^TM^, Envigo) mice were inoculated with 500,000 DIPG17-luciferase tagged cells in 1:1 suspension (200 μl total) with full media and Matrigel^TM^ (Fisher Scientific CB-40234) via subcutaneous injection in the left flanks. Two weeks post-injection, animals were injected intraperitoneally with luciferin and scanned with PerkinElmer IVIS Optical Imaging to monitor tumor growth. Animals were monitored biweekly until tumor size reached ∼100 mm^3^ (range of volumes 75 – 185 mm^3^) and tumor-bearing mice were randomly assigned to distinct drug arms or vehicle control treated groups when they reached adequate size. Animals were treated for 5 days at the following doses: Vehicle (n=4) (N-Methyl-2-pyrrolidone (NMP) as solvent and 30%PEG-400 + 5% Tween 80 + 65% D5W (PTD) as vehicle, all drugs were dissolved in the same solvent and vehicle), Larotrectinib 200mg/kg PO (*per os,* oral gavage) QD (every day), Trametinib (n=2) 1mg/kg PO (5 days on, 2 off), Dinaciclib (n=2) 40mg/kg (Monday-Wednesday-Friday), Avapritinib (n=3) 30mg/kg PO QD (5 days on, 2 days off), Ruxolitinib (n=2) 90mg/kg PO BID (180mg/kg/day), Venetoclax (n=3) 100 mg/kg PO QD (5 days on, 2 days off), Etoposide (n=3) 8mg/kg IP QD (5 days on, 2 days off), Mocetinostat (n=2) 50mg/kg PO QD (5 days on, 2 days off) and Napabucasin (n=3) 10mg/kg (Monday-Wednesday-Friday). All drugs were ordered from Selleckchem. Mice were euthanized for tumor isolation and scRNA-seq 2 hours after the last dose was given at day 5.

#### IVIS Spectrum Optical Imaging

Before imaging, mice were injected intraperitoneally with 10 μl/g luciferin (15mg/ml). Mice were then anesthetized with isoflurane at 2L/min and were imaged 10 minutes post-injection with the IVIS Spectrum Optical Imaging System (Caliper).

#### High Frequency Ultrasound Imaging System

3D-US imaging data sets were collected for each xenograft using the Visulasonics VEVO 3100 High Frequency micro-Ultrasound System (VisualSonics Inc.) designed for small animal imaging. For imaging acquisition, mice were anesthetized with 1.5-2% isoflurane in oxygen delivered at 0.75 L/min and were immobilized to be imaged. Xenografts were coated with warmed (37C) Aquasonic 100 US gel (Parker Laboratories) and centered in the imaging plane. Three-dimensional data were acquired along the entire length of the xenograft. For imaging analysis, images were imported into the VEVO software and tumor volume was calculated.

#### Assessment of tumor volume changes in the DIPG17 cell line-derived subQ murine model drug studies

Volumes were measured in the VEVO software. Tumor volume percentage changes were calculated with the formula shown below. Statistical significance for difference in tumor volume changes between each drug treatment arm and the vehicle control was computed by the Welch’s t-test with p ≤ 0.05 considered significant.

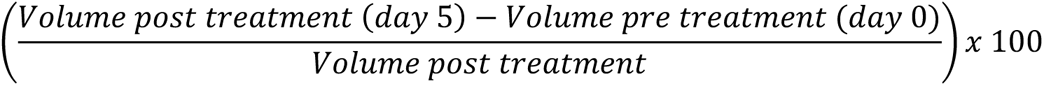

#### Single cell RNA-seq of DIPG17 cell line-derived mouse models including tumor dissociation, single cell isolation and Gel bead-in-EMulsion (GEM) generation

After tumor isolation, fat, fibrous and necrotic areas were removed from the tumor sample in a tissue culture hood. Then, tumor was cut into 1-2 mm pieces and transferred to a gentleMACS C Tube (Miltenyi Biotec #130-093-237) containing 4.7 mL of sterile media (Knockout DMEM/F12), 200 μl of Enzyme H, 100 μl of Enzyme R, and 25 μl of Enzyme A from the human Tumor Dissociation Kit (Miltenyi Biotec #130-095-929). C Tubes were placed in the gentleMACS Dissociator under the “Medium” Tumor type (gentleMACS Program 37C_h_TDK_2) for 18 minutes. Then, samples were filtered through a 70μm strainer (previously primed with 2 ml of sterile media). Samples were centrifuged at 250 *x g* at 4°C for 8 minutes. Then, resuspended in 400 μl of red blood cell (RBC) lysis buffer (Miltenyi Biotec #130-094-183) and incubated on ice. After 2 minutes, at least 3 times the volume of RBC was added to rebalance osmolarity and the sample was centrifuged at 250 *x g* at 4°C for 5 minutes and resuspended in HBBS + 0.04% BSA. Samples were then examined for cell viability and counts using the Countess II. Samples were then brough to the Columbia University Single Cell Analysis Core where libraries were prepared following the manufacturer’s user guide and sequenced on the Illumina NovaSeq 6000 Sequencing System. ScRNA-seq data were processed with Cell Ranger software’s default parameters. Cell Ranger performed default filtering for quality control, and produced a barcodes.tsv, genes.tsv, and matrix.mts file containing transcript counts for each sample. These data were loaded into the R version 4.2.2. for further analyses.

### Drug-perturbation analysis of a DIPG17 subQ mouse models

#### Quality control, normalization, and scaling

As a pre-processing step, low quality cells and genes without reads were removed from the analysis. Quality control was based on read depth and mitochondrial gene percentage. Samples with too many or too few reads are likely sequencing errors (doublets or empty oil droplets during the generation of single cells), while a high mitochondrial gene percentage is indicative of high cell stress, damage or dying cells. Cells with fewer than 1000 or more than 100,000 total UMIs or mitochondrial transcript fraction greater than 15% are removed in quality-control filtering. Filtered data are then normalized to log10(CPM + 1) and scaled using the Seurat SCTransform scaling function. Gene expression signatures were generated for all the single cells in the vehicle and drug treatment arms on a gene-by-gene basis by scaling to the centroid of expression across the vehicle cells (subtracting the mean expression across the vehicles and dividing by the sd).

#### Human-derived cell selection

The transformed, human-derived nature of the profiled cells was confirmed by consensus alignment of RNA reads to the human versus mouse reference genomes (Extended Data **Fig. 5c**). We then modeled a bimodal distribution by a finite mixture model that uses continuous distributions from the exponential family^84^. The parameters of the finite mixture model were estimated by the Expectation-Maximization (E-M) algorithm^85^. We used the em function in R, which included two parameters (a location parameter μ and a scale parameter σ) for each of the two probability distributions and a mixture parameter. The density of the bimodal distribution of concentrations thus reads:

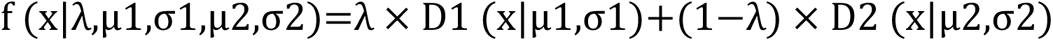

In our dataset of 24 samples, estimates for μ1 and σ1 (the location and scale parameters of the peak of the lower concentrations, which would be the mouse-infiltrated cells) were 24.9881 and 4.3501, respectively. For the highest concentrations (human-derived cells), the μ2 and σ2 were 80.2255 and 10.0286, respectively. We then used the fitted finite mixture model to identify a cutoff value that discriminated the two modes of the dataset (using the cutoff function in R). Additionally, we computed the probability for a datum to belong to distribution *D*1 (lower concentrations) or *D*2 (higher concentrations) at a given cutoff.

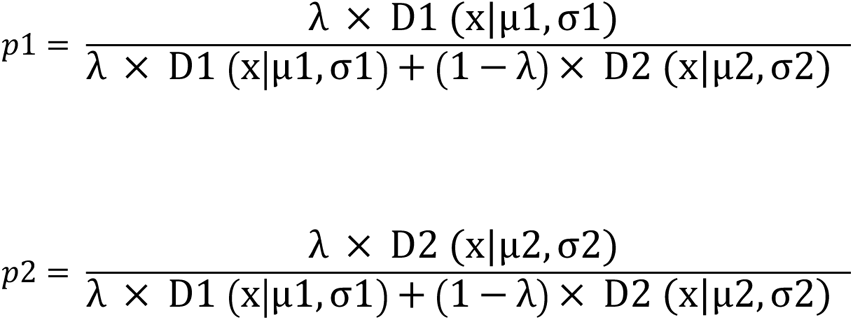

Cutoff was set at 43.3241%, with the probability of a datum chosen at random from *D*1 being more than 43.3241% equaling 0.000017 (Extended Data **Fig. 5c**). Given this, we selected cells with a Human UMI % of >50% as human tumor cells for further downstream analyses, knowing that the probability of a cell chosen at random with a UMI % >50% belonging to a non-tumor mouse cell (*D1* distribution) was virtually 0.

#### Protein activity inference by VIPER

Given that we were analyzing DMG human tumor single-cell data, the patient-specific single-cell regulatory networks generated from the six DMG patient tumors previously described were interrogated by metaVIPER to generate protein activity profiles for the subQ tumor model scRNA-seq. metaVIPER was run with gene expression signatures generated relative to the centroid of vehicle cells on a cell-by-cell basis producing protein activity scores (NES) for 2,821 regulatory proteins across the 95,687 tumor single cells across all the treatment conditions.

#### Assignment of a cell identity

To assign an individual cell identity to each of the 11,325 DMG cell line and the 95,687 DMG subQ tumor single cells profiled, we computed GSEA evaluating enrichment of the top 50 most differentially activated proteins (MRs) in each of the seven DMG patient tumor subpopulations as the gene set (Extended Data **Table 4**) in the protein activity signature of the model single cells on a cell-by-cell basis with the NES scores determined by 100 permutations. Then, p-values were calculated across the single cells for each subpopulation using 1 – pnorm(NES), and q-values were determined in order to incorporate multiple hypothesis testing correction by FDR. Based on the close similarity of OPC and OPCQ as well as OC and OPC/OC states, each cell was assigned an identity (OPC, OPCC, OC, AC, or OPC/AC) based on its most significant score. Specifically, those with significant enrichment in Clusters 0 (OPCQ) and 1 (OPC) MRs were labeled as OPC, Cluster 2 as AC, Cluster 3 as OPCC, Clusters 4 (OC) and 5 (OPC/OC) as OC, and Cluster 6 as OPC/AC. Only cells with an FDR-corrected p-value ≤ 0.05 were assigned a cell type.

#### Evaluation of drug-induced subpopulation-specific depletion

For each drug treatment arm, within each assigned cell identity, we assessed the log-fold change in proportion of cells belonging to that population in the drug treated versus vehicle tumors. We assessed each drug-treated replicate independently and calculated the mean log-fold change and standard error across the replicates. Given that OPC cell states—comprised of OPCQ, OPC, and OPCC cells—OC cell states—comprised of OC and OPC/OC cells—and AC cell states—comprised of AC and OPC/AC cells—shared similar drug predictions within them, we plotted the mean log-fold change with error bars representing the standard error as bar plots across the following three groups to represent whether drug effects were in line with the predictions: OPC (OPCQ, OPC, OPCC), OC (OC and OPC/OC), and AC (AC and OPC/AC).

We additionally produced bar plots for each drug arm depicting the mean and standard error of proportions of cells belonging to each of the following populations—OPC (OPCQ + OPC), OPCC, OC (OC + OPC/OC), AC, OPC/AC, and NS (non-significant)—in the drug-treated (n = 2-3) versus vehicle control treated (n = 4) samples. Standard error was computed as Standard Deviation ((SD)/√Number of samples and reflected in the error bars (Extended Data **Fig. 6**). A chi-square test (chisq.test function in R) was used to assess whether the distribution of proportion of cells belonging to each population was significantly different between the drug-treated cells (done separately for each treatment arm) versus the vehicle control-treated cells (Extended Data **Table 5**). This was done for all cells aggregated across the replicates in each treatment arm. We computed a chi-square p-value for each individual cell type versus the rest and adjusted the p-values for multiple hypothesis testing both across the six populations evaluated and the nine treatment arms using the FDR method.

#### Pharmacodynamic assessment of MR activity reversal *in vivo*

Pharmacodynamic assessment of MR activity reversal was assessed after 5 days of treatment, and 2 hours after the last dose on day 5. metaVIPER and the six DMG patient single cell networks were used to generate a differential protein activity signature (by z-score method for gene expression signature generation) for each single cell belonging to a drug-treated sample versus the four vehicle control treated samples for each of the cell types (OPC, OPCC, OC, AC, and OPC/AC) independently. Then, the signatures within each drug treatment arm were integrated by Stouffer’s method and a unique vector of drug vs. vehicle differential protein activity for each subpopulation was generated from the *in vivo* perturbations. The enrichment of DMG patient tumor-specific top and bottom 50 activated and inactivated candidate MRs in each subpopulation from the human single cell analysis in the drug vs. vehicle protein activity signature of the respective subpopulation in the model was assessed by aREA which provided a measure of MR-inversion. For drugs predicted by OncoTarget, NES activity of the respective MR was assessed in each population as a measure of on-target activity by that drug in each population.

#### Intracranial implantation of DIPG4423 cells to generate a syngeneic orthotopic DMG murine model

All mice were maintained under barrier conditions and experiments were conducted under the protocol approved by the Columbia University Institute of Comparative Medicine (ICM) and the Institutional Animal Care and Use Committee (IACUC) under protocol AC-AABL655. First, cells were detached using ACCUTASE^TM^ and centrifuged at 250g x 6 minutes. The pellet was then resuspended in 1 ml of room temperature ACCUTASE^TM^ for 3 minutes and pipetted up and down approximately 20 times. Then 3 ml of fresh media was added, and cells were resuspended and centrifuged again at 250g x 6 minutes. Cells were filtered through a 70μm cell strainer to remove any clumps. Cells were resuspended in a volume of 100,000 cells/μl and cell viability was assessed using the Countess 3FL Automated Cell Counter. All intracranial injections were conducted under sterile conditions. Before intracranial implantation, 6-week-old B6(Cg)-Tyrc-2J/J mice from Jackson laboratories were anesthetized with 1-2% isoflurane anesthesia in oxygen delivered at 0.75 L/min and were immobilized in a mouse stereotactic instrument (Stoelting). A 1-cm incision was made at the midline of the scalp to expose the sagittal suture and lambda of the skull. A burr hole 1 mm in diameter was made in the skull at a position 1.5 mm posterior to the lambda and 1.5 mm lateral to the sagittal suture. A Hamilton syringe containing 100,000 cells suspended in 1 μl full media was inserted 5 mm below the skull surface and injected at a rate of 0.1 mL/min over 10 minutes. Two minutes following implantation, the syringe was slowly removed from the mouse brain in 1 mm increments spaced 30 seconds apart. Then, the burr whole was covered with bone wax (Fisher Scientific #50-118-0260) and the incision was closed with three to four sutures (Fisher Scientific #50-118-0846). After suturing, triple antibiotic ointment was applied to the wound and animals were injected (IP) with 0.05mg/kg of buprenorphine (Covetrus, approved by DOH/DEA). Mice were put on a heating pad until they woke up from anesthesia and then were placed back in the cage. Wounds and sutures were thoroughly monitored every day for 7 consecutive days to ensure complete recovery. Tumor presence and location were confirmed 12 days post-injection with MRI.

#### Assessing model fidelity of the DIPG4423 syngeneic orthotopic DMG murine model

To assess whether we could expect similar drug-induced subpopulation-specific effects in this model to test our hypothesis that combining drugs targeting distinct tumor subpopulations would improve treatment response, it was critical to first validate the presence of all of the DMG patient tumor subpopulations within these orthotopic model tumors. In order to do so, we injected B6 mice and treated them with vehicle control (NMP) for 3 weeks, then tumors were isolated, dissociated into single cells and sequenced. We performed scRNA-seq on two vehicle controls and one normal brainstem, which yielded 20,362 and 2,978 high-quality single cells, respectively (after quality control). First, unsupervised clustering based on gene expression and UMAP projection with singleR annotation was performed using the standard Seurat pipeline. Just like in the six patient scRNA-seq samples, singleR (using mouse.rnaseq as a reference) mostly divided these 3 samples into glial, neuronal and immune system-related cells (with significant microglial presence). Then, we removed the immune system-related clusters and focused on re-clustering of putative tumor cells (clusters 0, 4, 5 and 9) (Extended Data **Fig. 8c**). After re-clustering of the glial and neuronal cells, the transformed, cancerous nature of cells in the two brainstem tumor samples was confirmed by inferCNV analysis using the normal brainstem cells (WT) from each cluster as a reference (Extended Data **Fig. 8d**). Clusters 0, 1, 4, 5 and 7 from the new unsupervised clustering were selected as malignant based on significantly aberrant copy number variations. This yielded 8,013 malignant cells for further downstream analyses (Extended Data **Fig. 8e**).

The gene expression data from the 8,013 tumor cells were then transformed into a gene expression signature by scaling each gene within each cell to the mean/sd across all tumor cells (internal signature). Given that regulatory networks can be species specific, we generated two murine DMG single cell networks by the standard application of ARACNe-AP to the normalized gene expression data from brainstem tumors #1 and #2 (one network per sample). We then inferred protein activity on the internal gene expression signature by interrogating these murine networks via metaVIPER. Unsupervised clustering of the tumor cells by protein activity by resolution-optimized Louvain clustering as done for the human and subQ model samples identified five clusters that segregated remarkably well with the DMG patient subpopulation cell type assignment as computed by enrichment analysis of the DMG patient subpopulation top 50 MRs in the DMG model single cell protein activity signatures described previously (**Fig. 6d**, Extended Data **Fig. 8f**). This supports that the tumor heterogeneity and population structure in these models is reflective of what is observed in patients, and thus these are good models to further validate our drug predictions orthotopically *in vivo*. We further validated conservation of the subpopulation-specific OncoTarget and OncoTreat predictions in this model by generating synthetic bulk samples from cells assigned to each population (sum of UMIs across the cells) and running OncoTarget/OncoTreat as for the DMG patient single cell analysis on the protein activity signature of each synthetic bulk sample relative to the centroid of the GTEx caudate samples.

#### Magnetic resonance imaging (MRI) and Image Analysis

A 9.4T MRI system (Bruker Medical, Boston, MA, USA) was utilized for verification of tumor location and tumor presence, as well as to monitor tumor growth and calculate tumor volumes. Mice received 1-2% isoflurane anesthesia in oxygen delivered at 0.75 L/min and were placed into a birdcage coil (diameter 35 mm). To validate tumor growth, MR images were obtained using a T2-weighted TurboRARE sequence. Images were imported in DICOM files and the free, open-source platform 3D Slicer (www.slicer.org) was used to measure tumor volume.

#### Brain fixation and Hematoxylin and Eosin (H&E) staining

Three weeks after injection (for vehicles and/or untreated) and until they showed neurological symptoms (for treated) mice were euthanized, whole brains were isolated and fixed in 15ml of formaldehyde overnight. After that, a sagittal cut was performed, and brains continued fixation at room temperature. After 12 hours, formaldehyde was replaced with cold saline (1X PBS) and stored at 4 degrees overnight. The following day, brains were brought to the Histology Core at Columbia University Irving Medical Center for paraffin embedding, H&E staining and full slide scans. Scans (.svs) were analyzed in Orbit Image Analysis Software Version 3.64.

#### Single-cell dissociation in orthotopic models

Three weeks after injection, untreated mice were euthanized for brainstem tumor single-cell isolation and Next-generation sequencing. Before euthanasia, mice were anesthetized with 2% isofluorane and perfused for 8 minutes with cold saline (1X PBS) and/or until the liver turned pale. Upon dissection of the brainstem, the tissue was placed in isolation media consisting of RPMI 1640 with GlutaMAX Supplement and HEPES (Gibco #72-400-120) with 0.01% 50X B27 supplement (Gibco #17-504-044). The tissue was then homogenized in a 15ml glass tissue grinder, filtered through a 70μm filter and spun down at 300g x 10 minutes at 4 degrees. The pellet was resuspended in 1ml of isolation media (per brainstem), 100ul of myelin removal beads were added (Miltenyi Biotec #130-096-733) and the mixture was incubated for 15 minutes. After one wash, the solution was then passed through LS positive separation columns (Miltenyi Biotec #130-042-401) to deplete the myelin. After another wash with single-cell buffer (HBBS + 0.04% BSA), the pellet was resuspended in 400 ul of RBC lysis buffer and incubated on ice for 2 minutes. Then, 1ml of single-cell buffer was added and sample was spun down at 250g x 5 minutes. Three more washes with single-cell buffer were performed and the final pellet was resuspended in the appropriate volume of single-cell buffer to have an approximate concentration of 1×10^6^ cells/mL. Cell counting and viability was assessed with Countess 3FL Automated Cell Counter.

#### Syngeneic orthotopic model survival studies

Two weeks post stereotactic injection and after confirming presence of tumor via MRI, mice were randomly enrolled into one of the treatment arms (monotherapy or combination therapy). All drugs were given 5 days on, 2 days off, to minimize drug toxicity. Monotherapy treatment arms were as described before. Combinatorial therapies were based on agents targeting different subpopulations and were as follows: avapritinib in combination with either larotrectinib 200mg/kg PO, venetoclax 100mg/kg, or ruxolitinib 180mg/kg/day (90 mg/kg PO BID); trametinib in combination with either venetoclax 100mg/kg or ruxolitinib 180mg/kg/day (90 mg/kg PO BID); and dinaciclib in combination with venetoclax 100mg/kg or ruxolitinib 180mg/kg/day (90 mg/kg PO BID). Toxicity was monitored by factors such as weight changes, body temperature, behavioral changes as per the protocol. dinaciclib+venetoclax was not well tolerated, and this treatment arm was discontinued. Drugs were always given with at least a 6-hour window between them to avoid as much animal distress as possible. Given ruxolitinib’s dosing regimen (BID) and short elimination half-life (3 hours), combinatorial therapy with ruxolitinib and an OPC targeting agent (avapritinib, trametinib or dinaciclib) was administrated with a 3-hour window in between the morning and the evening 90mg/kg ruxolitinib doses.

Mice were euthanized, according to protocol, as soon as they exhibited any significant neurological symptoms or weigh loss more than 15% from the initial day of treatment. Survival proportion graphs were generated using GraphPad Prism Version 9.5.1. and p-values for differences in survival between treatment arms was calculated by Log-rank (Mantel-Cox) test.

#### Synergy studies

Dose response curves were generated by plating cells at appropriate density of 1,000 cells per well for DIPG36 and DIPG4423, and 500 cells per well for SF8628 into 384 well white tissue culture plates (Greiner 781080) and incubated for 24 hours at 37C 5% CO2. Compounds were added to plates using the HP Digital Dispensing system, with each compound titrated from 40 mM to 2.6 nM, in triplicate, normalized in DMSO (0.4%). After 48 hours each plate was treated with 25 mL’s of CellTiter Glo (Promega Corp G7572) and read in enhanced luminescence mode on an EnVision Multi-Label plate reader (Revvity Corp.). Data was normalized utilizing on plate controls (DMSO only and Thimerosal positive control) using Pipeline Pilot Screening data tools.

For drug combinations, cells were plated as described above. After 24 hours, drug combinations were added via the HP digital dispensing system in 10 x 10 matrix with the previously determined IC50 used as the maximal concentration. The matrixes were run in triplicate and randomized with the appropriate controls on each plate (DMSO and Thimerosal positive control). After 48 hours, the plates were read as described above using CellTiter Glo. BLISS synergy scores were calculated using the synergyfinder.fimm.fi website (SynergyFinder 3.0.^86^) and Extended Data **Fig. 16a** was also generated using synergyfinder.

## Acknowledgements

We thank Dr. Michelle Monje for generously sharing the SU-DIPG-IV (DIPG4), SU-DIPG-VI (DIPG6), SU-DIPG-XVII (DIPG17) and SU-DIPG-XXXVI (DIPG36) cell lines. We also thank Dr. Rintaro Hashizume for sharing the SF8628 and SF7761 cell lines. We thank Dr. Oren Becher for sharing the DIPG4423 cell line. Lastly, we also thank the Columbia Genome Center, High-Throughput Screening, and Oncology Precision Therapeutics and Imaging Core (OPTIC) cores for their help.

This research was primarily funded by grants from Swim Across America and The Team Jack Foundation to J.P., as well as a Hyundai Hope on Wheels Impact Award to S.Z. and J.P., and the Gary and Yael Fegel Family Foundation. Additional funding support was provided by the Columbia University Irving Medical Center Herbert and Florence Irving Basic Science Scholar award and Pediatric Oncology Novel Research Fund Award awarded to J.P., as well as the Vagelos College of Physicians and Surgeons Interdisciplinary Research Initiatives Seed (IRIS) Fund. Further support was provided through the Postgraduate studies in North America and the Asia-Pacific region Fellowship by “La Caixa Foundation” and the Trainee Associate Member (TAM) predoctoral award from the Herbert Irving Comprehensive Cancer and Education Coordination centers at Columbia University, both awarded to E.C.F. T.T. was supported by fellowships from the German Research Foundation and ChadTough Defeat DIPG Foundation. R.D.G. was supported by the Hyundai Hope on Wheels Hope Scholar Award, Swim Across America, Rally Foundation, StacheStrong and the Musella Foundation. C-C.W. was supported by the Sebastian Strong Foundation, Swim Across America, the St. Baldrick’s Foundation Scholar Award, and the Pediatric Cancer Foundation. M.G. was partially supported by an ASCO Young Investigator Award as well as a grant from the Radiological Society of North America (RSNA). A.T.G. was supported, in part, by the Ruth L. Kirschstein National Research Service Award (NRSA) Institutional Research Training Grant (5T32GM007367-43), and in part by the NCI Ruth L. Kirschstein National Research Service Award Individual Fellowship (F30CA257765).

This work was further supported by the NCI Outstanding Investigator award R35 CA 197745, the NIH Shared Instrumentation Grants S10 OD012351 and S10 OD021764, and U01CA272610 (all to A.C.) and the NINDS Research Program Award R35 NS122339 (to R.W.-R). Additional grants included the NIH grant #P30 CA013696 (National Cancer Institute) and used the Oncology Precision Therapeutics and Imaging Core (OPTIC), and the NIH/NCI Cancer Center Support Grant P30CA013696 and used the Genomics and High Throughput Screening Shared Resource. This publication was supported by the National Center for Advancing Translational Sciences, National Institutes of Health, through Grant Number UL1TR001873. This study is also supported by NIH grants (R01 CA277605-01A1 and R01 NS132344-01) to Z.Z. The content is solely the responsibility of the authors and does not necessarily represent the official views of the NIH.

## Contributions

J.P., A.C., and E.C.F. designed the study and wrote the manuscript. S.Z. supported the conception of the study and assembly of the collaborative team at study onset. E.C.F. conducted and analyzed all experimental and animal work unless otherwise indicated and lead the computational analysis of the CRISPR/Cas9 KO screen data, the subcutaneous DMG murine model and human cell line scRNA-seq data, contributing to other computational analyses. L.T. analyzed the syngeneic orthotopic DMG model scRNA-seq, MR-inversion pharmacodynamic studies in the subQ model, managed the associated data repository, and shared expertise with other computational analyses performed by E.C.F. and J.P. J.W. and A.O. analyzed the DMG patient scRNA-seq data, including the single cell OncoTarget and OncoTreat studies. X.Z. conducted the experimental part of the CRISPR/Cas9 KO screen with supervision of Z.Z. L.V. procured, aligned, and contributed to the analysis of the DMG patient scRNA-seq data. P.L. provided expertise and contributed to the CRISPR/Cas9 KO screen analysis. A.T.G. provided expertise in the ARACNe3 algorithm and generated the DMG bulk network. The remaining bulk data analysis was conducted by J.P. who contributed to all the computational analyses conducted in this study. H.M., D.V.M., C.S., M.G., H-J.W. helped with animal work/husbandry and/or taught E.C.F. how to perform stereotactic injections and imaging. T.J.M., P.S.B., J.R.C., T.T. and R.W-R. generated the RNA-seq data for the three DMG patient biopsies and performed the subsequent drug screens. R.D.G. provided resources and guidance on animal work, which was conducted under her IACUC protocol. C-C.W. provided guidance with the syngeneic orthotopic mouse model. L.S., J.G. and S.Z. provided support with patient cohort assembly, clinical neuro-oncology expertise, and selection of clinically relevant drugs. All authors approved the final version of this manuscript.

## Extended Data Figure Legends

**Extended Data Figure 1.**
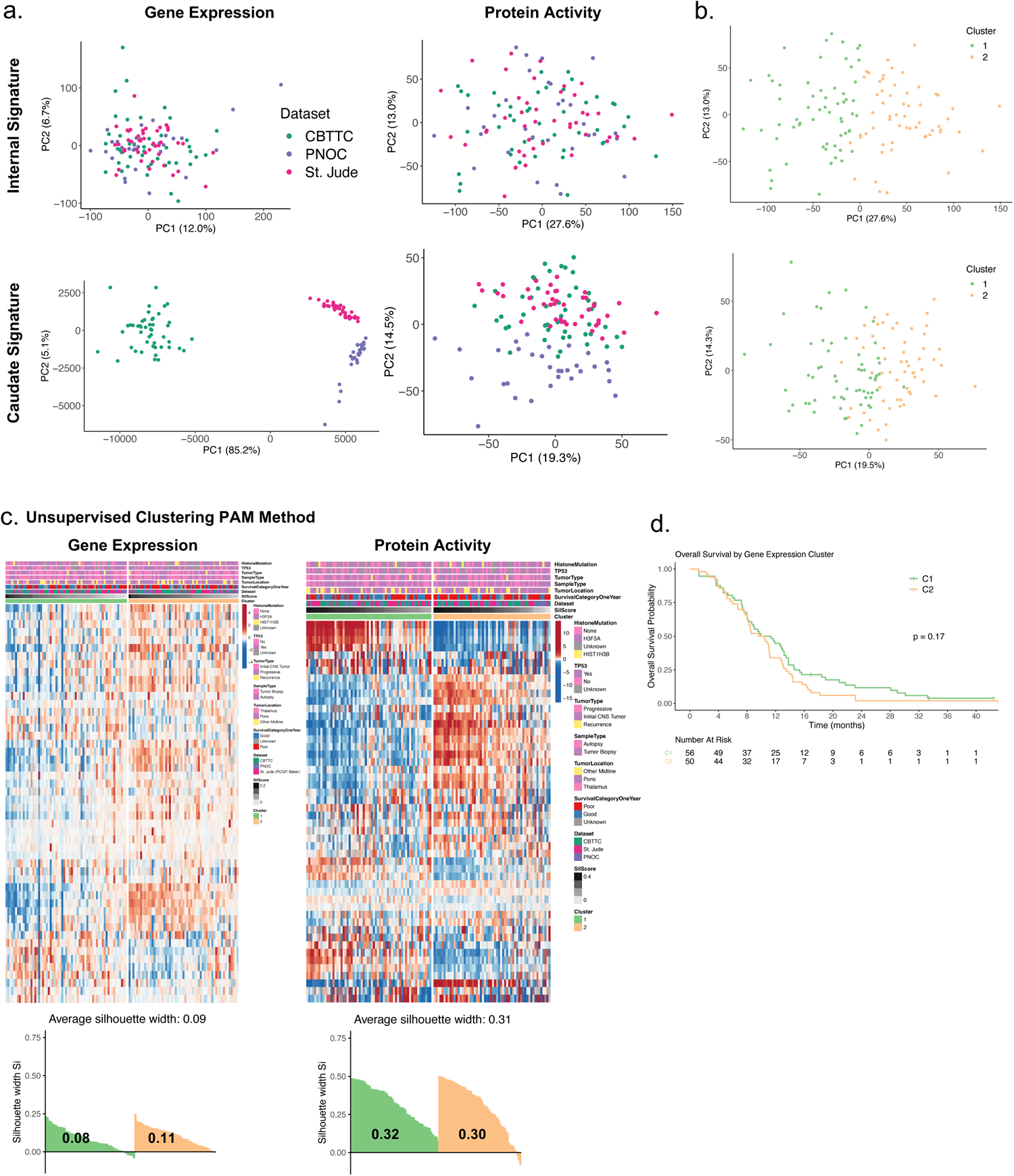
Gene expression versus protein activity analysis of 122 DMG bulk RNA-seq samples. **a.** PCA analysis of 122 DMG bulk RNA-seq patient sample gene expression (left) versus VIPER-inferred protein activity (right) signatures relative to an internal cohort-specific (top) vs. external normal GTEx caudate (bottom) reference. Samples are colored by dataset provenance with protein activity decreasing cohort-specific batch effects. **b.** PCA of the 122 DMG bulk samples by protein activity using the internal (top) and external (bottom) protein activity signatures with samples colored by cluster assignment based on unsupervised cluster analysis of the internal protein activity signatures demonstrates that cluster stratification is independent of dataset provenance depicted in a. **c.** Unsupervised clustering of DMG patient gene expression signatures versus protein activity signatures with cluster-specific average silhouette width scores depicted below each heatmap showing poor clustering strength in the context of gene expression as compared to protein activity. Heatmap reflects differential protein activity (internal reference) in each case and samples are annotated by relevant clinical and molecular features in the bars on top. **d.** DMG patient overall survival by gene expression-based cluster for 106 patients with outcome information demonstrates no survival stratification by gene expression-based analysis.

**Extended Data Figure 2.**
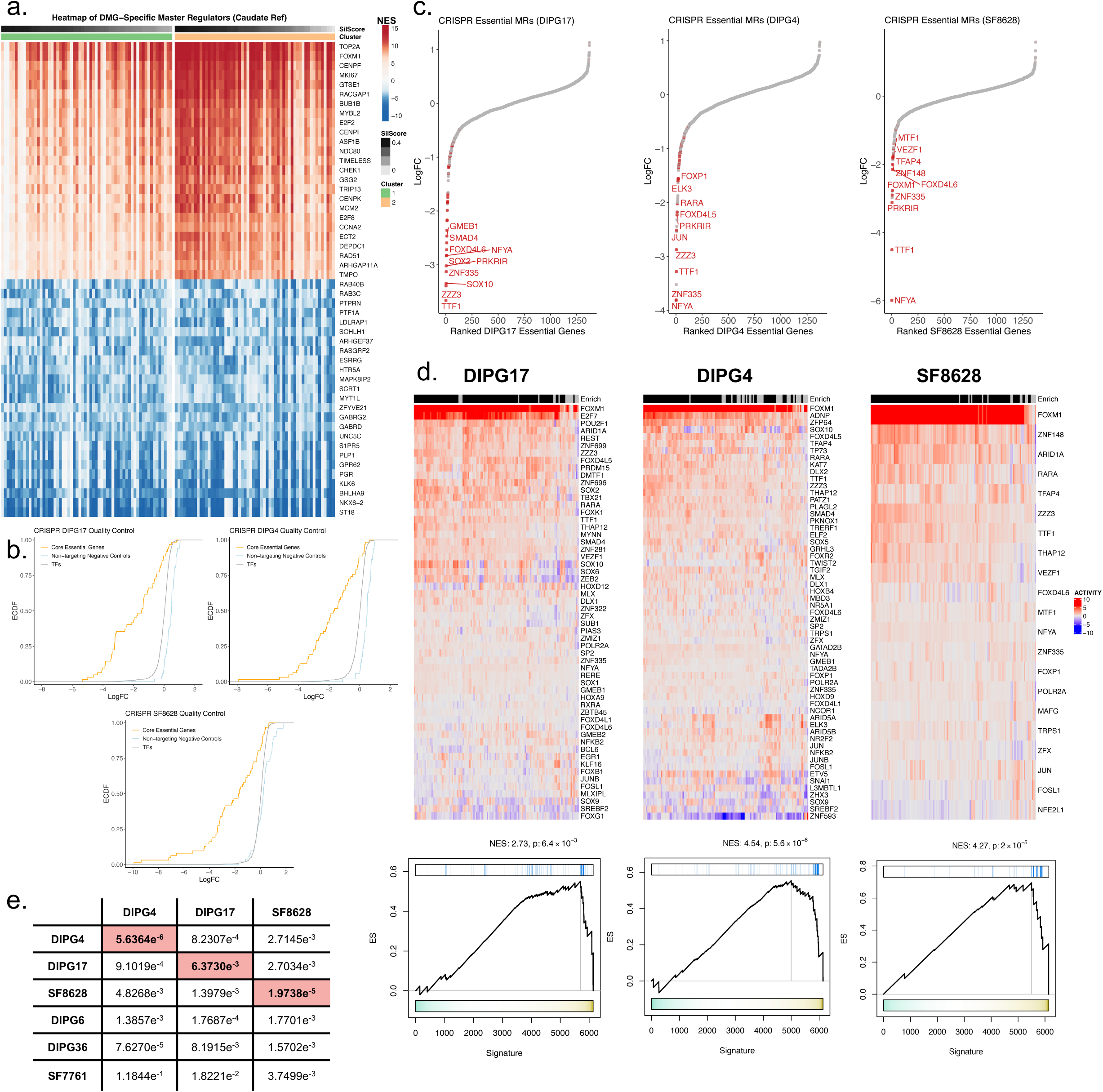
Biological validation of predicted DMG MRs through a pooled CRISPR/Cas9 knockout (KO) screen. **a.** Heatmap of the top and bottom 25 MR proteins based on Stouffer-integrated VIPER-inferred protein activity normalized enrichment scores (NES) across DMG patient samples. Patient samples in columns are divided into their respective unsupervised clusters as in Fig. 1a, demonstrating a common tumorigenic signature across patients. **b.** Empirical Cumulative Distribution (ECDF) of the logFC between the two timepoints (t0 and t1) for each TF evaluated in the CRISPR screen. Core essential positive controls (orange) and non-targeting negative controls (blue) have an overall more negative and positive logFC distribution, respectively. **c.** Scatter plot showing the CRISPR DMG gene essentiality signature for each of the three DMG cell lines tested with genes ranked by their log2FC essentiality score with the most essential genes on the left. VIPER-inferred DMG candidate MRs (Stouffer integrated p-value ≤ 10^-5^ across all patients using the GTEx caudate signature) are shown in red and are enriched among the most essential genes in each cell line. **d.** Heatmap of VIPER-inferred differential protein activity (caudate signature) for the CRISPR essential MRs in each cell line independently. The enrich annotation bar reflects presence (black) or absence (gray) of statistical significance (p ≤ 0.05, one-tailed GSEA) for enrichment of essential genes (negative log2FC and FDR ≤ 0.05) in that cell line in the sample’s differential protein activity signature. The genes displayed for each cell line are those that belong to the leading edge of at least one DMG sample from this analysis. Underneath are results of GSEA analysis demonstrating significant enrichment of essential genes (blue ticks) in each cell line in the ranked Stouffer-integrated protein activity signature of the samples showing statistically significant enrichment from d. **e.** GSEA computed p-values for enrichment of each cell line’s essential genes as assessed by CRISPR in the VIPER-inferred protein activity signature (caudate ref) of six DMG cell lines profiled by bulk RNA-seq. For two of three cell lines, DIPG4 and SF8628, the optimal result was achieved when assessing enrichment of essential genes in genes differentially active in the same cell line, confirming VIPER’s ability to nominate sample-specific MRs.

**Extended Data Figure 3.**
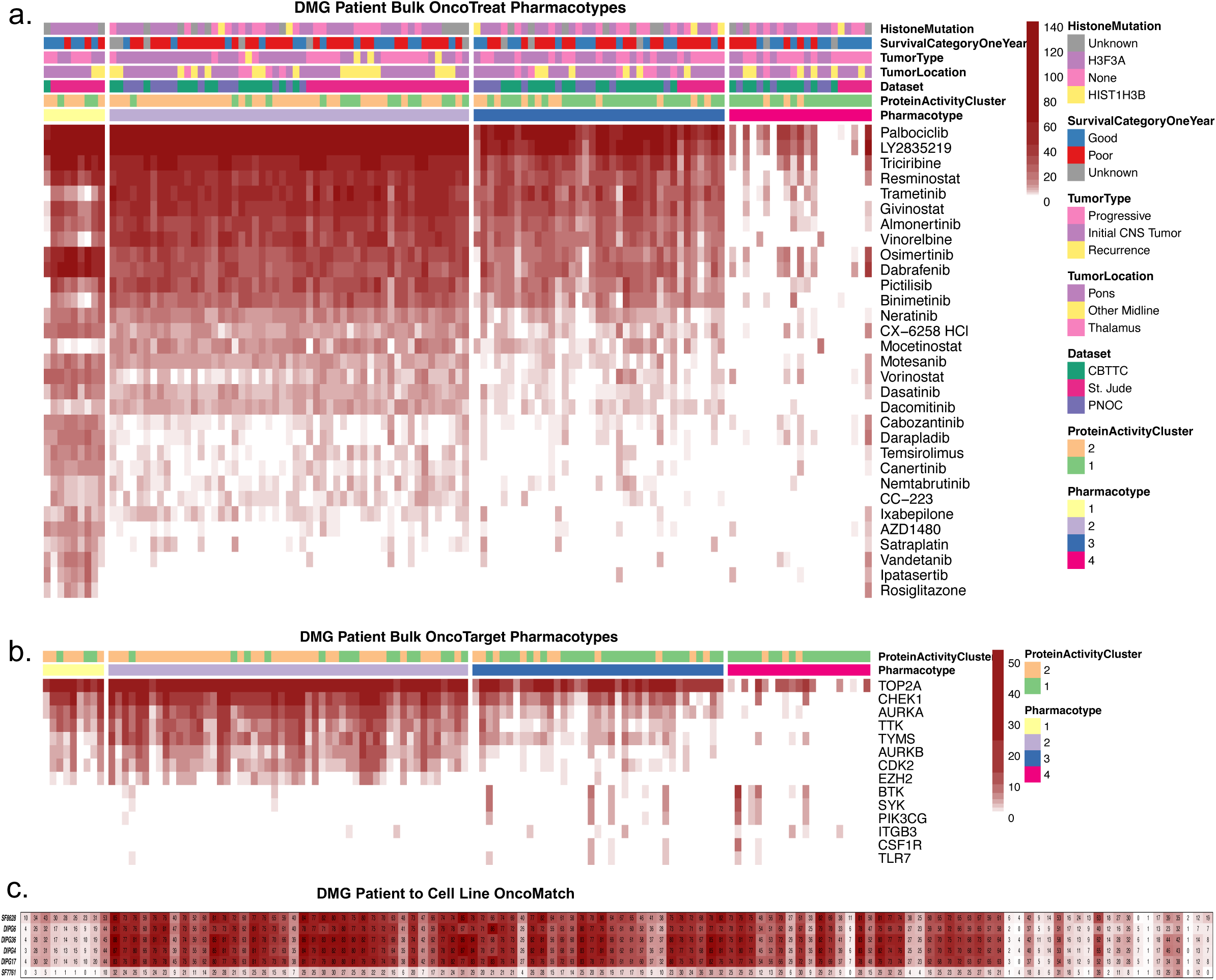
Prediction of MR-inverter drugs by OncoTarget and OncoTreat from bulk RNA-seq samples. **a.** Heatmap showing the top OncoTreat predictions (-log10(Bonferroni p-value)) across 122 DMG bulk samples with unsupervised cluster analysis identifying four main pharmacotypes that significantly stratified with bulk protein activity clusters defined in Fig. 1a. **b.** Heatmap showing the protein activity of actionable proteins computed by OncoTarget with samples ordered as in a. OncoTarget proteins that were significantly active in at least one sample at p ≤ 10^-5^ were included. **c.** OncoMatch of 6 human DMG cell lines (rows) to each of the 122 DMG patients (columns). Values represent -log10(FDR-adjusted p-values) for enrichment of the patient MRs in the cell line protein activity signature.

**Extended Data Figure 4.**
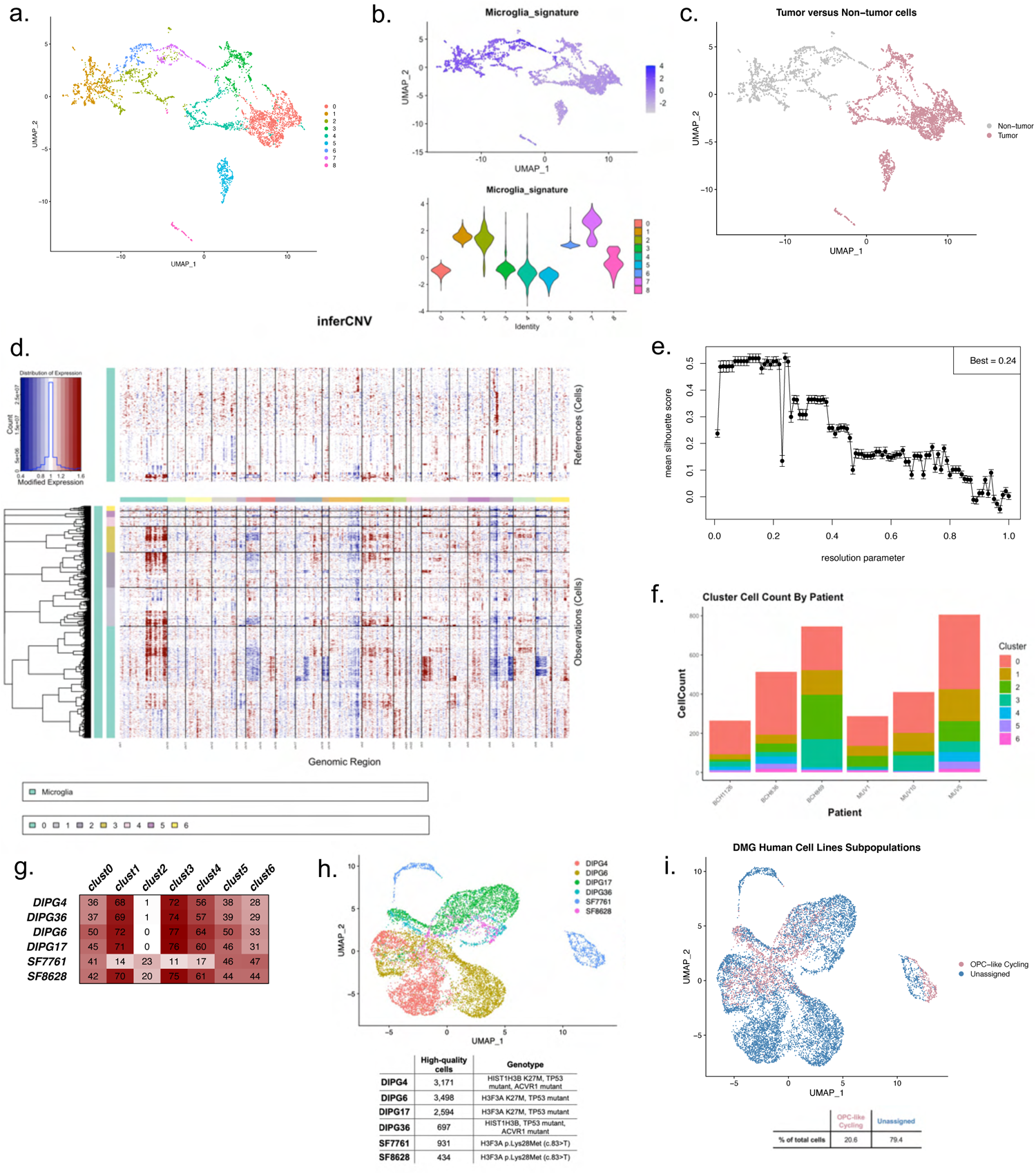
Single cell RNA-seq analysis of DMG patient tumors and cell line models. **a.** UMAP projection of 4,860 single cells profiled by scRNA-seq from 6 DMG patient tumor biopsies using VIPER-inferred protein activity (internal reference) with clusters assigned by resolution-optimized Louvain clustering. **b.** Cells in the UMAP from a. are colored based on GSEA enrichment score for enrichment of a microglia gene signature in the cell’s differential protein activity signature. This, along with the respective violin plots below depicting these enrichment scores across the cells belonging to each unsupervised cluster, were used to identify clusters 1, 2, 6, and 7 as non-tumor immune cells depicted in the UMAP in **c. d.** inferCNV analysis of the presumed tumor cells using cells belonging to the microglia clusters identified in a-c as a reference, demonstrating cells with aberrant copy number changes in each of the remaining clusters which were presumed to be candidate tumor cells. **e.** Resolution-optimized Louvain clustering Silhouette Scores for unsupervised clustering of DMG tumor cells by protein activity. Mean and Standard Deviation of Silhouette scores for each resolution value ranging along the x axis from 0 to 1 at intervals of 0.01 are depicted. The best resolution (mean silhouette score of 0.52) was the one chosen as the final resolution parameter for the clustering of the single cells in the 6 DMG patients. **f.** Cluster cell count (y axis) across the seven identified tumor subpopulations by patient indicates that cells from virtually all clusters co-exist in each of the six patients (x axis). **g.** OncoMatch of 6 human DMG cell lines (rows) to each DMG single cell subpopulation (columns). Values represent - log10(fdr-adjusted p-values) for enrichment of the subpopulation MRs in the cell line protein activity signature. **h.** UMAP of VIPER-inferred protein activity across 11,325 single cells profiled by scRNA-seq across the 6 DMG patient cell lines described in the table below. **i.** UMAP from h. with each cell annotated by the DMG patient single cell subpopulation whose MRs are most significantly enriched in that cell’s differential protein activity signature by GSEA. Unassigned cells reflect those whose GSEA FDR corrected p-value > 0.05 for all populations.

**Extended Data Figure 5.**
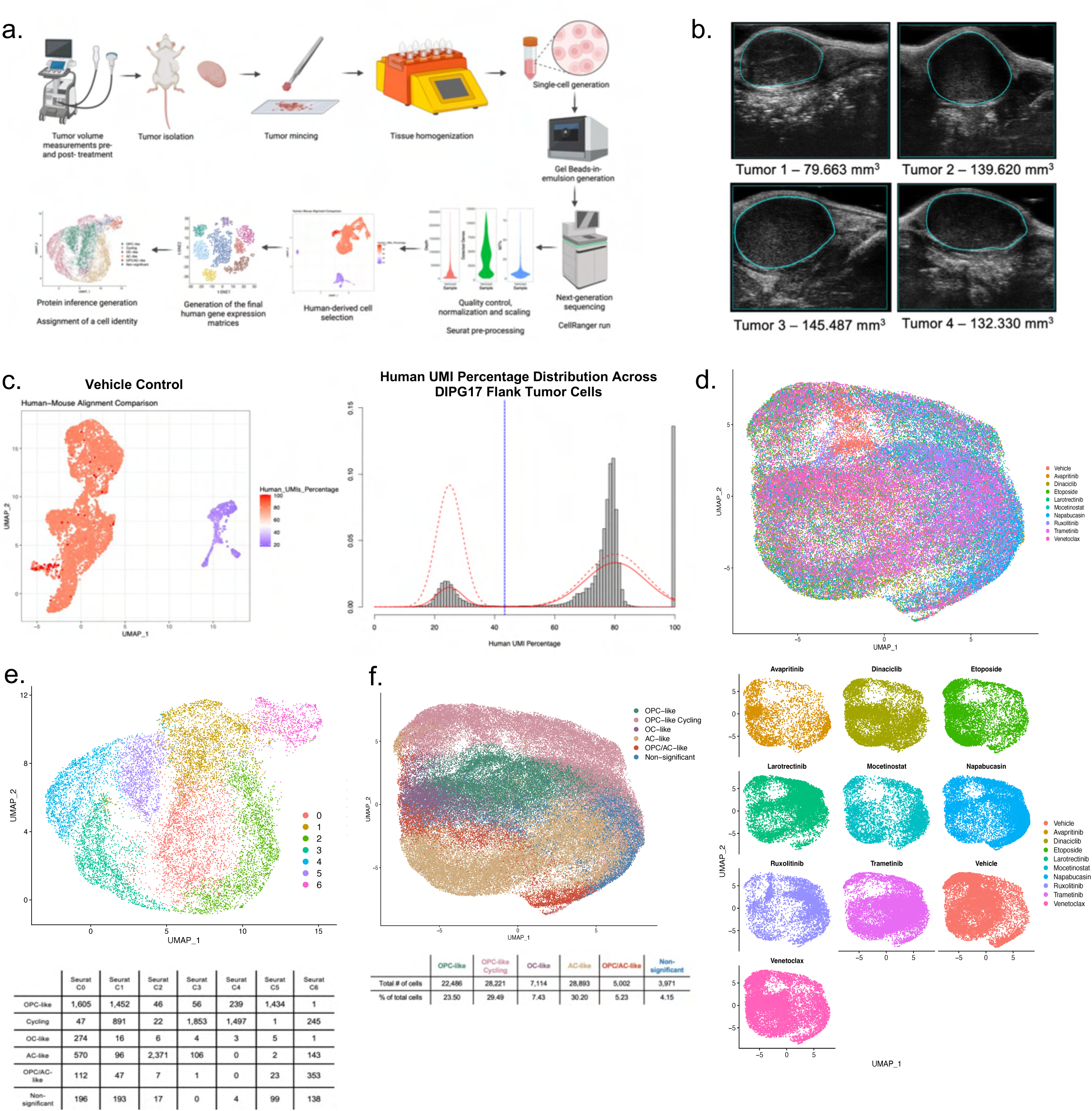
Drug vs. vehicle-treated single-cell RNA-seq in subcutaneous DMG patient cell line (DIPG17)-derived mouse tumors for nine drugs predicted by single-cell OncoTarget/OncoTreat. **a.** Experimental and analytical workflow. **b.** Ultrasound images of four example flank tumors pre-treatment after reaching adequate tumor volume. **c.** Example heat plot showing the proportion of reads from a scRNA-seq profile of a vehicle control-treated tumor that mapped to the human genome reference demonstrating clear separation of likely human, and thus tumor, cells (red) vs. mouse, and thus microenvironment, cells (blue). Distribution of Human UMI percentages across all profiled single cells in this experiment is depicted in the histogram to the right. A fitted finite mixture model was computed (red distribution with dashed line reflecting 95% CI) and used to identify an optimal cutoff value of Human UMI percentage to segregate tumor vs. non-tumor cells (blue line). **d.** UMAP projection of VIPER-inferred protein activity across the 95,687 single cells profiled by scRNA-seq across 9 distinct drug arms and vehicle control (24 total samples) colored by treatment condition and split by treatment arm below showing conserved subpopulation structure across treatment conditions and no sample-specific batch effects. **e.** UMAP depicting resolution optimized Louvain clustering of protein activity profiles from 14,176 single cells across four vehicle control-treated tumors. Table below shows the number of cells in each cluster annotated as each of the five DMG patient tumor cell states based on the most significant enrichment of each cell states’ MRs in the cell’s protein activity signature computed by GSEA (cells with GSEA adjusted p-value > 0.05 across all cell states were labeled as Non-significant). We see segregation of DMG patient tumor cell states with the unsupervised tumor model clusters supporting the fidelity of this model at recapitulating patient tumor heterogeneity. **f.** UMAP from d. with cells annotated by their most significantly matched DMG patient tumor cell state.

**Extended Data Figure 6.**
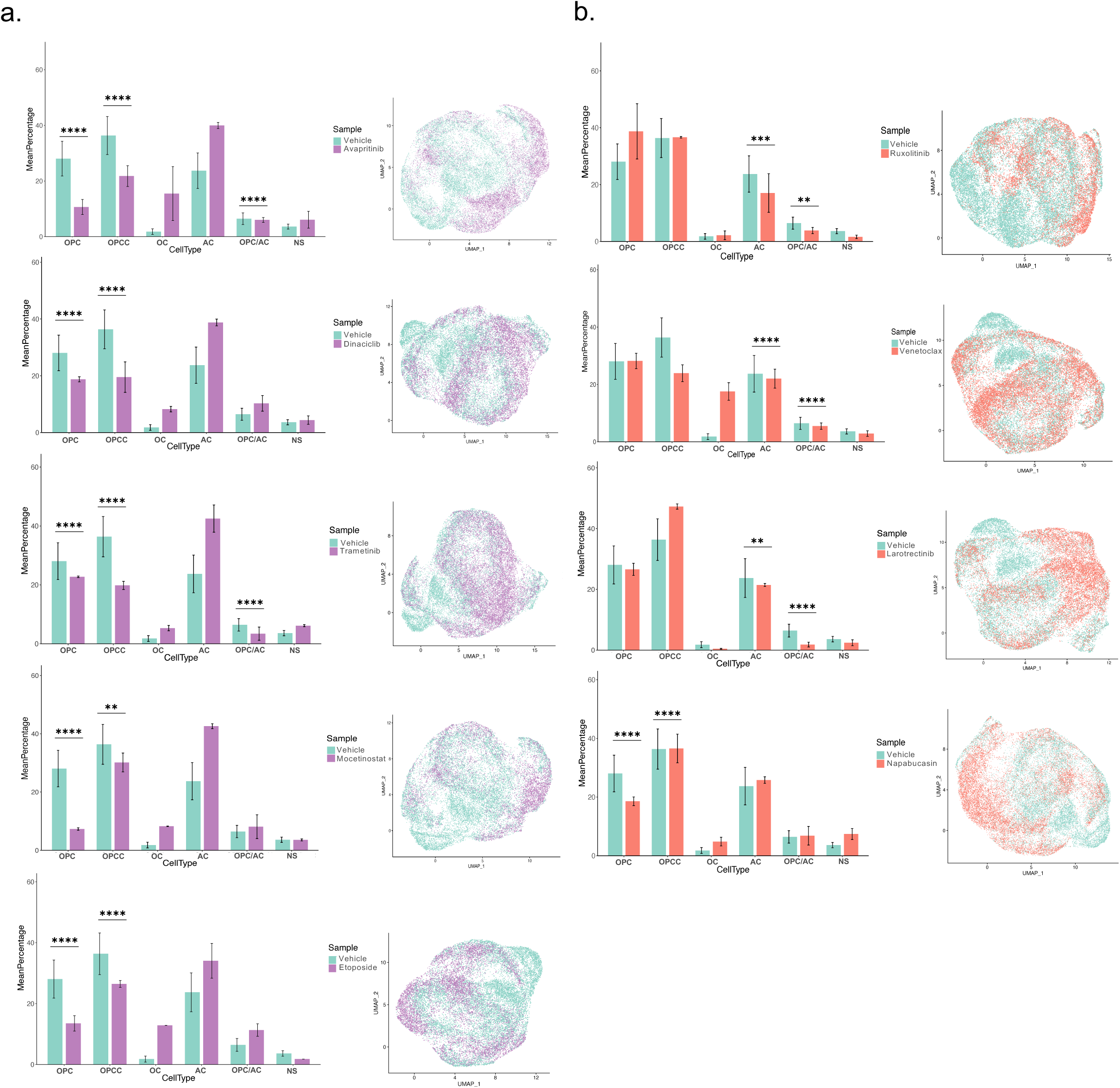
Subpopulation changes for nine drugs predicted by single-cell OncoTarget/OncoTreat and their respective vehicles in subcutaneous DMG patient cell line (DIPG17)-derived mouse tumors. **a.** Subpopulation comparison bar plots (depicting mean percentage) for all 5 drugs targeting the OPC-like cell states (purple) compared to vehicles (green) (with Standard Error (SE)) (left) and UMAP projections of the 14,176 vehicle-treated and 38,615 cells treated with agents targeting OPC-like cell states (right) for (from top to bottom) avapritinib, dinaciclib, trametinib, mocetinostat and etoposide. **b.** Subpopulation comparison bar plots (depicting mean percentage) for all 4 drugs targeting the AC-like states (orange) compared to vehicles (green) (with Standard Error (SE)) (left) and UMAP projections of the 14,176 vehicle-treated and 42,896 cells treated with agents targeting AC-like cell states (right) for (from top to bottom) ruxolitinib, venetoclax, larotrectinib and napabucasin. ✱ denote chi-square p-value significance, where ✱p<0.05, ✱✱p<0.01, ✱✱✱p<0.001, ✱✱✱✱p<0.0001. Error bars indicate +/- Standard Error (SE).

**Extended Data Figure 7.**
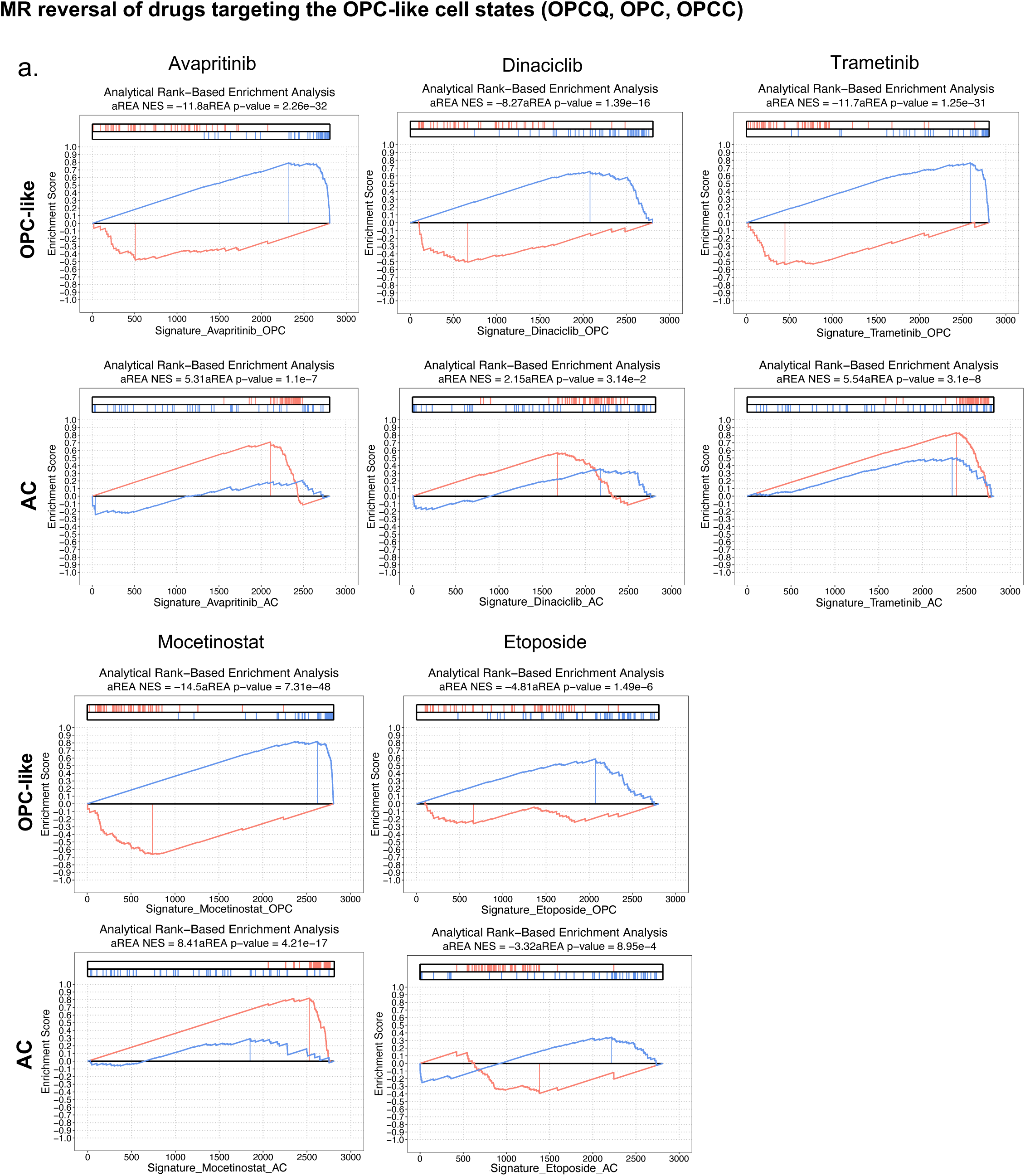
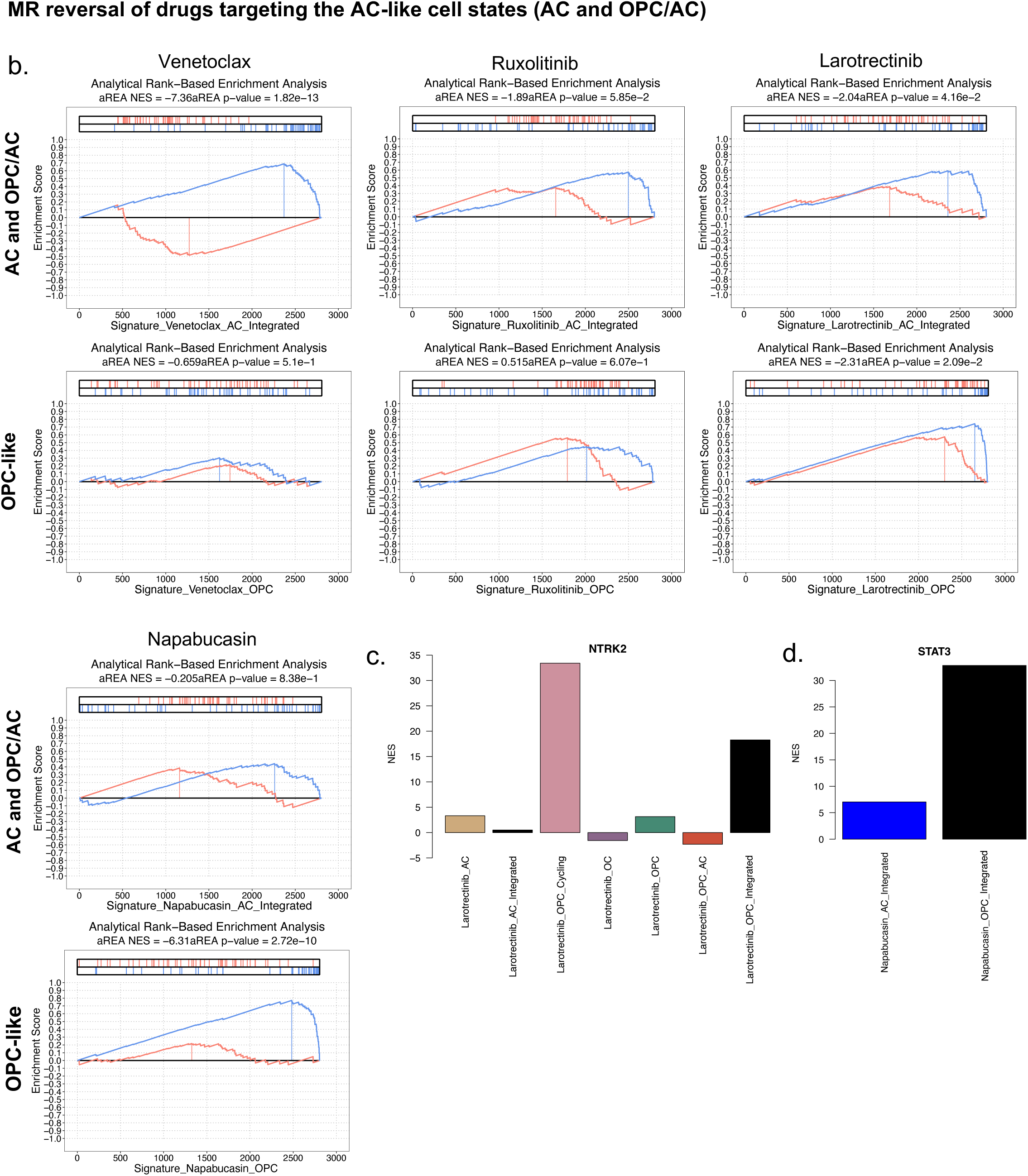
Pharmacodynamic assessment of tumor checkpoint module (TCM) inversion after five days of treatment in subcutaneous DMG patient cell line (DIPG17)-derived mouse tumors. **a.** Analytical Rank-based Enrichment Analysis (aREA) of the top (red ticks) and bottom (blue ticks) 50 DMG patient population-specific MRs in the differential protein activity signature of drug- vs. vehicle-treated single cells within that population in the DIPG17 mouse tumors for all 5 drugs targeting the OPC-like cell states. Avapritinib, dinaciclib, trametinib, mocetinostat and etoposide all significantly invert the TCM in OPC-like cells (negative aREA NES) but not the AC-like state (positive aREA NES). **b.** Drugs targeting the AC-like states (AC and OPC/AC), conversely, invert the TCM in AC-like cells (negative aREA NES) but not the OPC-like state. **c.** Differential protein activity of NTRK2 in drug- vs. vehicle-treated cells within the AC- and OPC-like populations, confirms direct targeting of NTRK2 activity in OC- and OPC/AC-like cells by larotrectinib and **d.** suggests napabucasin targeting of cell states is not due to direct targeting of STAT3 activity.

**Extended Data Figure 8.**
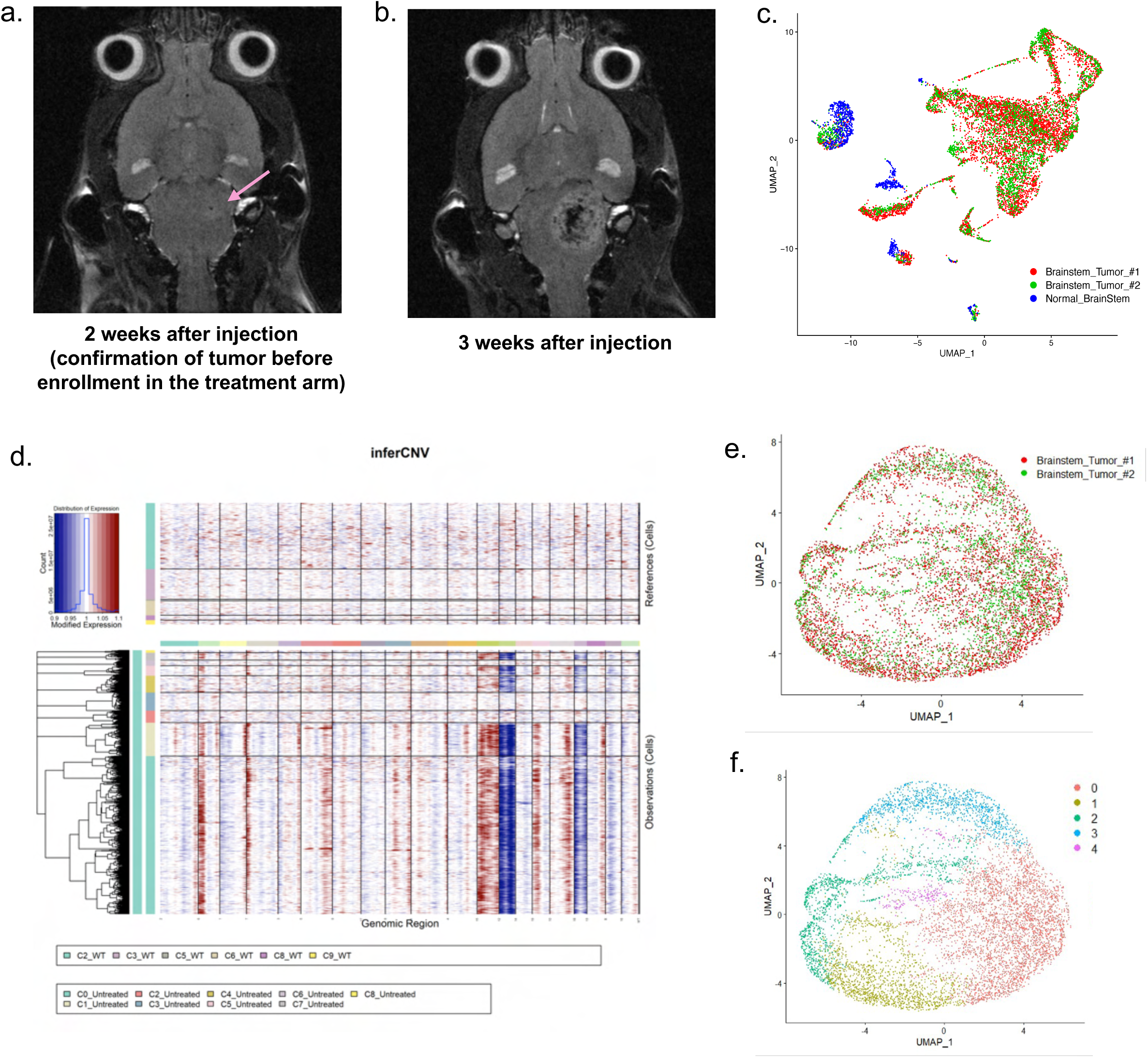
Generation of a syngeneic orthotopic DMG murine models and validation of model fidelity and heterogeneity by single cell RNA-seq. **a.** MRI images of the brainstem tumor in the DIPG4423 syngeneic mouse model at **a.** 2 weeks after injection before enrollment in the treatment arm (pink arrow indicating correct injection site). Enhancement around the area of injection indicates tumor presence and **b.** 3 weeks after treatment with vehicle control. **c.** UMAP projection of after the removal of the immune system-related clusters showing all three tumor samples (two brainstem tumors and one control) **d.** InferCNV analysis of brainstem tumor samples (untreated) using the control brainstem (WT) as a reference. InferCNV reveals a total of 8,013 malignant cells in clusters 0, 1, 4, 5 and 7 indicated by the high number of copy number alterations in these clusters. Clusters not included in the reference or output did not have enough cells to be included in the analysis. **c.** UMAP depicting protein activity inference of the 2 DMG samples (8,013 tumor cells) and **d.** UMAP from c. resolution optimized Louvain clustering of protein activity profiles.

**Extended Data Figure 9.**
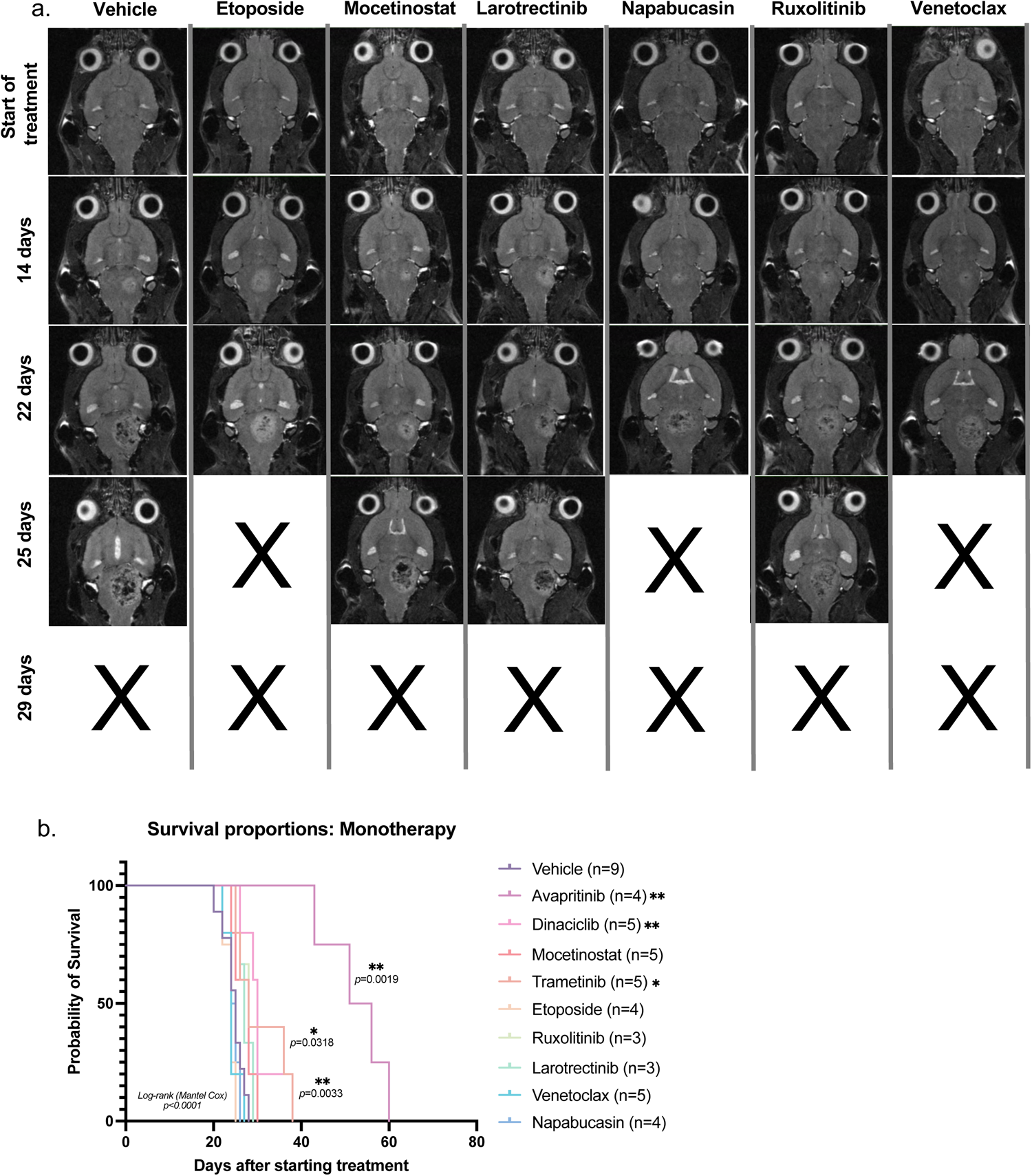
MRI showing tumor progression in syngeneic orthotopic DMG murine models for all monotherapy drug arms with no significant survival benefit. **a.** MRIs of monotherapy treatments with no significant survival differences compared to vehicle controls. Every image depicts one mouse at the indicated time (rows) for a particular treatment (column). X denotes mice have succumbed to the tumor. **b.** Survival analysis (Kaplan-Meier) of all monotherapy drug arms with significant differences in survival with respect to vehicle controls assessed by log-rank test. Avapritinib, dinaciclib and trametinib showed significant survival compared to vehicle controls, as denoted by stars. All other treatments were not significant. ✱ denote p-value significance, where ✱p<0.05, ✱✱p<0.01, ✱✱✱p<0.001, ✱✱✱✱p<0.0001

**Extended Data Figure 10.**
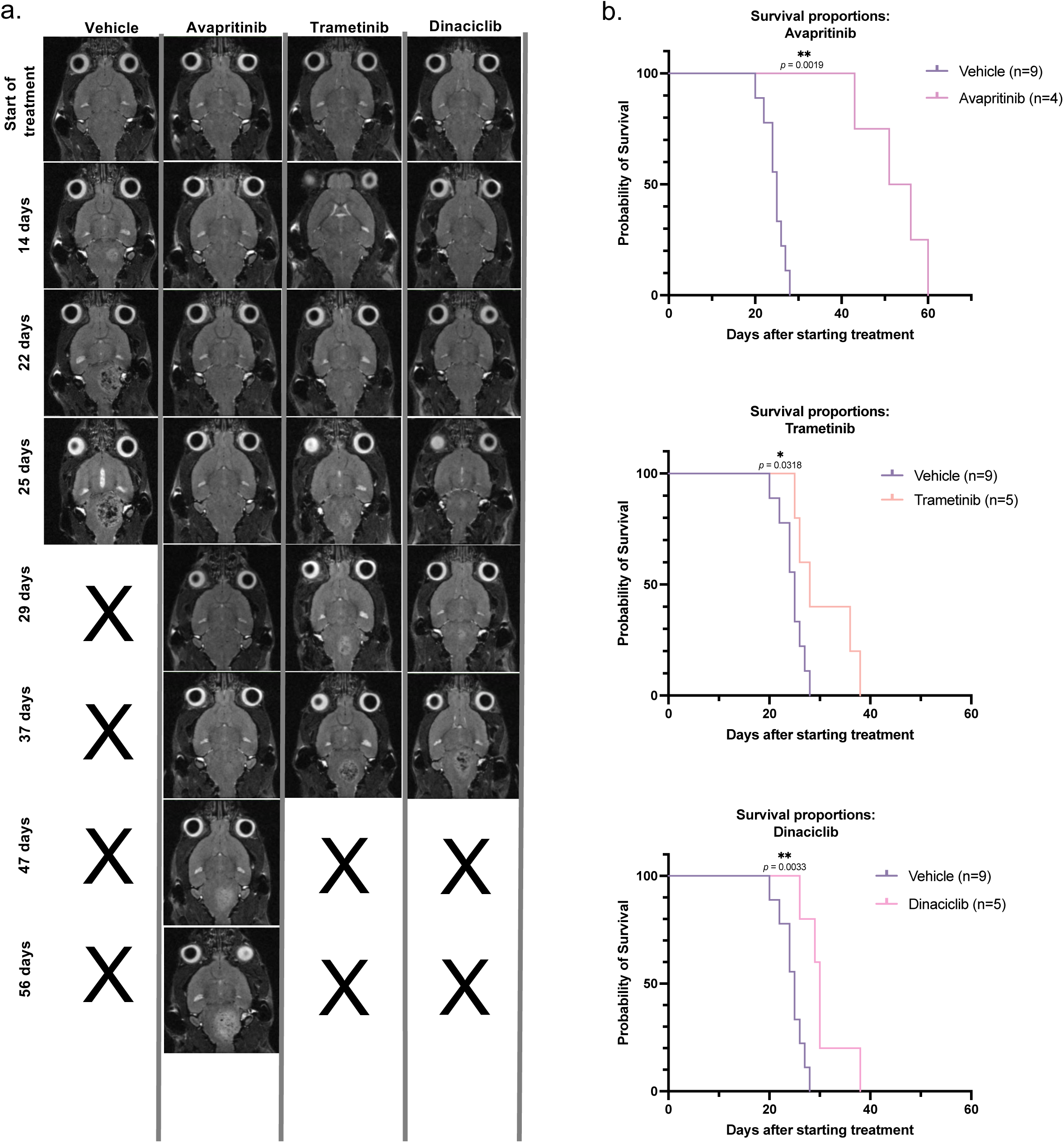
MRI showing tumor progression in syngeneic orthotopic DMG murine models for all monotherapy drug arms with significant survival benefit. **a.** MRIs of monotherapy treatments (avapritinib, trametinib and dinaciclib) with significant survival differences compared to vehicle controls. Every image depicts one mouse at the indicated time (rows) for a particular treatment (column). X denotes mice have succumbed to the tumor. **b.** Survival analysis (Kaplan-Meier) of all significant monotherapy arms separately (avapritinib, trametinib and dinaciclib, top to bottom) compared to vehicle controls. ✱ denote p-value significance, where ✱p<0.05, ✱✱p<0.01, ✱✱✱p<0.001, ✱✱✱✱p<0.0001.

**Extended Data Figure 11.**
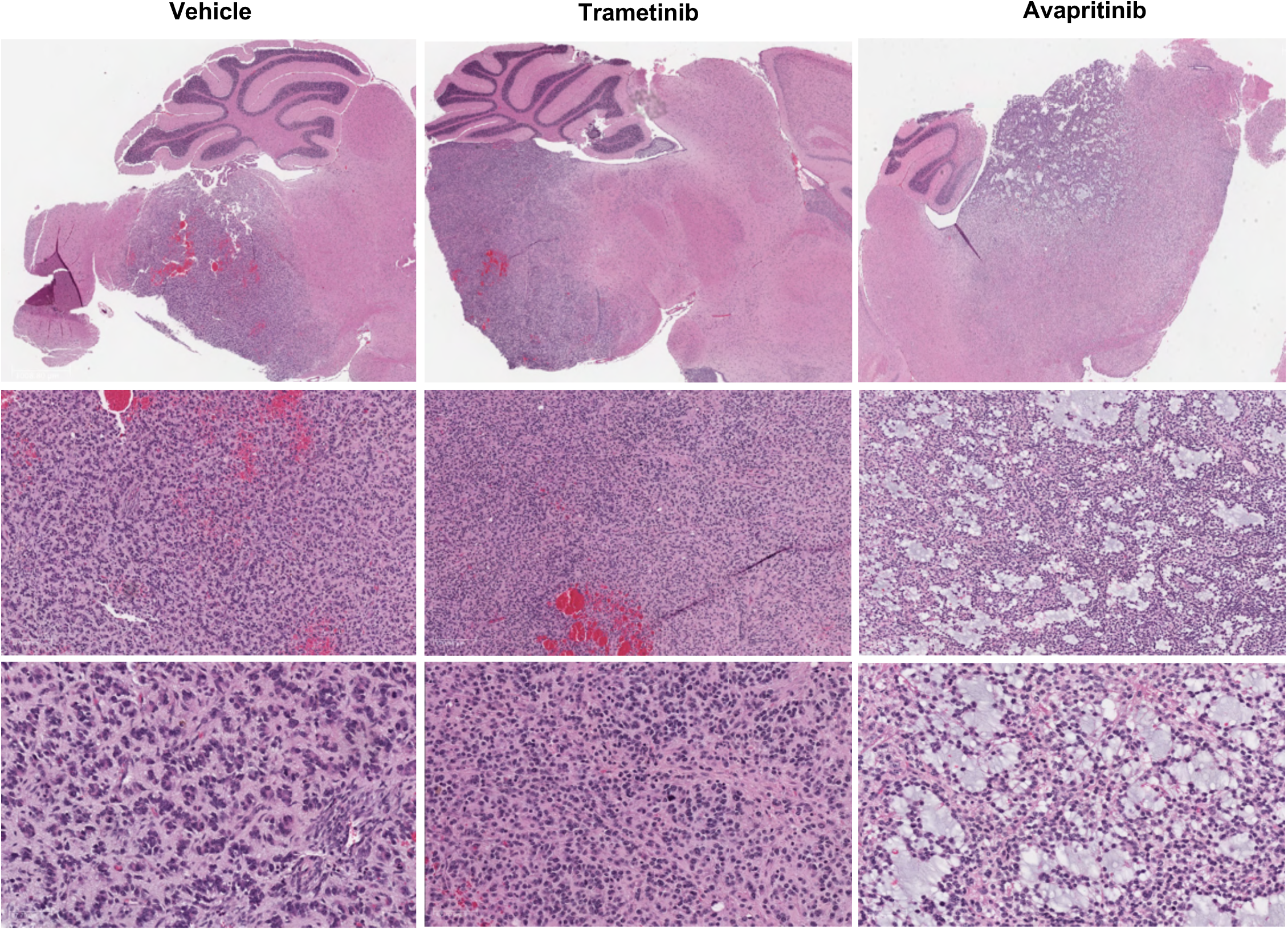
Treatment with avapritinib results in morphologically distinct tumors. Low (1008.80μm, top images) and high powered (170.98μm, middle and 67.70μm bottom) images showing H&E stains of end stage tumors in vehicle, trametinib, and avapritinib treated. Avapritinib treated tumors (right) show altered morphology represented by diffuse infiltration and a notable absence of necrosis not present in the vehicle controls or other treatment arms (trametinib).

**Extended Data Figure 12.**
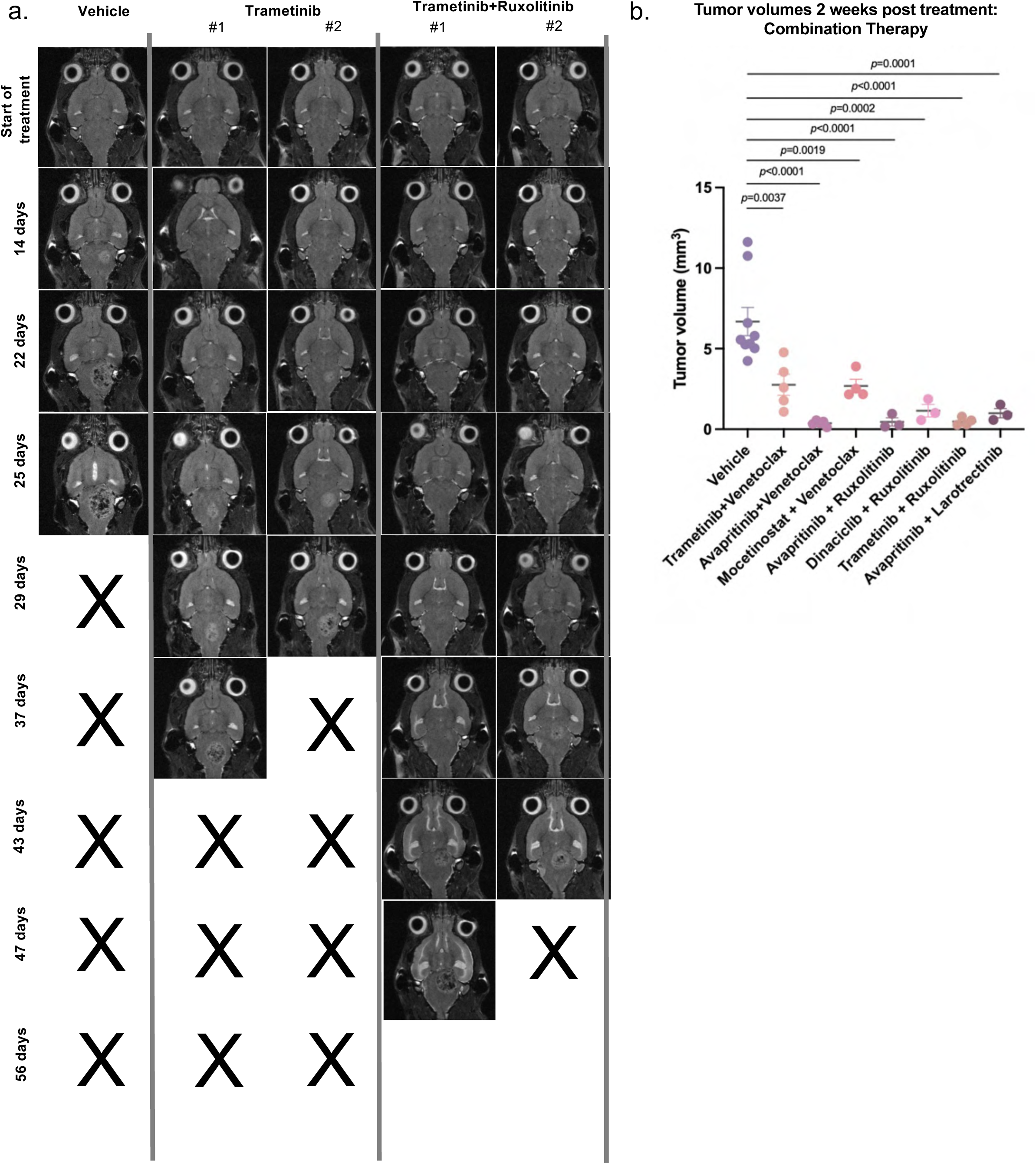
MRI showing tumor progression in syngeneic orthotopic DMG murine models for trametinib drug arms with significant survival benefit. **a.** MRI images of trametinib treatment arms (alone and in combination) with significant survival differences compared to vehicle controls for monotherapy, and compared to monotherapy for combination therapy (trametinib+ruxolitinib). Every image depicts one mouse (two mice shown for each trametinib treatment arm) at the indicated time (rows) for a particular treatment (column). X denotes mice have succumbed to the tumor. **b.** Tumor volume changes two weeks after treatment with combination therapy for all drug arms compared to vehicle controls.

**Extended Data Figure 13.**
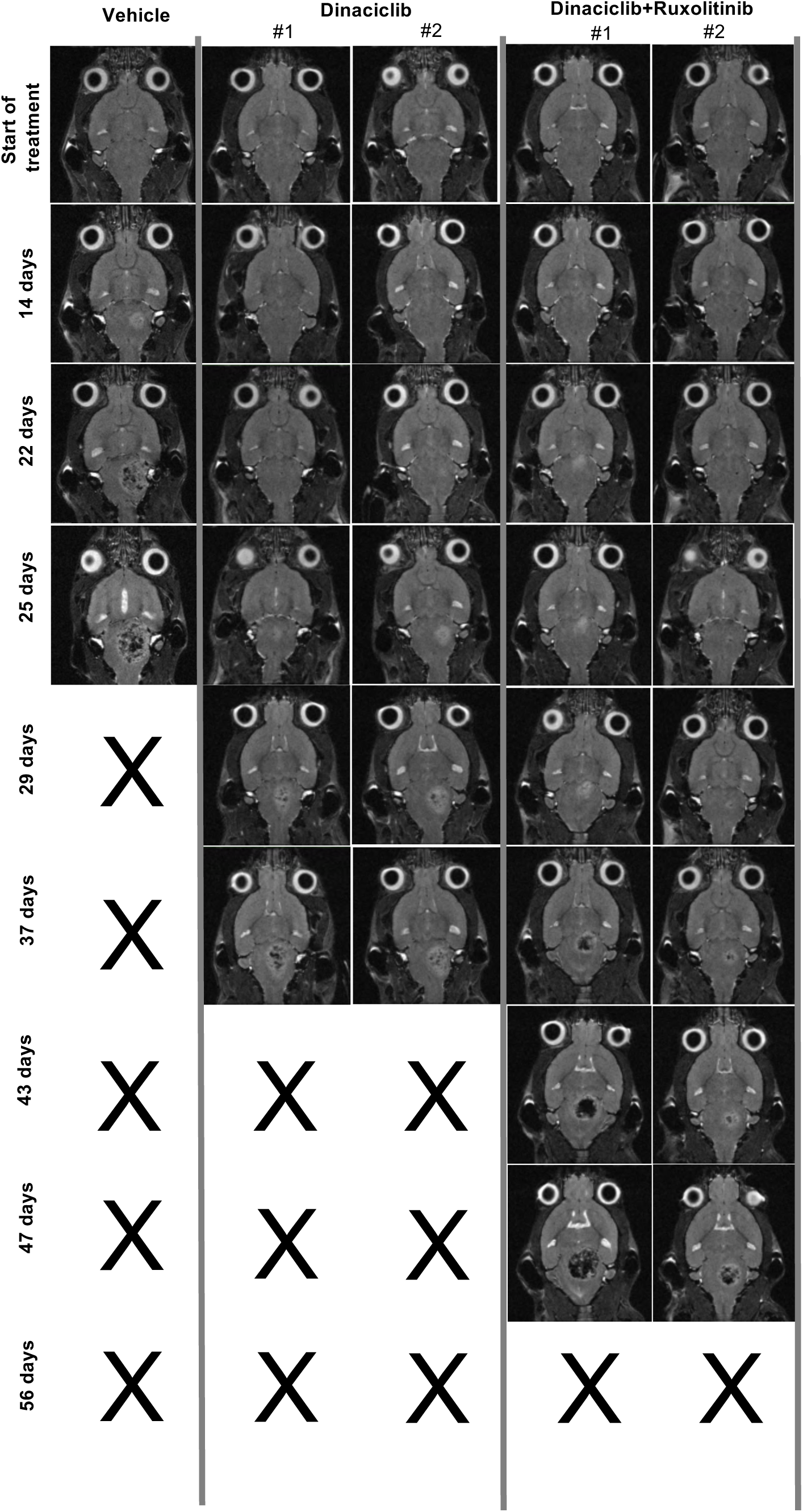
MRI showing tumor progression in syngeneic orthotopic DMG murine models for dinaciclib drug arms with significant survival benefit. **a.** MRI images of dinaciclib treatment arms (alone and in combination) with significant survival differences compared to vehicle controls for monotherapy and compared to monotherapy for combination therapy (dinaciclib+ruxolitinib). Every image depicts one mouse (two mice shown for each dinaciclib treatment arm) at the indicated time (rows) for a particular treatment (column). X denotes mice have succumbed to the tumor.

**Extended Data Figure 14.**
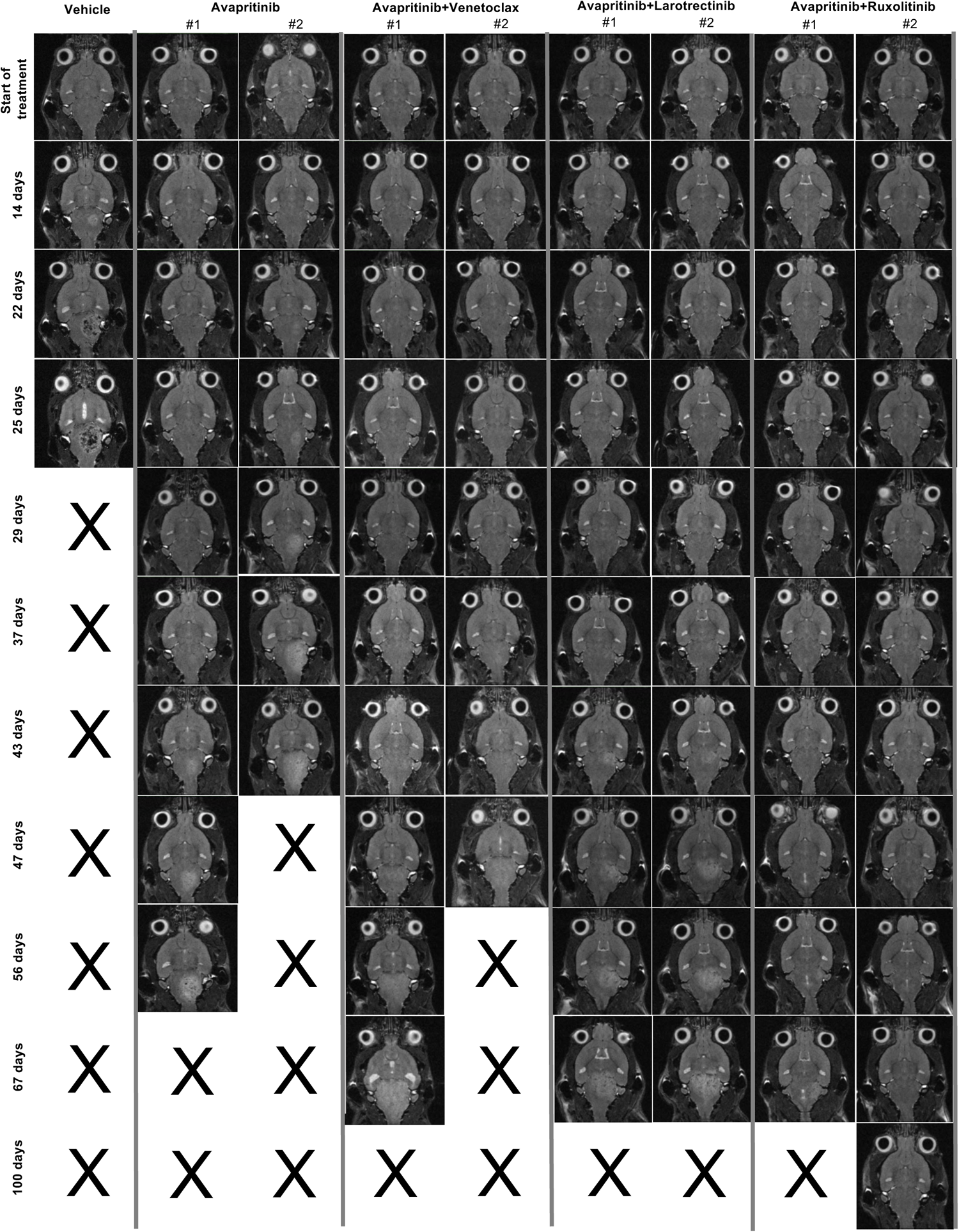
MRI showing tumor progression in syngeneic orthotopic DMG murine models for all avapritinib drug arms. **a.** MRI images of avapritinib treatment arms alone and in combination (avapritinib+venetoclax, avapritinib+larotrectinib, avapritinib+ruxolitinib). Every image depicts one mouse (two mice shown for each avapritinib treatment arm) at the indicated time (rows) for a particular treatment (column). X denotes mice have succumbed to the tumor.

**Extended Data Figure 15.**
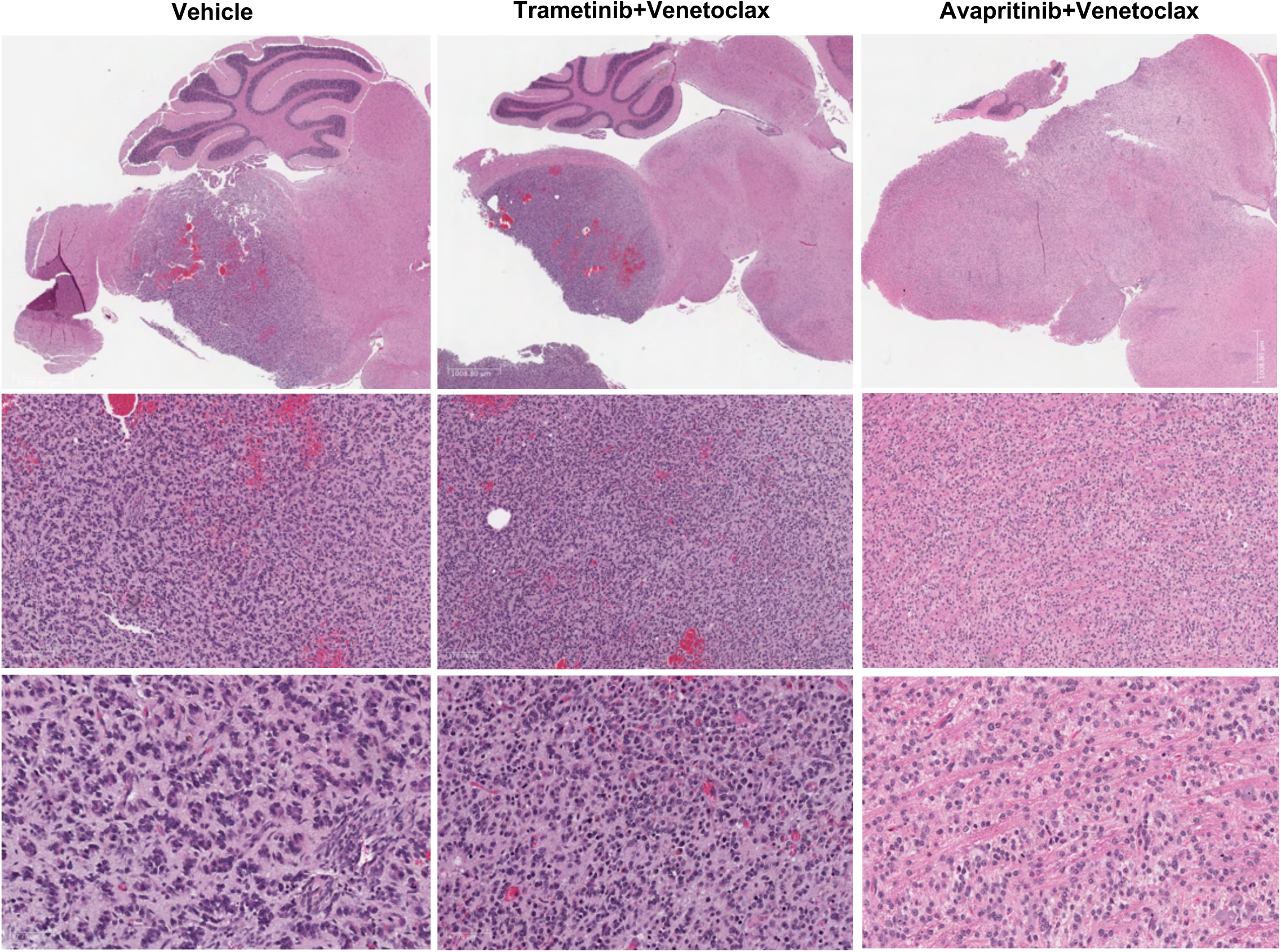
Treatment with avapritinib in combination with drugs targeting AC-like cell states results in morphologically distinct tumors. Low (1008.80μm, top images) and high powered (170.98μm, middle and 67.70μm bottom) images showing H&E stains of end stage tumors in vehicle, trametinib+venetoclax, and avapritinib+venetoclax treated. Avapritinib treated tumors (right) show altered morphology represented by diffuse infiltration and a notable absence of necrosis not present in the vehicle controls or other combination therapy treatment arms (trametinib+venetoclax).

**Extended Data Figure 16.**
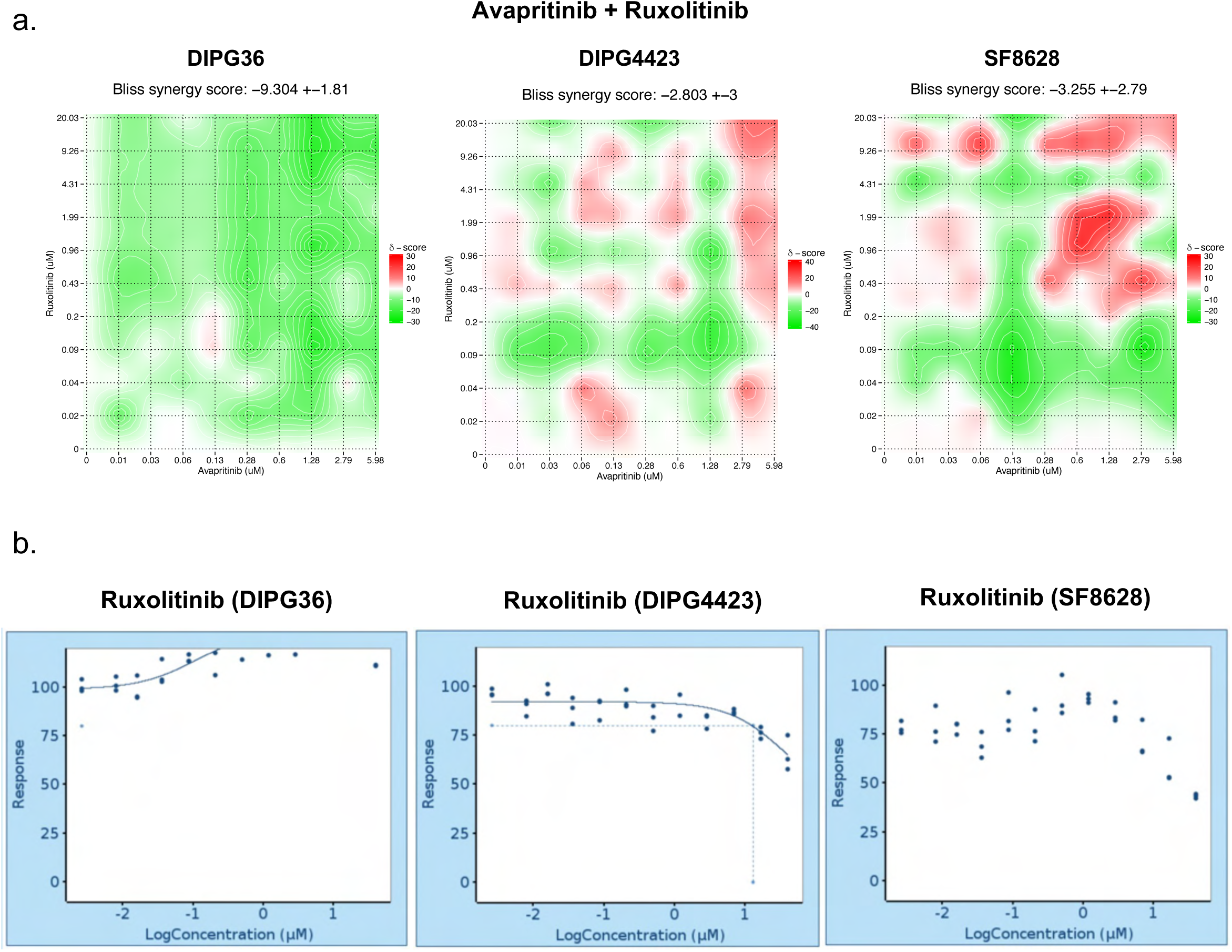
BLISS independence studies confirm that effects of combination therapies targeting distinct cell states are not due to synergy of effects in the predominant tumor subpopulations. **a.** BLISS synergy studies for avapritinib+ruxolitinib in three distinct cell lines: DIPG36 (human), DIPG4423 (murine), and SF8628 (human) depict negative Bliss synergy scores, indicating absence of synergy for this drug combination *in vitro*. **b.** Dose-response curves for ruxolitinib in the 3 different DMG cell lines caused limited cell death in all three cell lines with DIPG36 (left) successfully expanding while on treatment.

